# Chemoenzymatic syntheses of fluorine-18-labeled disaccharides from [^18^F]FDG yield potent sensors of living bacteria *in vivo*

**DOI:** 10.1101/2023.05.20.541529

**Authors:** Alexandre M. Sorlin, Marina López-Álvarez, Sarah J. Rabbitt, Aryn A. Alanizi, Rebecca Shuere, Kondapa Naidu Bobba, Joseph Blecha, Sasank Sakhamuri, Michael J. Evans, Kenneth W. Bayles, Robert R. Flavell, Oren S. Rosenberg, Renuka Sriram, Tom Desmet, Bernd Nidetzky, Joanne Engel, Michael A. Ohliger, James S. Fraser, David M. Wilson

**Affiliations:** Department of Radiology and Biomedical Imaging, University of California, San Francisco, San Francisco, CA 94158, USA; Department of Pathology and Microbiology, University of Nebraska Medical Center, Omaha, NE 68198, USA; Department of Medicine, University of California, San Francisco, San Francisco, CA 94158, USA; Department of Biotechnology, Ghent University, Gent, Belgium; Institute of Biotechnology and Biochemical Engineering, Graz University of Technology, Graz, Austria; Department of Radiology, Zuckerberg San Francisco General Hospital, San Francisco, CA 94110, USA; Department of Bioengineering and Therapeutic Sciences, University of California, San Francisco, San Francisco, CA 94158, USA

**Keywords:** Infection, imaging, positron emission tomography, chemoenzymatic synthesis

## Abstract

Chemoenzymatic techniques have been applied extensively to pharmaceutical development, most effectively when routine synthetic methods fail. The regioselective and stereoselective construction of structurally complex glycans is an elegant application of this approach, that is seldom applied to positron emission tomography (PET) tracers. We sought a method to dimerize 2-deoxy-[^18^F]-fluoro-D-glucose ([^18^F]FDG), the most common tracer used in clinical imaging, to form [^18^F]-labeled disaccharides for detecting microorganisms *in vivo* based on their bacteria-specific glycan incorporation. When [^18^F]FDG was reacted with β-D-glucose-1-phosphate in the presence of maltose phosphorylase, both the α-1,4 and α-1,3-linked products 2-deoxy-[^18^F]-fluoro-maltose ([^18^F]FDM) and 2-deoxy-2-[^18^F]-fluoro-sakebiose ([^18^F]FSK) were obtained. This method was further extended with the use of trehalose (α,α-1,1), laminaribiose (β-1,3), and cellobiose (β-1,4) phosphorylases to synthesize 2-deoxy-2-[^18^F]fluoro-trehalose ([^18^F]FDT), 2-deoxy-2-[^18^F]fluoro-laminaribiose ([^18^F]FDL), and 2-deoxy-2-[^18^F]fluoro-cellobiose ([^18^F]FDC). We subsequently tested [^18^F]FDM and [^18^F]FSK *in vitro,* showing accumulation by several clinically relevant pathogens including *Staphylococcus aureus* and *Acinetobacter baumannii,* and demonstrated their specific uptake *in vivo.* The lead sakebiose-derived tracer [^18^F]FSK was stable in human serum and showed high uptake in preclinical models of myositis and vertebral discitis-osteomyelitis. Both the synthetic ease, and high sensitivity of [^18^F]FSK to *S. aureus* including methicillin-resistant (MRSA) strains strongly justify clinical translation of this tracer to infected patients. Furthermore, this work suggests that chemoenzymatic radiosyntheses of complex [^18^F]FDG-derived oligomers will afford a wide array of PET radiotracers for infectious and oncologic applications.

---

Biocatalysis is now frequently used in chemical synthesis, both for the development of new building blocks and late-stage modification of complex molecules^1^. Chemoenzymatic syntheses of polysaccharides are particularly appealing, given the challenges of regioselectivity and stereoselectivity using standard organic methods^2, 3^. Glucose-based polysaccharides are amenable to enzymatic incorporation of unnatural monosaccharide units, suggesting the possibility of using the common clinical imaging tracer 2-deoxy-[^18^F]-fluoro-D-glucose ([^18^F]FDG) as a synthon for building complex glycans compatible with *in vivo* positron emission tomography (PET) imaging. Both chemical and enzymatic transformations of [^18^F]FDG represent promising ways to develop new imaging tools, given the wide availability of this tracer. For example, [^18^F]FDG can (1) be reduced with NaBH_4_ to produce the *Enterobacteriaceae*-targeted PET radiotracer 2-deoxy-[^18^F]-fluoro-D-sorbitol ([^18^F]FDS)^4, 5^, (2) be derivatized with amines and other substituents to yield analyte-sensitive “caged” prodrugs^6, 7^ and (3) be employed as a prosthetic group to label drugs and peptide structures^8, 9^. These approaches can be used to easily transform [^18^F]FDG into tracers targeting bacterial metabolism, the tumoral microenvironment, or specific oncologic targets, for example the prostate-specific membrane antigen (PSMA) found in prostate cancer.

Our previous studies describing molecular tools for bacterial infection have focused on PET tracers labeling the bacterial cell wall, including D-[3-^11^C]alanine^10^ and D-[methyl-^11^C]methionine^11–13^, as well as the folic acid pathway using α-[^11^C]PABA^14^. In addition to maltose-derived PET radiotracers, there have been several elegant methods in the last decade applied to bacteria using siderophore-derived probes^15, 16^, radiolabeled trimethoprim derivatives^17^, and [^18^F] sugar alcohols including [^18^F]FDS^4^ and 2-deoxy-[^18^F]-fluoro-D-mannitol^18^. The most advanced radiotracer in bacteria-specific PET imaging is [^18^F]FDS, which has been studied in numerous advanced preclinical models of infection as well as in infected patients^19^. A recent innovation in [^18^F]FDS is its rapid, kit-based, and on-demand radiosynthesis from [^18^F]FDG^20^. One limitation of [^18^F]FDS is lack of sensitivity for gram-positive pathogens, which are major causes of musculoskeletal infections including vertebral discitis-osteomyelitis (VDO)^21, 22^. Therefore, we and others have a sustained interest in developing [^18^F]-labeled PET tracers targeting *S. aureus* and other gram-positive organisms.

In the current study, we used readily available [^18^F]FDG to construct [^18^F]-labeled dimers in one step via chemoenzymatic syntheses. This approach contrasts with most chemical methods reported to generate [^18^F]glycans, which typically use a structurally complex/ protected precursor, and [^18^F]fluoride incorporation via S_N_2 displacement at sterically amenable sites^23^. Based on prior work, a phosphorylase-catalyzed approach appeared feasible^24–28^, with several glucose-derived dimers highlighted in **Figure 1A**. Phosphorylases (glycosyltransferases, E.C. 2.4) are enzymes that catalyze the addition of an inorganic phosphate group to an carbohydrate acceptor with breaking an *O*-glycosidic bond. This reaction can be run “in reverse” to construct disaccharides from D-glucose-1-phosphate and a glucose derivative (**Figure 1B**). This strategy might therefore be used to rapidly fabricate [^18^F]-labeled disaccharides from [^18^F]FDG.

**Figure 1.**
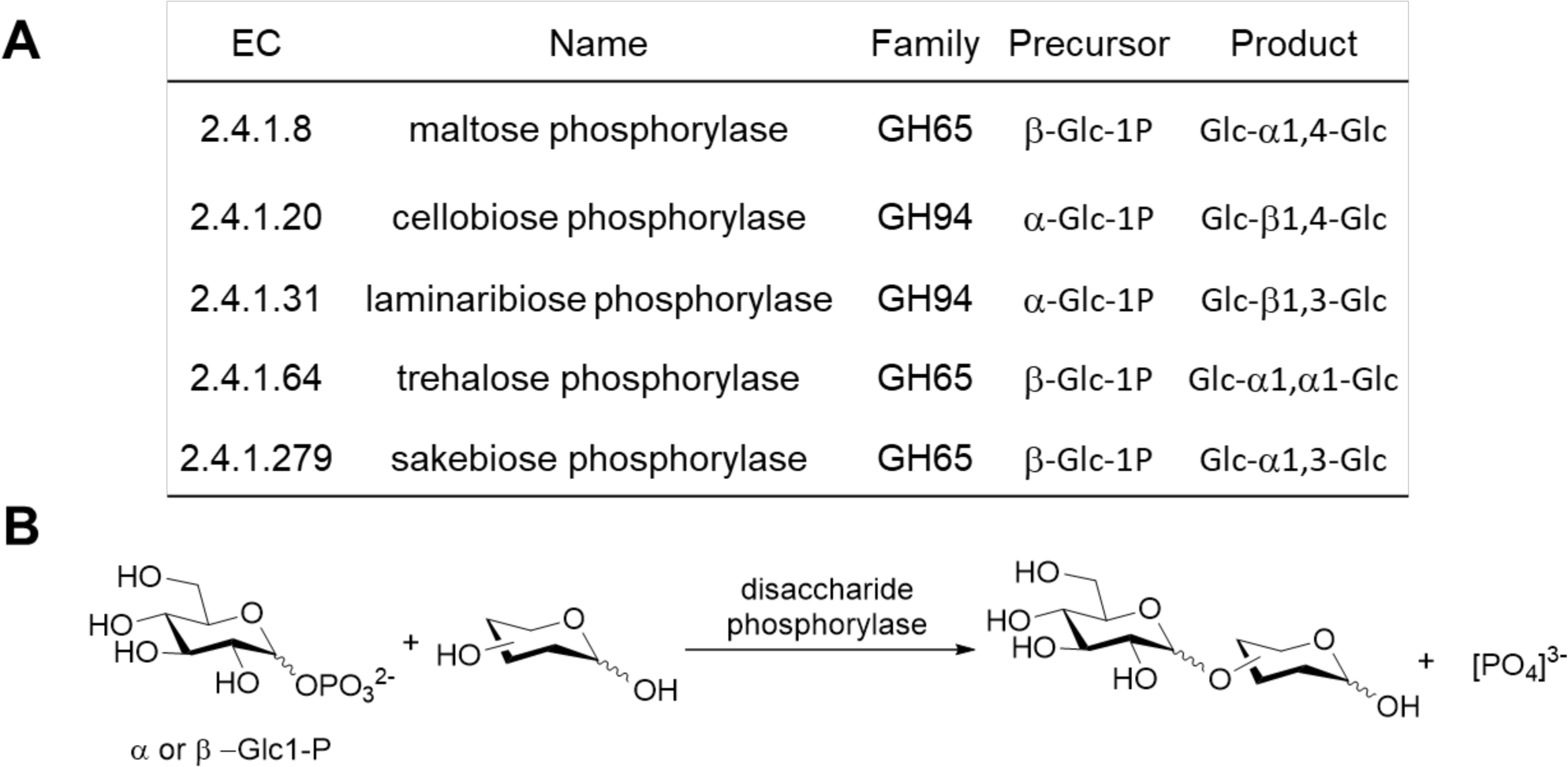
(A) Partial list of disaccharide phosphorylases with the potential for chemoenzymatic synthesis of disaccharides via reverse phosphorolysis. (B) Reverse phosphorolysis of substrates using disaccharide phosphorylase with either α- or β-glucose-1-phosphate.

The first radiotracer we aimed to synthesize using this enzymatic method was the maltose derived tracer 2-deoxy-[^18^F]-fluoro-maltose ([^18^F]FDM). Our interest in this probe was based on the potential specificity of microbial maltose metabolism^29–31^ and the recent development of maltodextrin transporter-targeted imaging methods^32–35^. Maltodextrin (D-glucose units with α-1,4 glycosidic linkages) and its structural relatives are important energy sources for bacteria. These oligosaccharides are taken up by the maltodextrin transporter, which are present in both gram-positive and gram-negative bacterial species but not found in mammalian cells. In order to enzymatically obtain [^18^F]FDM from [^18^F]FDG, we focused on commercially available maltose phosphorylase (E.C. 2.4.1.8), which has been used to synthesize a variety of disaccharides^36–40^.

In this work, we report the one step radiosyntheses of [^18^F]FDM and α-1,3-product, 2-deoxy-2-[^18^F]-fluoro-sakebiose ([^18^F]FSK), from easily accessible [^18^F]FDG, using maltose phosphorylase as catalyst. Both newly reported [^18^F]-labeled radiotracers were accumulated by important human pathogens including *S. aureus* and were specific to bacterial infection *in vivo*. This phosphorylase-catalyzed method was extended to additional [^18^F]-labeled disaccharides of high biomedical interest including 2-deoxy-2-[^18^F]fluoro-trehalose ([^18^F]FDT), 2-deoxy-2-[^18^F]fluoro-laminaribiose ([^18^F]FDL), and 2-deoxy-2-[^18^F]fluoro-cellobiose ([^18^F]FDC).

## RESULTS

### Maltose phosphorylase-catalyzed radiosynthesis of [^18^F]FDM from clinical [^18^F]FDG showed the α-1,3-product [^18^F]FSK as a minor product

Our initial goal was to demonstrate that the phosphorylase-catalyzed chemoenzymatic synthesis of an [^18^F] disaccharide from [^18^F]FDG was a viable alternative to conventional radiochemical approaches. Although several bacteria-specific glycans could be constructed chemoenzymatically, we focused on maltose-derived PET radiotracers given the strong literature precedent^33–35^. We therefore synthesized the [^18^F]FDG-derived, 2-position labeled [^18^F]FDM to target the bacterial maltose receptor. First, the precursor, β-D-glucose-1-phosphate (βGlc1-P) was synthesized in three steps starting from acetobromo-α-D-glucose with 75% overall yield (**Figure 2A**). Cyclotron-produced [^18^F]FDG was then added to a mixture of βGlc1-P and maltose phosphorylase in citrate buffer (0.1M, pH=6.0), and stirred at 37°C for 20 minutes. The reaction led to the formation of [^18^F]FDM with 72% ± 4% decay-corrected radiochemical yield (RCY) and the α-1,3-linked product [^18^F]FSK with 15%± 3% decay-corrected RCY (*N* = 25) (**Figure 2B**). Each [^18^F]-labeled product was isolated using semi-preparatory HPLC (**Figure 2C**) followed by formulation for subsequent *in vitro* and *in vivo* studies.

**Figure 2.**
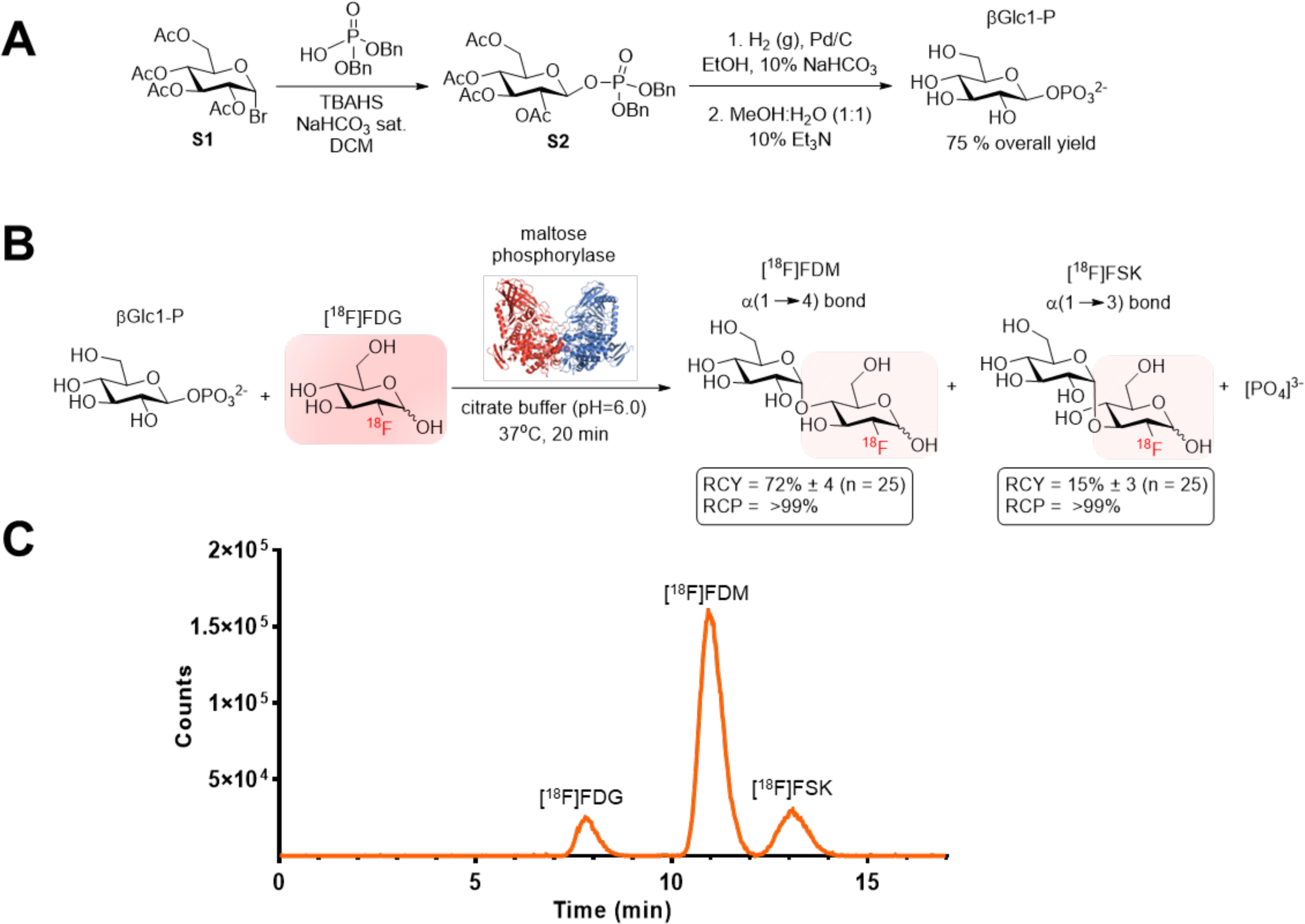
Radiochemical syntheses of 2-deoxy-2-[^18^F]fluoro-maltose ([^18^F]FDM) and 2-deoxy-2-[^18^F]fluoro-sakebiose ([^18^F]FSK). (A) Synthesis of β-D-glucose-1-phosphate (βGlc1-P) precursor starting from bromide **S1**. (B) Enzymatic radiosynthesis of [^18^F]FDM and [^18^F]FSK from 2-deoxy-2-[^18^F]fluoro-D-glucose [^18^F]FDG and βGlc1-P using maltose phosphorylase. (C) Radio HPLC analysis of crude products using a YMC-Pack Polyamine II column.

### Phosphorylase-catalyzed radiosyntheses of [^18^F]FDT, [^18^F]FDL and [^18^F]FDC from clinical [^18^F]FDG

Several additional [^18^F]-labeled disaccharides might be used to image important human pathogens *in vivo*. The unique metabolism of the α,α-1,1 linked disaccharide trehalose by *Mycobacterium tuberculosis* has been previously targeted for therapy and imaging^41–44^. The β-linked disaccharides laminaribiose and cellobiose could potentially be leveraged for the metabolic imaging of fungal infections given the presence of β-1,3 and β-1,4 linkages in β-D-glucans (BDG), found in the cell walls of fungi^45–47^. An existing clinical assay (serum and cerebrospinal fluid) detects fungal β-1,3 linkages via the coagulation cascade of the horseshoe crab to detect invasive fungal infections caused by *Aspergillus* and *Candida* species^48^. With these potential imaging applications in mind, we attempted [^18^F]-radiosyntheses of the additional disaccharides indicated in **Figure 1A**. The use of trehalose-, cellobiose- and laminaribiose-phosphorylase in combination with [^18^F]FDG and adequate α− or β-D-glucose-1-phosphate precursor, led to the formation of [^18^F]FDT (decay-corrected RCY 62% ± 4%; *N* = 5), [^18^F]FDL (decay-corrected RCY 96% ± 3%; *N* = 5), and [^18^F]FDC (decay-corrected RCY 81% ± 7%; *N* = 5) (**Figure 3**). Of note, direct radiosynthesis of [^18^F]FSK via sakebiose phosphorylase produced only a modest yield of the desired product (decay-corrected RCY 5% ± 3%; *N* = 5). The [^18^F]FSK tracer was therefore obtained via maltose phosphorylase as described above, for subsequent *in vitro* and *in vivo* studies.

**Figure 3.**
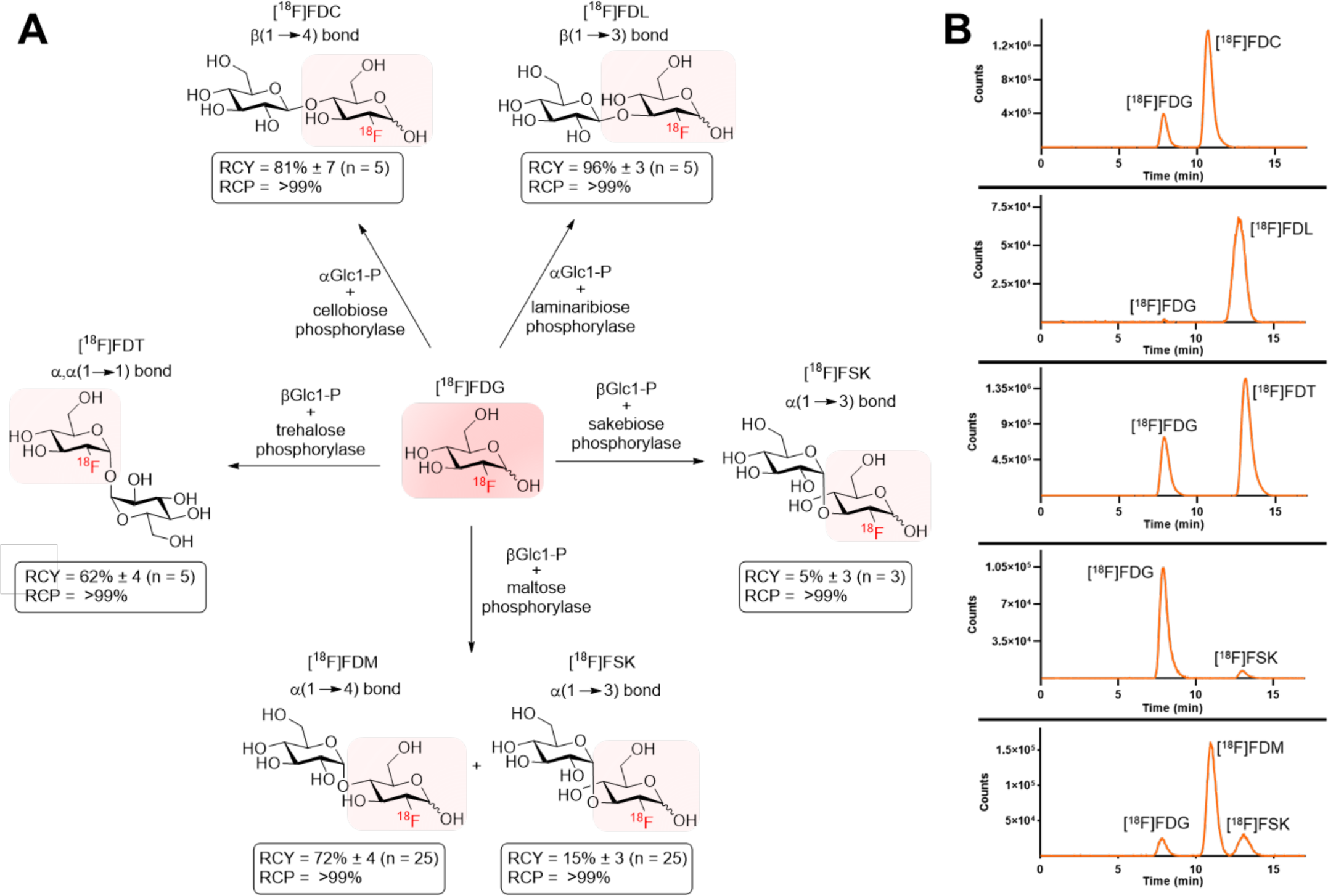
(A) Chemoenzymatic radiosyntheses of [^18^F]FDT, [^18^F]FDL, [^18^F]FDC, [^18^F]FDM and [^18^F]FSK from [^18^F]FDG. All reactions were carried out at 37°C, stirring for 20 min, using 6 mg (0.020 mmol) of precursor, 0.3 mg of enzyme (3-6 units) and 10-15mCi [^18^F]FDG in 0.5 mL of citrate buffer (0.1M, pH = 6.0). (B) Radio HPLC analysis of each enzymatic reaction using a YMC-Pack Polyamine II column.

### In vitro studies using [^18^F]FDM and [^18^F]FSK showed high tracer accumulation in key human pathogens including S. aureus and A. baumannii

We hypothesized that the disaccharides [^18^F]FDM and [^18^F]FSK would have different microbial sensitivity from previously reported tracers, with accumulation in the key human pathogen *S. aureus*. We therefore studied these tracers *in vitro* to assess incorporation in a variety of clinically relevant bacteria and establish microbial specificity. The gram-positive bacteria *S. aureus* and the gram-negative *Enterobacteriaceae Escherichia coli* and *Klebsiella pneumoniae* were used to assess assimilation of [^18^F]FDM and [^18^F]FSK and compare to their uptake to that of reported PET tracers [^18^F]FDG and [^18^F]FDS (**Figure 4A**). The glucose derivative [^18^F]FDG, used frequently in the clinic for oncologic and neuroimaging applications, was incorporated into all three bacteria. In contrast, as expected [^18^F]FDS accumulated in *E. coli* and *K. pneumoniae* but not *S. aureus*^4^. Both [^18^F]FDM and [^18^F]FSK showed high incorporation in *S. aureus* and *K. pneumoniae,* similar to that of [^18^F]FDG but low uptake in *E. coli*. Further bacteria pathogens were tested with [^18^F]FDM and [^18^F]FSK. (**Figure 4B**). Overall, the two radiotracers demonstrated similar sensitivities towards gram-positive and gram-negative bacteria. They showed high uptake in *Staphylococcus epidermis*, *A. baumannii* and *Enterobacter cloacae*, but low uptake in *Listeria monocytogenes, Pseudomonas aeruginosa*, *Salmonella typhimurium* and *Proteus mirabilis*. There was only background level of incorporation of [^18^F]FDM and [^18^F]FSK into heat-killed bacteria for all species studied. (**Supp. Fig. S1**). [^18^F]FDM and [^18^F]FSK demonstrated increased bacterial accumulation over time in bacterial cultures (**Supp. Fig. S2**), and showed a high degree of retention in “efflux” experiments (**Supp Fig. S3**). To determine if the differential uptake of [^18^F]FDM and [^18^F]FSK by bacteria reflected differential use of sugar transporters, *S. aureus* cultures were incubated with [^18^F]FDM and [^18^F]FSK in the presence of increasing concentrations of unlabeled maltose (**Figure 4C**), suggesting that they both use a maltodextrin transporter for uptake. To determine if there was strain-to-strain variability in tracer accumulation, we compared uptake in 4 different clinical isolates of methicillin-resistant *S. aureus* (MRSA) (**Figure 4D**). Three of the isolates exhibited similar levels of uptake for both tracers, whereas one isolate showed diminished uptake of [^18^F]FDM.

**Figure 4.**
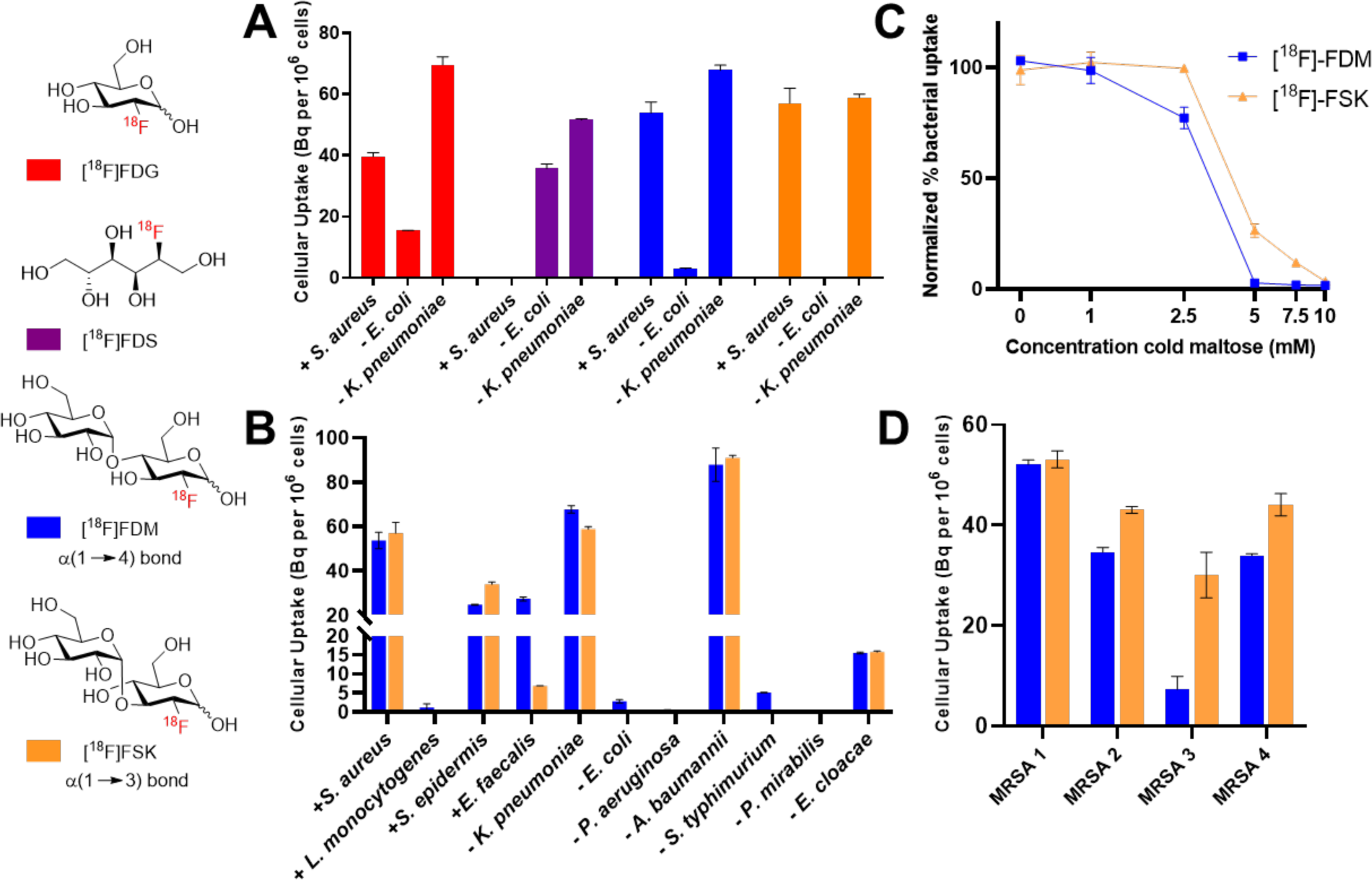
(A) *In vitro* bacteria uptake for [^18^F]FDG, 2-deoxy-2-[^18^F]-fluoro-D-sorbitol ([^18^F]FDS), [^18^F]FDM and [^18^F]FSK. (B) *In vitro* bacteria uptake of [^18^F]FDM and [^18^F]FSK in gram-positive and gram-negative pathogens. (C) Competition of [^18^F]FDM and [^18^F]FSK uptake with increasing concentrations of unlabeled (cold) maltose in *S. aureus.* (D) *In vitro* bacteria uptake of [^18^F]FDM and [^18^F]FSK in methicillin-resistant *S. aureus* (MRSA) clinical strains.

### [^18^F]FDM and [^18^F]FSK is stable in human serum and its degradation in mouse serum can be abrogated by use of an α-glucosidase inhibitor

In preparation for *in vivo* studies in mice and humans, we assessed the stability of [^18^F]FDM and [^18^F]FSK incubated with mouse and human serum using radio HPLC. In mouse serum, both radiotracers exhibited increasing time-dependent hydrolysis to [^18^F]FDG, while they remained stable in human serum. (**Supp. Fig. S4**). This observation may be explained by the increased abundance of α-glucosidase, which has been reported to hydrolyze maltodextrin-based tracers^49^, in murine versus human serum. The α-glucosidase enzyme is predicted to degrade maltose from the non-reducing end, hydrolyzing [^18^F]FDM and [^18^F]FSK to glucose and [^18^F]FDG (**Supp. Fig. S5**). In contrast, α-amylase (EC 3.2.1.1), which is present in human serum, degrades maltodextrin from the reducing end. Unlike longer chain maltodextrin-based tracers (>3 units), [^18^F]FDM and [^18^F]FSK would be anticipated to be resistant to α-amylase and thus stable in human serum^49^.

We therefore tested the ability of the α-glucosidase inhibitors voglibose, acarbose and miglitol (**Supp. Fig. S6**), which are commonly used as diabetes treatment, to prevent the α-glucosidase-mediated degradation of [^18^F]FDM and [^18^F]FSK in mouse serum. First, we verified that voglibose and miglitol did not affect accumulation of [^18^F]FDM and [^18^F]FSK in culture-grown *S. aureus*, whereas acarbose did (**Supp. Fig. S7**). We then compared inhibitor potency in mouse serum and found that voglibose was the most potent inhibitor to prevent [^18^F]FDM and [^18^F]FSK degradation (**Supp. Fig. S8**). Together, these studies suggested that the concurrent administration of voglibose with [^18^F]FDM and [^18^F]FSK in murine studies allowed a better approximation for future human performance.

### In vivo analysis of [^18^F]FDM and [^18^F]FSK in a murine model of bacterial infection demonstrated higher signal to background for the sakebiose-derived tracer

We first tested [^18^F]FDM and [^18^F]FSK in non-infected mice, to both assess tracer stability and to detect potential contributions of the normal microbiome to tracer signals. When voglibose was added to tracer injection, *in vivo* analysis of [^18^F]FDM and [^18^F]FSK in conventionally-raised mice^10^ (*N* = 5), which have bacteria colonization of their gut, showed background signal only in kidney and bladder (**Supp. Fig. S9**). In contrast, when voglibose was omitted from the injection of [^18^F]FDM and [^18^F]FSK, the resulted signal was similar to [^18^F]FDG, with high heart and brain uptake, demonstrating the utility of the inhibitor (**Supp. Fig. S10**).

We next evaluated a murine model of acute bacterial infection to test whether [^18^F]FDM and [^18^F]FSK uptake could detect live bacteria in *in vivo* mouse models of infection. We chose the MRSA myositis model as it has been studied extensively in tracer development and used to compare tracer accumulation in infected tissues (harboring live bacteria) versus sterile inflammation (reflecting the host immune response)^4, 10, 11, 14^. Mice were inoculated with live MRSA in the left shoulder and with a 10-fold higher dose of heat-killed MRSA in the right shoulder. Following tracer injection in the presence of voglibose, both [^18^F]FDM and [^18^F]FSK accumulated at the site of injection of live MRSA but not of heat-killed MRSA (*N* = 6 for each tracer) (**Figure 5**). The region-of-interest (ROI) analysis revealed [^18^F]FDM and [^18^F]FSK uptake at the side of live MRSA injection was 6.1 and 6.5-fold higher (*P* <0.0001) respectively than at the side of heat-killed MRSA injection. These results were further corroborated by *ex vivo* analysis (tissue harvesting and gamma counting) which showed that the mean [^18^F]FDM and [^18^F]FSK accumulation in tissues inoculated with live MRSA was respectively 3.8 and 4.7-fold higher than that seen for tissues inoculated with heat-killed bacteria (*P* = 0.001 for [^18^F]FDM and *P* value < 0.0001 for [^18^F]FSK) (**Supp. Fig S11-S12**). In contrast, the glucose-transporter targeted parent tracer [^18^F]FDG accumulated equally at injection sites inoculated with either live or heat-killed MRSA, as quantified by *in vivo* and *ex* vivo analysis (**Supp. Fig S13**). We thus conclude that both the [^18^F]FDM and [^18^F]FSK tracers are specific for live versus heat-killed MRSA and have the potential to distinguish bacterial inflammation from sterile inflammation. However, as [^18^F]FSK proved to have better *in vivo* performance than [^18^F]FDM, it was chosen for testing in additional preclinical models.

**Figure 5.**
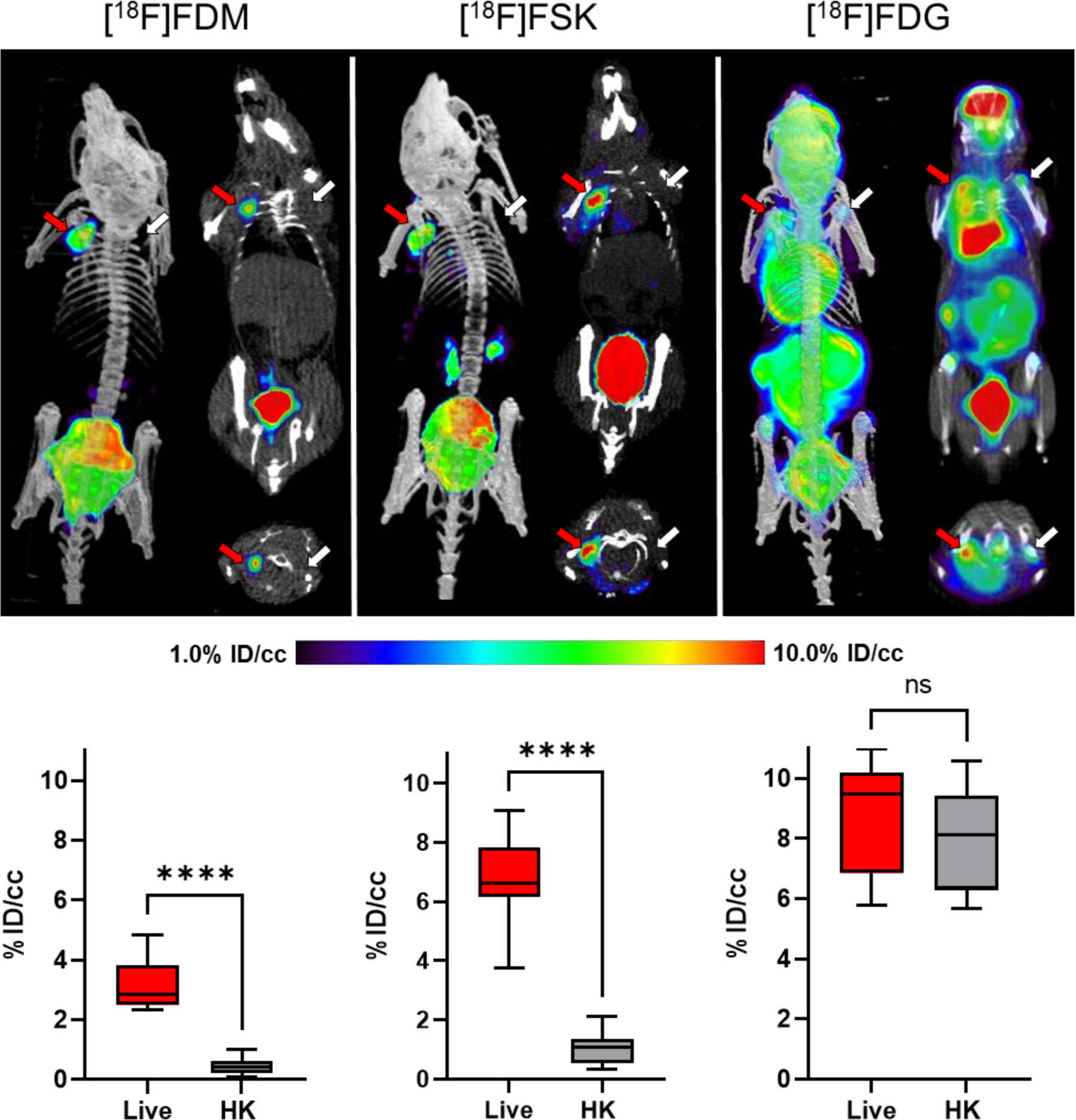
μPET-CT imaging of MRSA myositis in mice with [^18^F]FDM, [^18^F]FSK and [^18^F]FDG. The red arrows indicate the site of inoculation with live bacteria, while the white arrows correspond to heat-killed bacteria. The corresponding bar graphs indicate region-of-interest (ROI) analysis. As reflected by the images, the mean [^18^F]FDM and [^18^F]FSK accumulation for tissues infected with live bacteria was respectively 6.1 and 6.5-fold higher than that seen for heat-killed inoculation (*P* <0.0001). In contrast, this difference was not seen for [^18^F]FDG.

### Preclinical models of vertebral discitis-osteomyelitis, and A. baumannii myositis suggested that [^18^F]FSK could be used in challenging clinical settings

Based on our encouraging results with MRSA *in vitro* and *in the murine myositis model*, we next studied a rat model of vertebral discitis-osteomyelitis using [^18^F]FSK. Rats were inoculated with live *S. aureus* (Xen-29, bioluminescent strain) in the third intervertebral space from the base of the tail and with heat-killed *S. aureus* (Xen-29) in the fifth intervertebral space. The tail was imaged by luminescence emission using a Xenogen IVIS® 50 to verify the location and presence of the bacterial infection (**Figure 6a**). After 4 days, CT was used to image damage to the disc and bone, revealing the development of disc-space narrowing and end-plate degeneration, which mimics human bacterial spinal infections (**Figure 6b**). Imaging of the tail using [^18^F]FSK was performed at day 4 (*N* = 5) and day 10 (*N* = 3) following inoculation of bacteria (**Figure 6c and Supp. Fig. S14-S15**). In both cases [^18^F]FSK accumulated at the infection site, with ROI analyses demonstrating 2.8-fold higher signals at day 4 (*P* <0.0001) and 3.1-fold higher signals at day 10 (*P* <0.0001), in the third intervertebral space versus the fifth intervertebral space (**Figure 6d and Supp. Fig. S15**).

**Figure 6.**
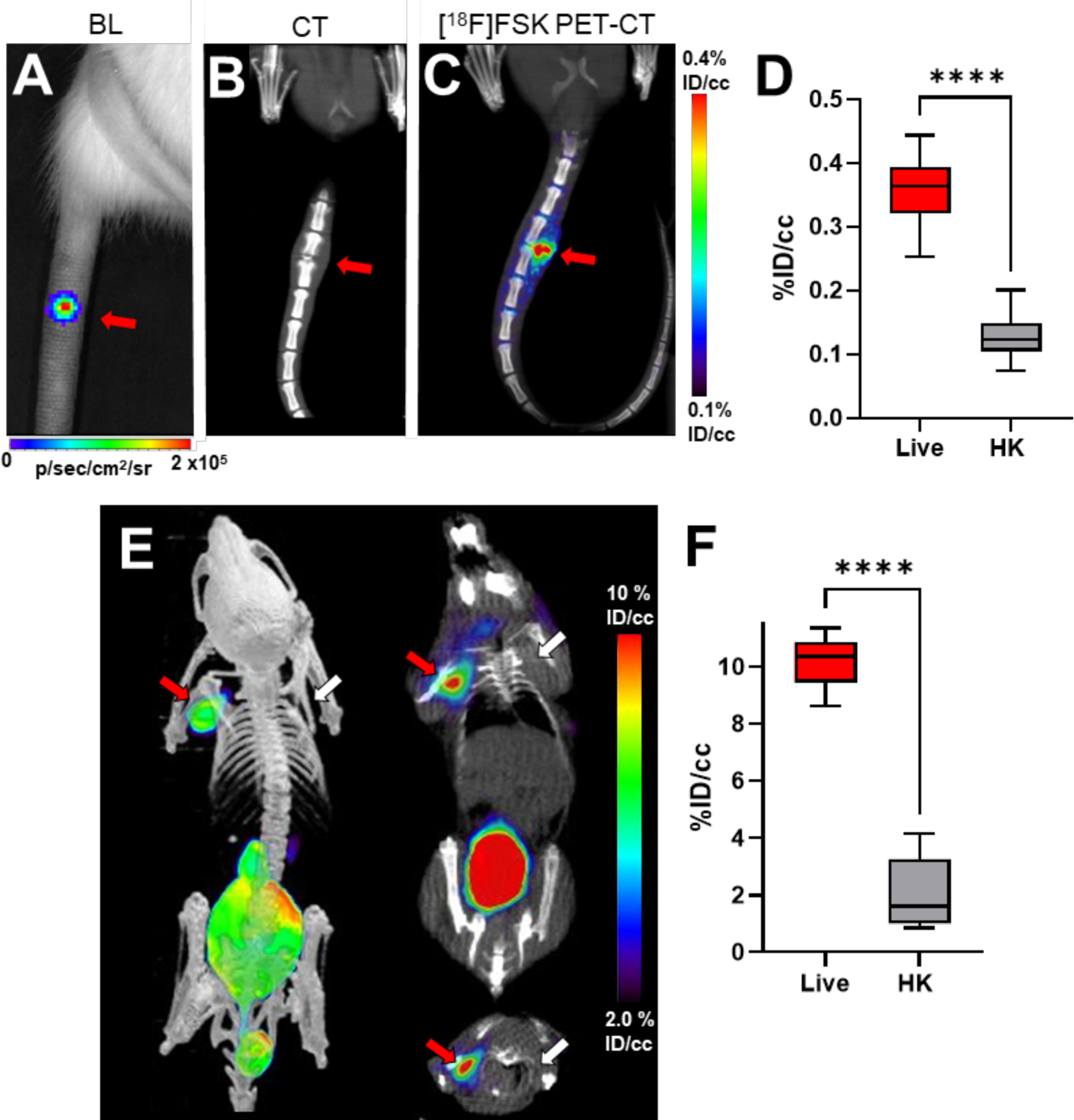
Imaging of *S. aureus* in vertebral discitis-osteomyelitis (VDO) rat models and *A. baumannii* in a myositis mouse model using [^18^F]FSK. (A) Optical tomography image of rat tail showing bioluminescent signal from *S. aureus* Xen29 inoculation. (B) Computed tomography study performed at 10 days highlights the similarity between rodent and human discitis osteomyelitis. (C) PET/CT imaging of *S. aureus* Xen29 vertebral discitis-osteomyelitis (VDO) in rat (*N* = 5) with [^18^F]FSK. (D) ROI analysis showing increased signal in segments inoculated with live bacteria versus background (*P* < 0.0001). (E) PET/CT imaging of *A. baumannii* myositis in mice (*N* = 6) with [^18^F]FSK. The red arrows indicate the site of inoculation with live bacteria, while the white arrows correspond to heat-killed bacteria. (F) ROI analysis showing increased signal in infected muscle versus inflammation (*P* < 0.0001). *For Table of Contents Use Only*

After showing the efficacy of [^18^F]FSK *in vivo* to detect *S. aureus* infection, we further studied *A. baumannii* in the murine myositis model, as it is a common cause of soft tissue infections in the battlefield. We injected *A. baumannii* as described above, and performed μPET/CT following [^18^F]FSK injection. The tracer accumulated at the site of live *A. baumannii* injection into the left shoulder but not at the site of heat-killed *A. baumannii* injection into the right shoulder (**Figure 6e**). ROI analysis revealed [^18^F]FSK accumulation in the live *A. baumannii* injected muscle was 4.9-fold higher than at the heat-killed *A. baumannii* injected muscle (*P* <0.0001) (**Figure 6f**). *Ex vivo* analysis revealed that tracer uptake at the site of live *A. baumannii* injection was 3.9-fold higher compared to the site of heat-killed *A. baumannii* injection (*P* <0.0001) (**Supp. Fig. S16**). Together, these studies employing two clinically important human bacterial pathogens, *S. aureus* and *A. baumannii*, in two different murine models that mimic challenging to treat human infections, demonstrate the potential of using [^18^F]FSK to image human bacterial infections in the clinic.

## DISCUSSION

Bacteria-specific metabolic pathways have been exploited by antimicrobial agents for decades, with infection imaging a more recent application of this approach. In recent years, numerous compelling methodologies have been further validated in patients most notably [^18^F]FDS, which is highly sensitive for *Enterobacteriaceae* and which can be efficiently radiosynthesized from the common tracer [^18^F]FDG^20^, making it a more practical tool for potential clinical use. To further impact infectious disease management in the clinic, we will require imaging tools with both straightforward radiosyntheses, and applicability to a broader range of bacteria, including the common human gram-positive bacteria *S. aureus*.

Tracers labeled with radionuclides with a shorter half-life [^11^C; t_1/2_ = 20 min]^10–12, 14^ or studied primarily in academic centers [^89^Zr; t_1/2_ = 78 hrs]^50^ face significant challenges in the acute care/ emergency setting, highlighting the need for infection imaging methods that might be more broadly useful. We therefore developed a chemoenzymatic method for pathogen-targeted PET radiotracers, benefitting from (1) the efficiency of chemoenzymatic reactions, versus standard PET radiochemical methods and (2) the general availability of [^18^F]FDG as a synthon. A major challenge in the chemical synthesis of [^18^F]-labeled carbohydrates is the short half-life (109.7 min) of fluorine-18, requiring complex precursors, late-stage S_N_2 radiofluorination via [^18^F]-fluoride, and less sterically hindered labeling sites^23^. In contrast, the current report describes a rapid (20 min), one-step radiosynthesis using [^18^F]FDG and commercially available precursors, with high regioselectivity/stereoselectivity and potential metabolic advantages conferred via 2-position [^18^F] labeling. With the goal of developing an on-demand *S. aureus*-sensitive tracer, we used maltose phosphorylase to dimerize [^18^F]FDG into the α-1,4 and α-1,3 disaccharides [^18^F]FDM. and [^18^F]FSK. The sakebiose-derived tracer showed outstanding performance characteristics in its ability to detect living bacteria *in vivo* including *S. aureus*.

Sakebiose (also known as nigerose) has been described as an “uncommon sugar,” and investigated for its potential as an alternative sweetener, oral probiotic and immunopotentiating therapy^51–53^. Most published work on sakebiose has focused on production of the disaccharide via enzymatic degradation of dextrans or synthesis using phosphorylases, whose activity may be modified via mutagenesis^54–57^. Little is known about its transport and metabolism, although the current study suggests that sakebiose (at least the described 2-[^18^F] derivative) shares transport mechanism(s) with maltose for the bacteria studied, since [^18^F]FDM and [^18^F]FSK incorporation were inhibited by cold maltose to a similar degree *in vitro*. Further studies are needed to better understand its mechanism of bacterial incorporation, which may drive the discovery of additional metabolic imaging tools. The described radiosynthesis of [^18^F]FSK does indicate that the chemoenzymatic construction of complex PET radiotracers from common synthons including [^18^F]FDG is feasible, to the extent that the required tracers, glycan-derived “acceptors,” and enzymes are available. Furthermore, the reported engineering of disaccharide phosphorylases suggests that changing the distribution of [^18^F] products obtained may be possible. In the current case, maltose phosphorylase could be altered to increase the radiochemical yield of the desired [^18^F]FSK tracer.

As stated previously, we believe that [^18^F]FDM and [^18^F]FSK have outstanding potential as clinical PET tracers, especially for the detection and monitoring of *S. aureus* infections. Both can be produced quickly and efficiently from widely available [^18^F]FDG, without the need for regioselective precursor modification and chemical protection/ deprotection. Most interestingly, the 2-position ^18^F substituted maltose derivative appears to confer numerous advantages over the 6-position ^18^F derivative originally reported by Namavari *et al*.^34, 58^, in terms of stability and microorganism export. Although the 6-position is more chemically accessible, the corresponding 6-[^18^F] derivative is both more vulnerable to defluorination, and prohibits 6-position phosphorylation, which is a major mechanism of [^18^F]FDG retention (via hexokinase, E.C. 2.7.1.1). Indeed, in terms of *in vitro* stability, bacterial export, and *in vivo* performance [^18^F]FDM and [^18^F]FSK more closely mimic the “second generation” PET tracer 6”-[^18^F]-fluoromaltotriose^35, 59^. Explicit comparison between [^18^F]-labeled α-1,4 linked oligomers (maltose, maltotriose, and maltodextrin) would be helpful to guide clinical implementation and is the basis of ongoing laboratory efforts.

In conclusion, we have developed chemoenzymatic method for the radiosyntheses of [^18^F]-labeled disaccharides from the readily available precursor [^18^F]FDG. The strategy was used to generate both α- and β-linked [^18^F] disaccharides of high biomedical interest for pathogen-specific imaging, specifically [^18^F]-labeled derivatives of maltose, sakebiose, trehalose, laminaribiose and cellobiose. We anticipate that this approach may be used to construct other complex [^18^F] glycans and facilitate the on-demand radiosynthesis of PET radiotracers for infection and other diseases.

## MATERIALS AND METHODS

### General Chemistry and Radiochemistry

Full descriptions of chemical and radiochemical syntheses as well as the analytical techniques used are provided in the Supporting Information. Unless otherwise noted, all reagents were obtained commercially and used without further purification. Radioisotopes were generated in the UCSF radiopharmaceutical facility.

### Synthesis of **β**-D-glucose-1-phosphate (**β**Glc1-P)

β-D-glucose-1-phosphate (βGlc1-P) was synthesized in three steps from 1-bromo-α-D-glucose tetraacetate (75 % overall yield, gram scale). For detailed methods and characterization of each compound, please see the Supporting Information.

### General enzymatic radiosynthesis of [^18^F]FDC, [^18^F]FDL, [^18^F]FDT, [^18^F]FDM and [^18^F]FSK

In a 4 mL borosilicate vial containing a PTFE stir bar, phosphorylase (0.3-0.5 mg, 3-6 units) and α− or β−Glc1-P (6 mg, 20 mmol) were added. A solution of [^18^F]FDG (15-30 mCi) in citrate buffer (pH=6.0) was directly transferred to the vial and the mixture was stirred at 37 °C for 20 min. The mixture was diluted with MeCN then filtered through a C18 light cartridge, and subsequently purified via semi-preparative HPLC using a YMC column Polyamine Pack II, 10 mm (73% MeCN/27 % H_2_O). The [^18^F]-labeled disaccharide product was isolated in 5-7 mL fractions. The fractions were then diluted with MeCN (40 mL) before being passed through a Sep-pak Plus NH_2_ Cartridge at 5 mL/min to trap each dimer product. After flushing the cartridge with air and N_2_ gas, the tracer was eluted using a saline solution for direct formulation for *in vitro* or *in vivo* use. Radiochemical yields and purity of [^18^F]-product were confirmed by analytical HPLC. The synthesis and characterization of cold standards are described in the Supporting Information.

### Uptake of [^18^F]FDM and [^18^F]FSK in gram-positive and gram-negative bacteria *in vitro*

S. aureus, L. monocytogenes, S. epidermidis, E. faecalis, K. pneumoniae, E. coli, P. aeruginosa, A. baumannii, S. typhimurium, P. mirabilis and E. cloacae were grown overnight in lysogeny broth (LB) in a shaking incubator at 37 °C. Overnight cultures were diluted to an optical density at 600 nm (OD_600_) of 0.05 and grown to exponential phase (∼ 0.4-0.6). Bacterial cultures were incubated with 24 µCi of [^18^F]FDM and [^18^F]FSK at 37 °C for 90 minutes for uptake studies and also at 30 min and 60 min for temporal evaluation. After tracer incubation, 500 µL of the bacterial cultures were transferred to Spin-X LC 1.5 mL tubes (0.22 µm) and were centrifuged (6 min, 13200 rpm) to separate bacterial cells and supernatant. Bacterial cells were then washed 1x with phosphate buffered saline (PBS) to remove any tracer not taken up by bacteria. Heat-killed bacterial samples used as control were prepared by incubating the bacterial cultures at 90 °C for 30 min. Retained radiotracer within samples was then counted using an automated gamma-counter (Hidex). Blocking experiments were performed by adding cold maltose (0.01 mM to 10 mM) together with 24 µCi of [^18^F]FDM and [^18^F]FSK and following the same protocol. Efflux experiments were performed by incubating the bacteria with 24 µCi of [^18^F]FDM and [^18^F]FSK for 30 minutes, then pelleting the bacteria and replacing the media with fresh LB. The cultures were then incubated for an additional 30 min, then similar method was used to separate bacteria cells and supernatant. Radioactivity for both was counted using a gamma-counter (HIDEX) to obtain residual activity.

### Animal experiments

All animal procedures were approved by the UCSF Institutional Animal Care and Use Committee and were performed in accordance with UCSF guidelines. CBA/J mice (female, 8-10 weeks old) and Sprague/Dawley rats (female, 10–12 weeks old) were used for the experiments. Mice and rats were housed in individually ventilated cages under normal diet in groups of 3 rats or 5 mice, with ad libitum access to food and water throughout the experiment. Prior to infection and during imaging the animals were anesthetized with 5% isoflurane. Mice and rats were inoculated with *S. aureus* Xen29, MRSA, *K. pneumoniae*, or heat-killed bacteria as described previously^10^. At different time points further specified below, the mice and rats were imaged using a Xenogen IVIS® 50 instrument or Inveon µPET-CT following the injection of [^18^F]FDM, [^18^F]FSK or [^18^F]FDG.

### *In vivo* [^18^F]FDM and [^18^F]FSK dynamic imaging in a myositis mouse model

Mice were inoculated with MRSA or *A. baumannii* (∼ 2×10^7^ CFU) in the left deltoid muscle and 10-fold higher bacterial load of heat-killed bacterial in the right deltoid muscle. After 12h, [^18^F]FDM or [^18^F]FSK was injected via teil vain (∼ 100 μL, 200 μCi, containing 1 mg voglibose). The mice were then imaged by µPET-CT using the same protocol previously described^10^ (90 min dynamic PET scan, 5 min CT). The resulting µPET-CT images were analyzed with Amide’s Medical Image Data Examiner as described below.

### *In vivo* [^18^F]FSK dynamic imaging in a VDO rat model

*S. aureus* Xen29 was used to induce discitis in 5 Sprague/Dawley Rats (Charles River). Xen29 is a bioluminescent *S. aureus* strain that carries a stable copy of the *Photorhabdus luminescens* lux operon (*luxABCDE*) and was used in the study to verify successful bacterial inoculation in the third intervertebral space. Xen29 was grown overnight in LB containing 100 μg/L of kanamycin as aforementioned described and diluted in PBS to obtain the desired bacterial load for infection (∼ 2×10^7^ colony forming units, CFU). Rats (*N* = 5) were inoculated with Xen29 live and heat-killed (10-fold higher bacterial load) in the third and fifth intervertebral space respectively. At days 0 and 2, the rats were imaged using a Xenogen IVIS® 50 imaging system to detect bioluminescence signal and confirm the infection. At days 4 and 10, [^18^F]FSK injection (∼ 200 μL, 500 μCi, 5 mg Voglibose) was performed using a tail vein catheter. One hour post tracer injection, rats were transferred to the µPET-CT system (Siemens) and imaged using a 90 min dynamic PET acquisition scan followed by a 5 min µCT scan for attenuation correction and anatomical co-registration. Anesthesia was maintained during bioluminescence and µPET-CT imaging using 5% isofluorane. Resulting µPET-CT images were analyzed using AMIDE and %ID/cc was used for quantitative comparison. %ID/cc values were established via 8 mm^3^ region of interest ROI’s using the spherical tool. Region of interest (ROI) analysis from resulting µPET-CT images was used to compare tracer performance. Resulting bioluminescence images were analyzed with Living Image Software 3.2.

### Data analysis and statistical considerations

For synthesis, radiochemical yield incorporates decay-correction for ^18^F (t_1/2_=109.7 min). *In vitro* data was normalized to CFU’s for sensitivity analysis to account for differential growth rates between organisms. All *in vivo* PET data were viewed using open-source AMIDE software. Quantification of uptake was performed by drawing spherical regions of interest (5-8 mm^3^) over indicated organs on the CT portion of the exam and expressed as percent injected dose per gram. All statistical analysis was performed using GraphPad Prism v 9. Data were analyzed using an unpaired two-tailed Student’s t-test. All graphs are depicted with error bars corresponding to the standard error of the mean.

## Supporting Information

Detailed information regarding synthesis, *in vitro*, and *in vivo* experiments not reported in the main text.

## Author contributions

DMW, JE, MO, and JSF proposed and supervised the overall project. AMS, MLA, AAA, KWB obtained *in vitro* data or supported *in vitro* studies. AMS, SJR, MLA and JB performed the chemistry and radiochemistry. TD and BN supplied phosphorylase enzymes. AMS, MLA, RS, KNB and SS performed μPET-CT imaging studies and subsequent data analysis. AMS, MLA performed *ex vivo* analysis. AMS, MLA, DAH, RRF, JE, OSR, JSF, DMW wrote and edited the paper.

## Acknowledgements

The authors would like to thank Drs. Gayatri Gowrishankar, Niren Murthy, Bin Shen, Dima Hammoud and Sanjay Jain for helpful discussions.

## Notes

The authors declare no competing financial interests.

---

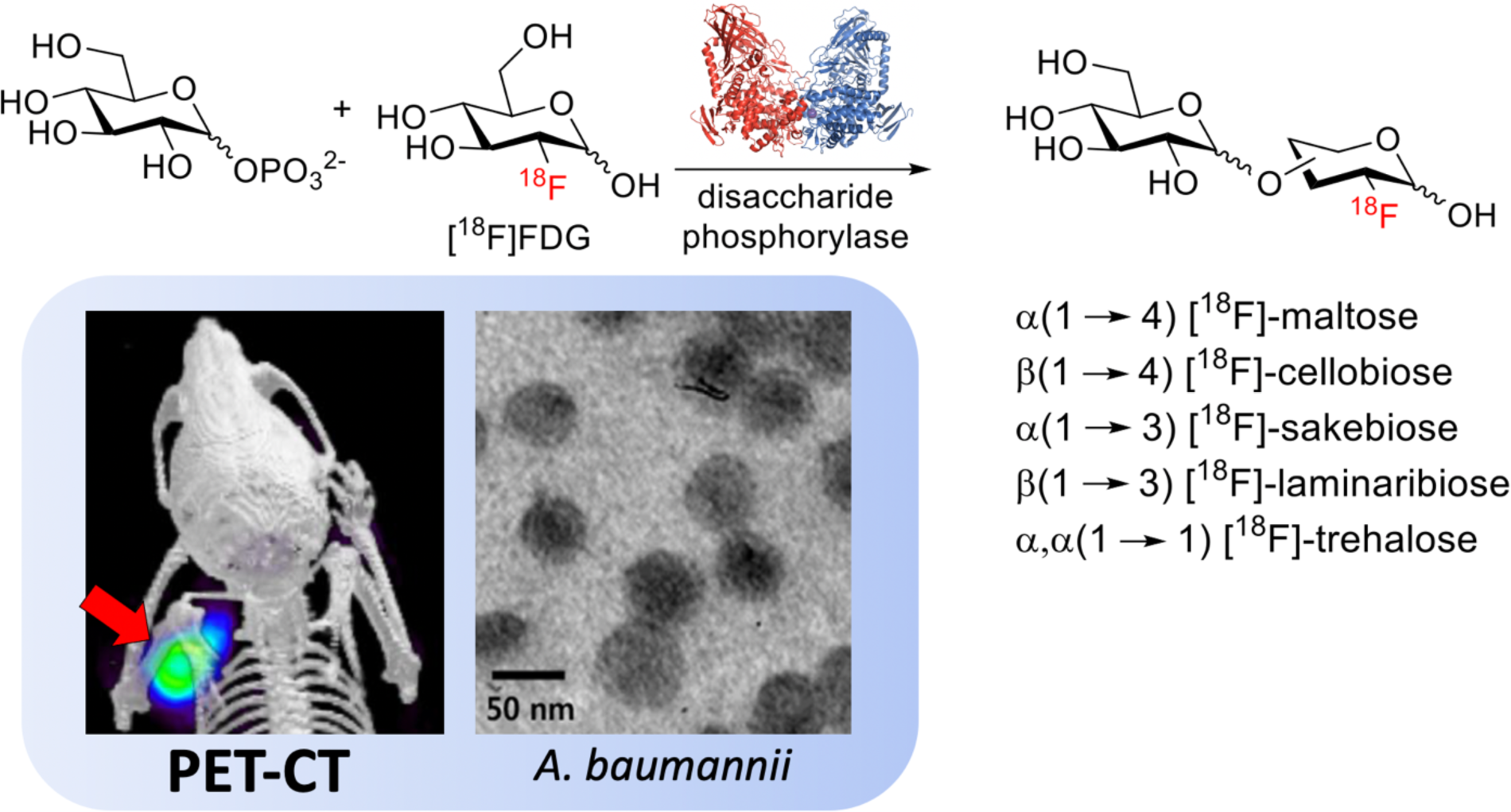
Synopsis. This manuscript reports the rapid chemoenzymatic syntheses of five [^18^F]-labeled disaccharides from the common oncologic radiotracer [^18^F]FDG. Both [^18^F]FDM (α-1,4) and [^18^F]FSK (α-1,3) tracers showed high accumulation in important clinical pathogens *in vitro* and *in vivo* with the lead tracer [^18^F]FSK representing a strong candidate for near-term clinical translation.

