## Supplemental Information for "Chemoenzymatic syntheses of fluorine-18-labeled disaccharides from [^18^F]FDG yield potent sensors of living bacteria *in vivo*"

### A. Supplemental Figures

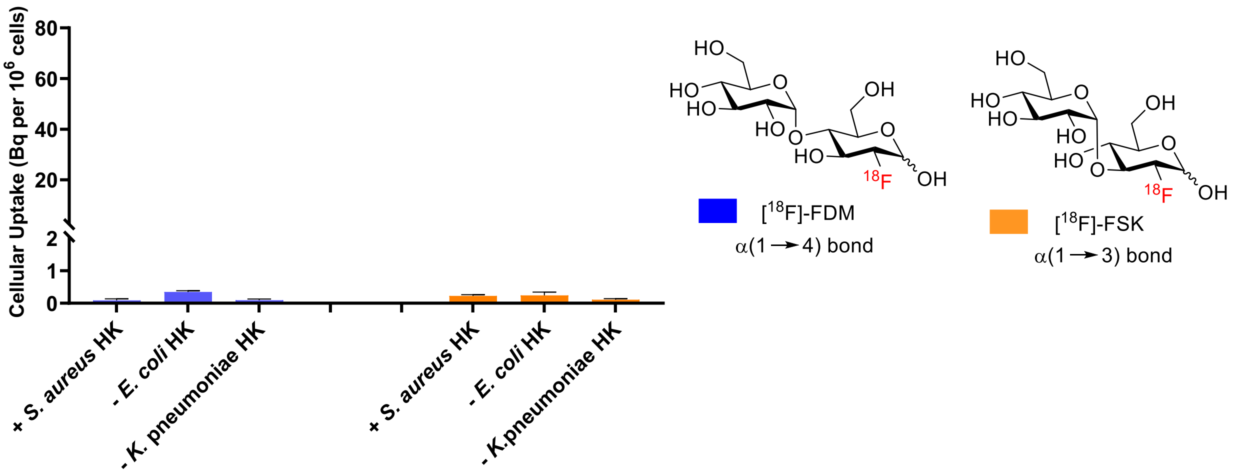

**Figure S1.** *in vitro* uptake of [$^{18}$F]FDM and [$^{18}$F]FSK in heat-killed bacteria.

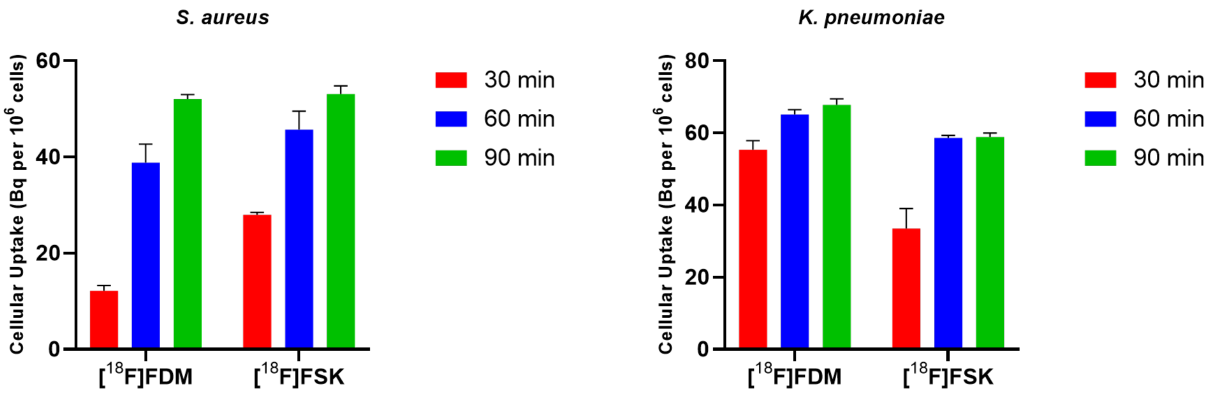

**Figure S2.** *in vitro* bacteria uptake [$^{18}$F]FDM and [$^{18}$F]FSK in *S. aureus* and *K. pneumoniae* at 30 min, 60 min and 90 min.

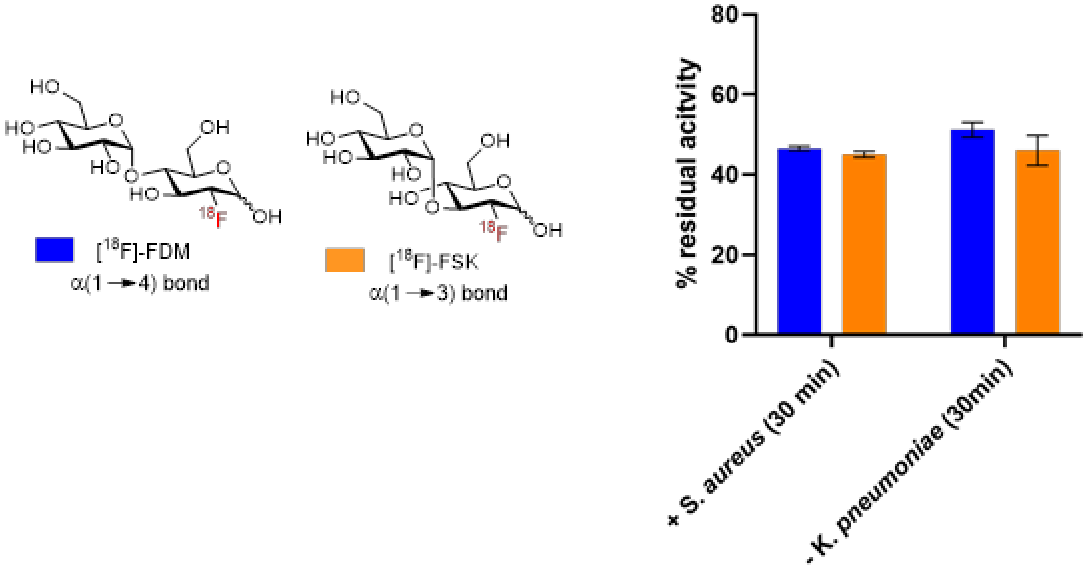

**Figure S3.** Efflux experiment: plot showing the residual activity observed at 30 min post efflux of [$^{18}$F]FDM and [$^{18}$F]FSK in *S. aureus* and *K. pneumoniae* following an initial 30 minutes incubation.

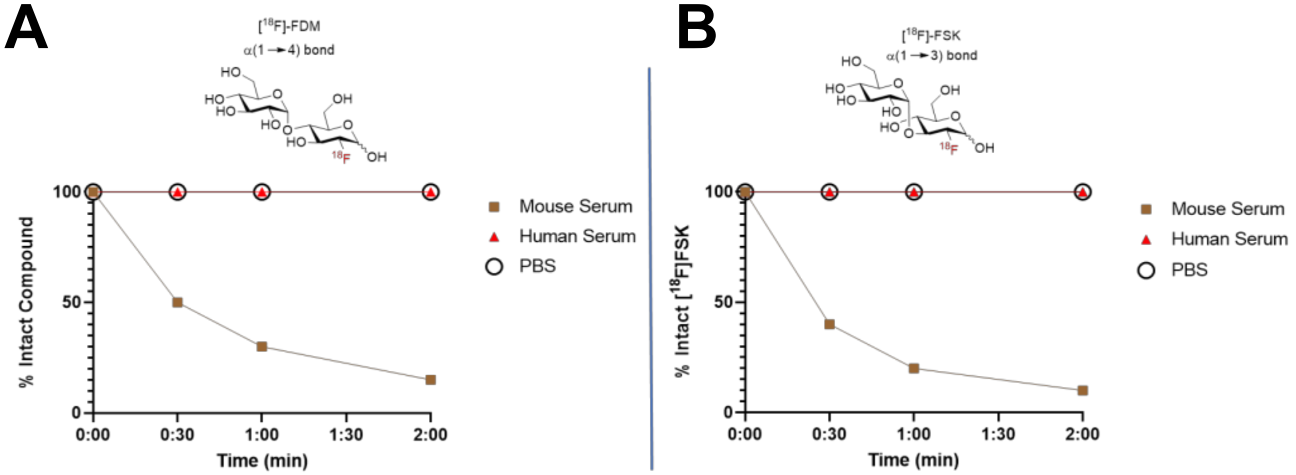

**Figure S4.** Stability of [$^{18}$F]FDM and [$^{18}$F]FSK in human and mouse serum.

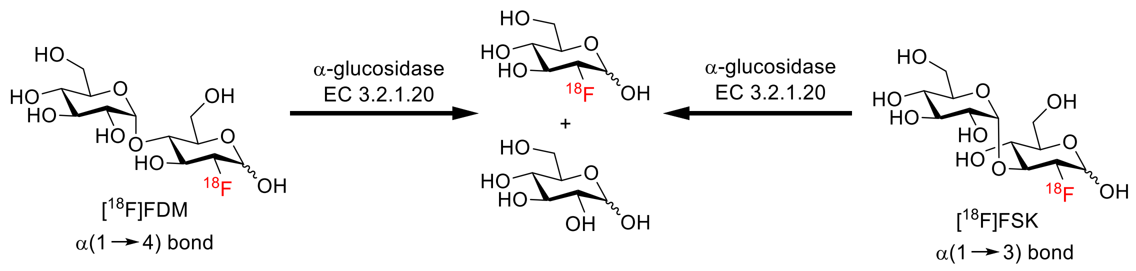

**Figure S5.** Schematic of α-glucosidase hydrolysis of [$^{18}$F]FDM and [$^{18}$F]FSK. In murine serum, α-glucosidase catalyzes the decomposition of [$^{18}$F]FDM and [$^{18}$F]FSK into [$^{18}$F]FDG and glucose.

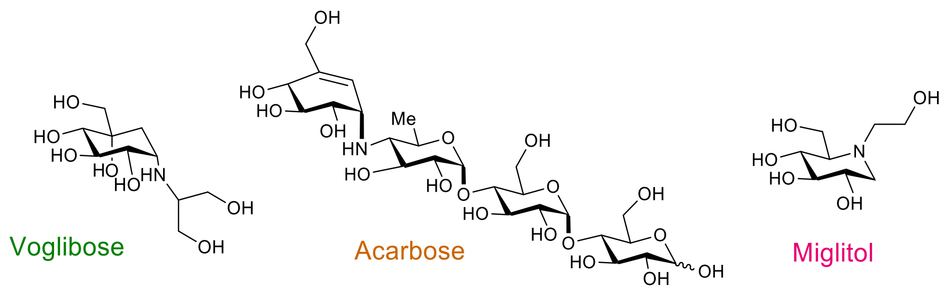

**Figure S6.** Inhibitors of α–glycosidase that have the potential for higher stability of [$^{18}$F]FDM and [$^{18}$F]FSK in mouse serum.

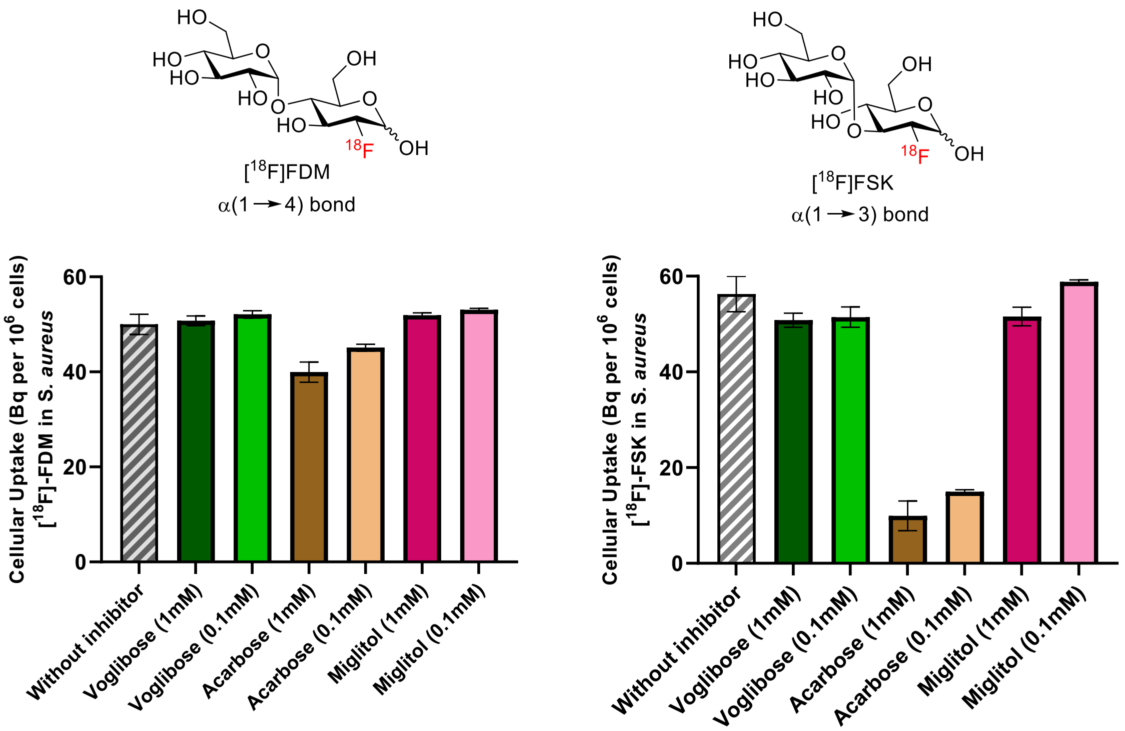

**Figure S7.** *In vitro S. aureus* uptake for [$^{18}$F]FDM and [$^{18}$F]FSK in the presence of the indicated α-glucosidase inhibitors.

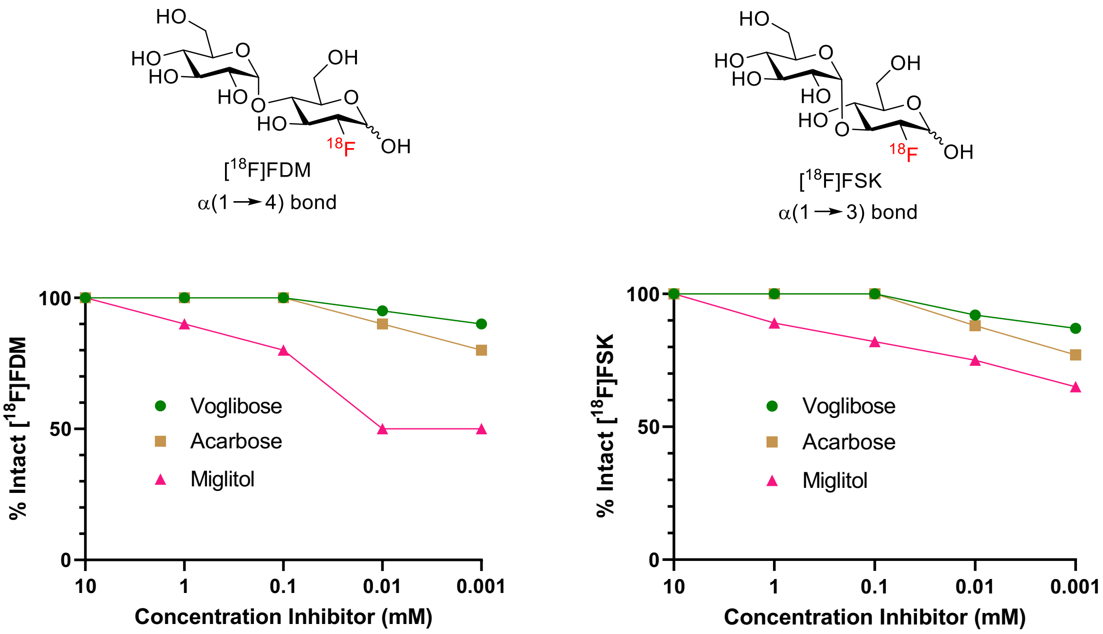

**Figure S8.** Dose response of [$^{18}$F]FDM and [$^{18}$F]FSK in mouse serum exposed to increasing concentrations of α-glucosidase inhibitors (0.001 to 10 mM).

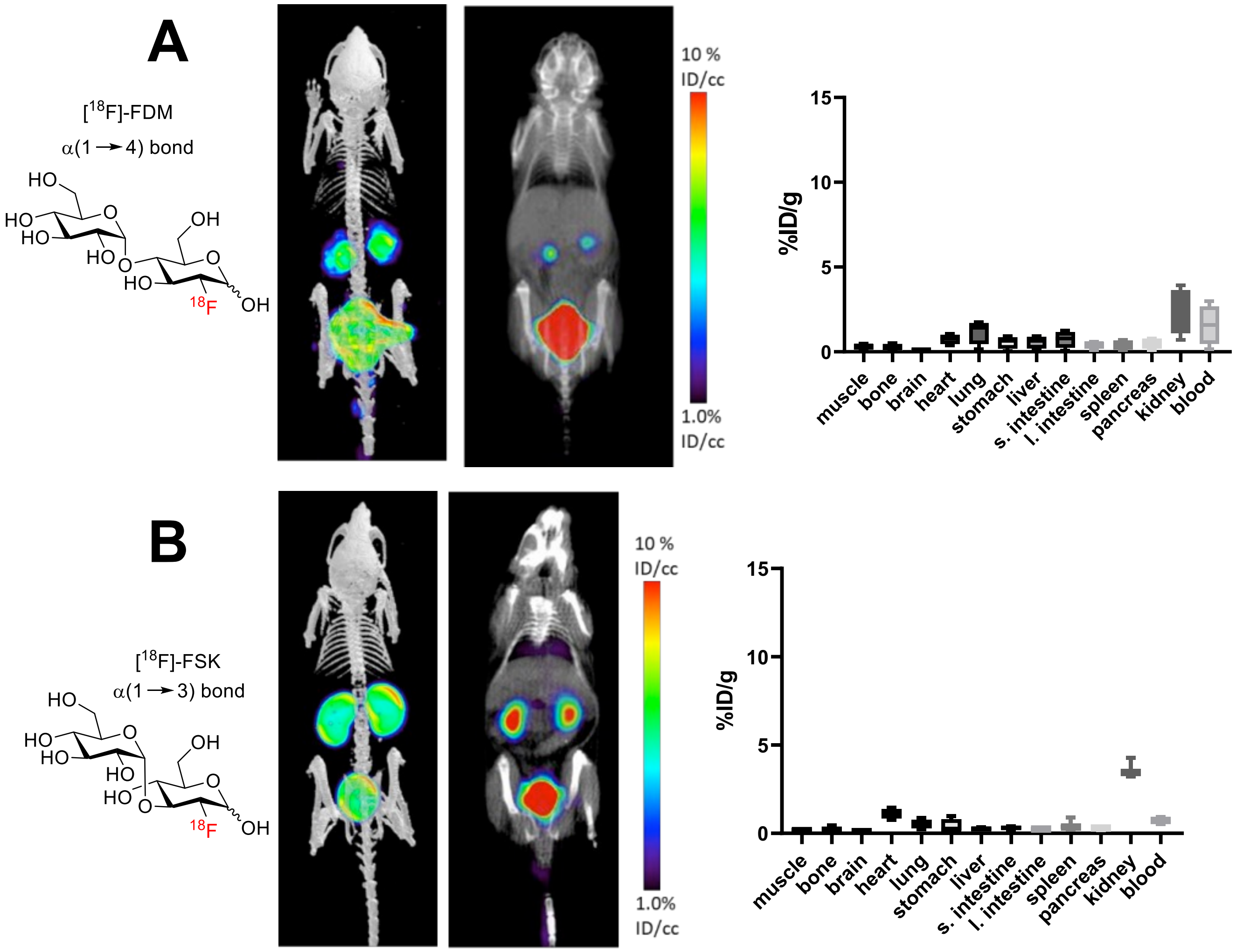

**Figure S9.** *In vivo* experiment: [$^{18}$F]FDM and [$^{18}$F]FSK in mouse, No bacteria, Inhibitor: Voglibose (1mg/ inj), Injection: ≈200uCi 18F-tracer, N = 5. *Ex vivo* data was obtained following tissue harvesting on a gamma counter.

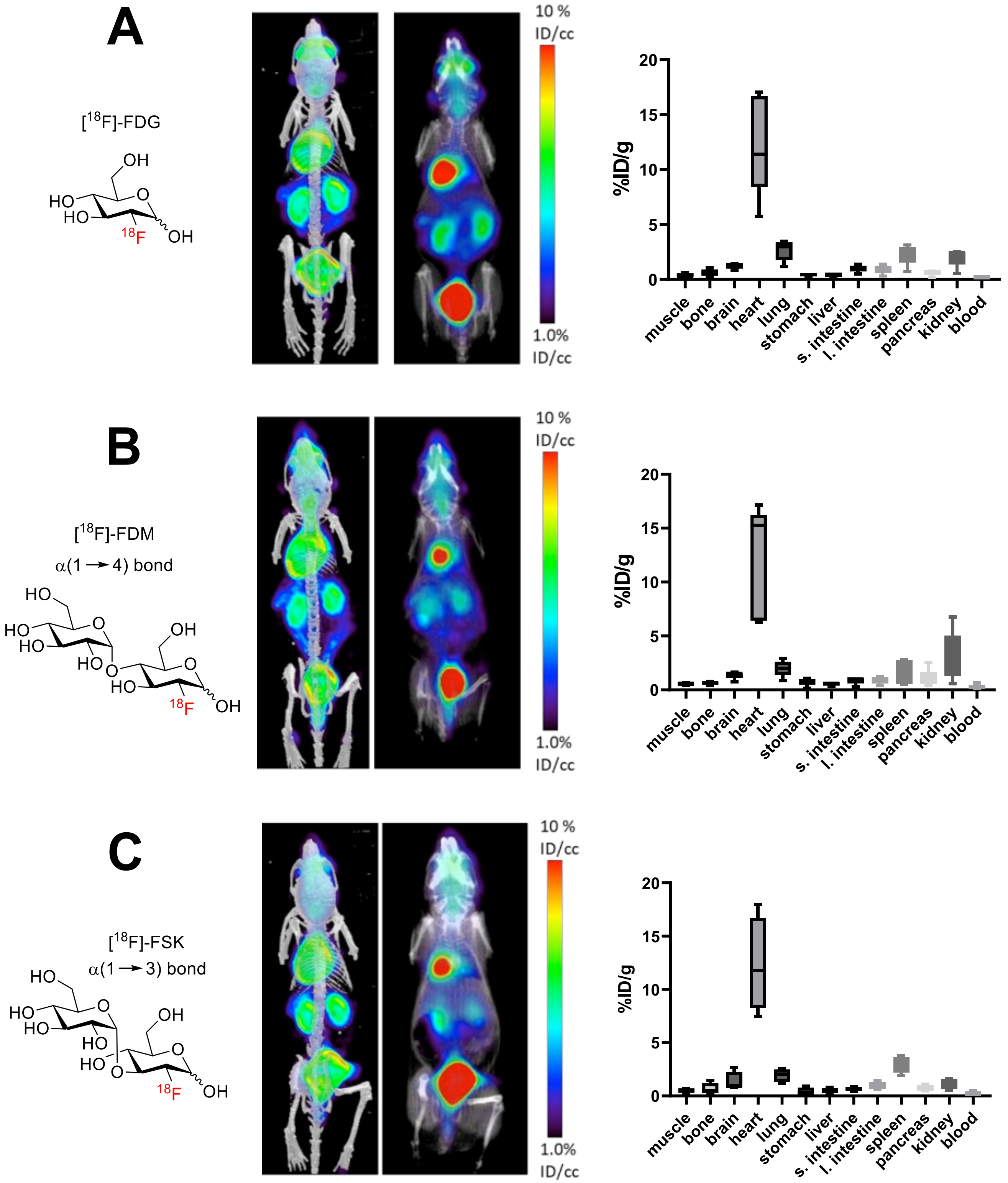

**Figure S10.** *In vivo* experiment: [$^{18}$F]FDG, [$^{18}$F]FDM and [$^{18}$F]FSK in mouse, No bacteria, No inhibitor, Injection: ≈200uCi 18F-tracer, N = 5. *Ex vivo* data was obtained following tissue harvesting on a gamma counter.

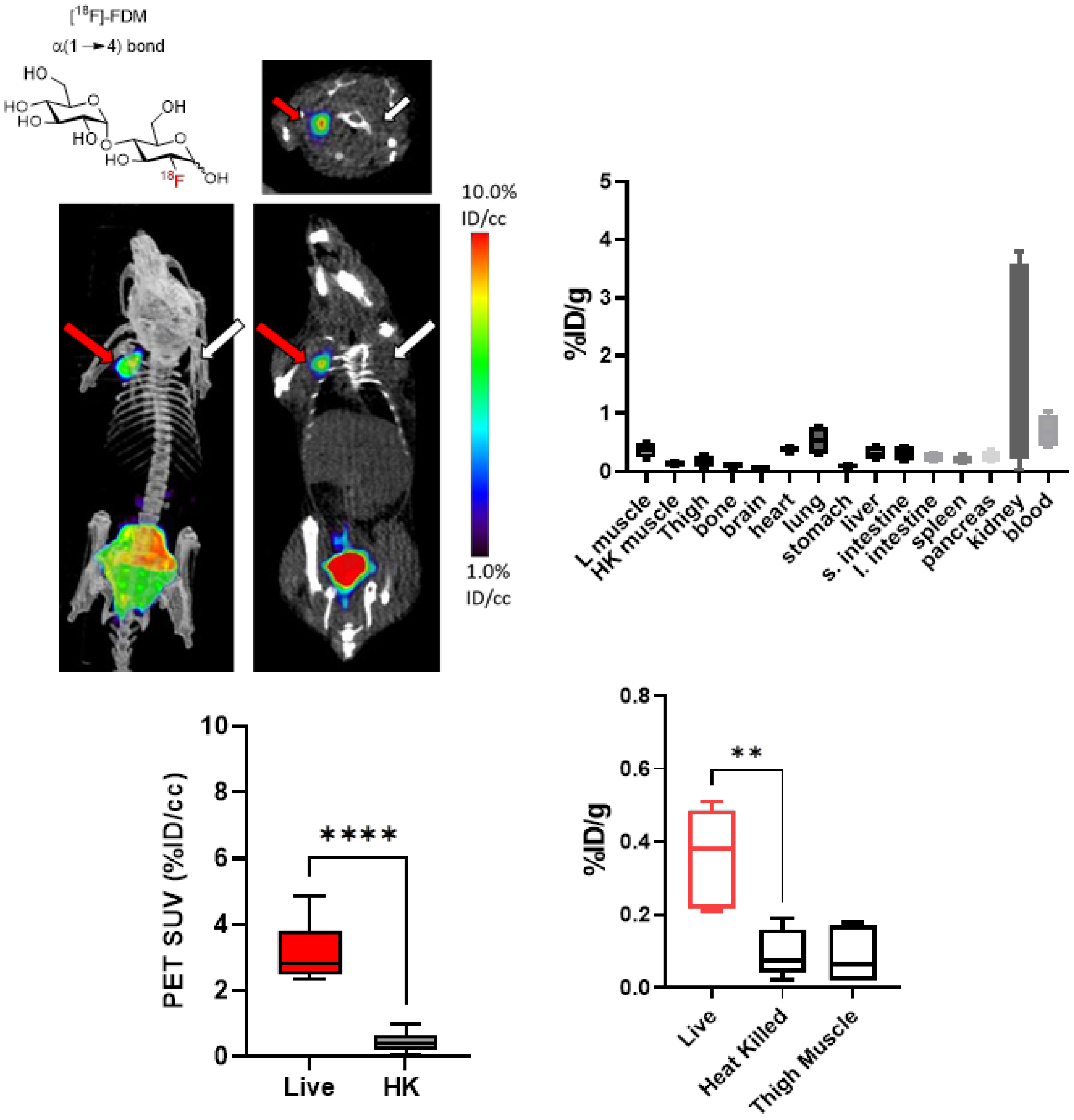

**Figure S11.** *In vivo* experiment: [18F]FDM in mouse myositis model, *S. aureus* (MRSA 01), Inhibitor: Voglibose (1mg/ inj), Injection: ≈200uCi 18F-tracer, N = 6. The red arrows indicate the site of inoculation with live bacteria, while white arrows indicate the site of inoculation with heat-killed bacteria. *Ex vivo* data was obtained following tissue harvesting on a gamma counter. ROI analysis, *live vs HK*: 6.1 fold excess, **** *P* value <0.0001, *ex vivo* analysis, *live vs HK*: 3.9 fold excess, ** *P* value = 0.001 (unpaired t-test).

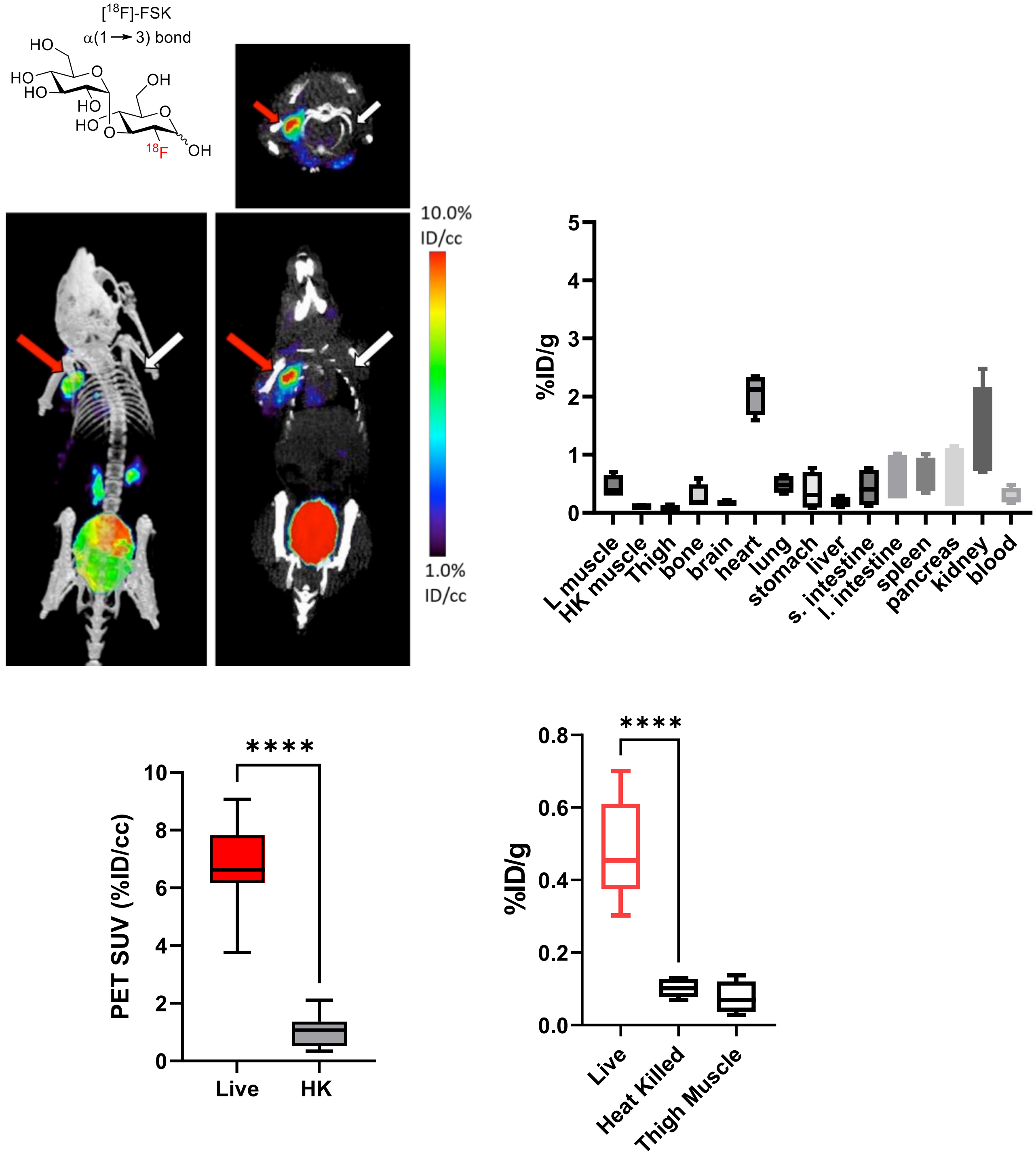

**Figure S12.** *In vivo* experiment: [18F]FSK in mouse myositis model, *S. aureus* (MRSA 01), Inhibitor: Voglibose (1mg/ inj), Injection: ≈200uCi 18F-tracer, N = 6. The red arrows indicate the site of inoculation with live bacteria, while white arrows indicate the site of inoculation with heat-killed bacteria. *Ex vivo* data was obtained following tissue harvesting on a gamma counter. ROI analysis, *live vs HK*: 6.5 fold excess, **** *P* value <0.0001, *ex vivo* analysis, *live vs HK*: 4.7 fold excess, **** *P* value < 0.0001 (unpaired t-test).

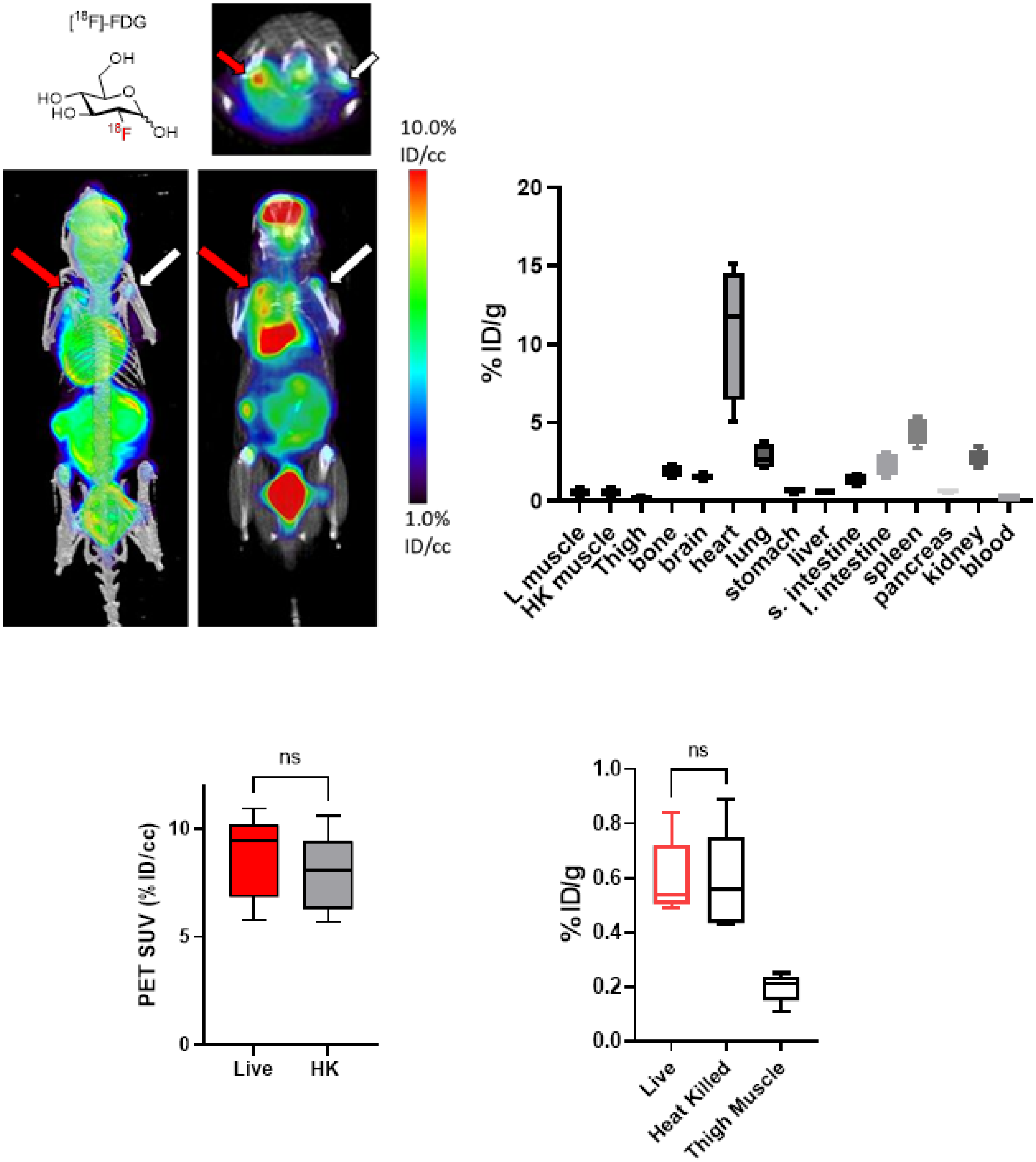

**Figure S13.** *In vivo* experiment: [$^{18}$F]FDG in mouse myositis model, *S. aureus* (MRSA 01), Inhibitor: Voglibose (1mg/ inj), Injection: ≈200uCi 18F-tracer, N = 5. The red arrows indicate the site of inoculation with live bacteria, while white arrows indicate the site of inoculation with heat-killed bacteria. *Ex vivo* data was obtained following tissue harvesting on a gamma counter. ROI analysis, *live vs HK*: 1.10 fold excess, ns, *ex vivo* analysis, *live vs HK*: 1.02 fold excess, ns (unpaired t-test).

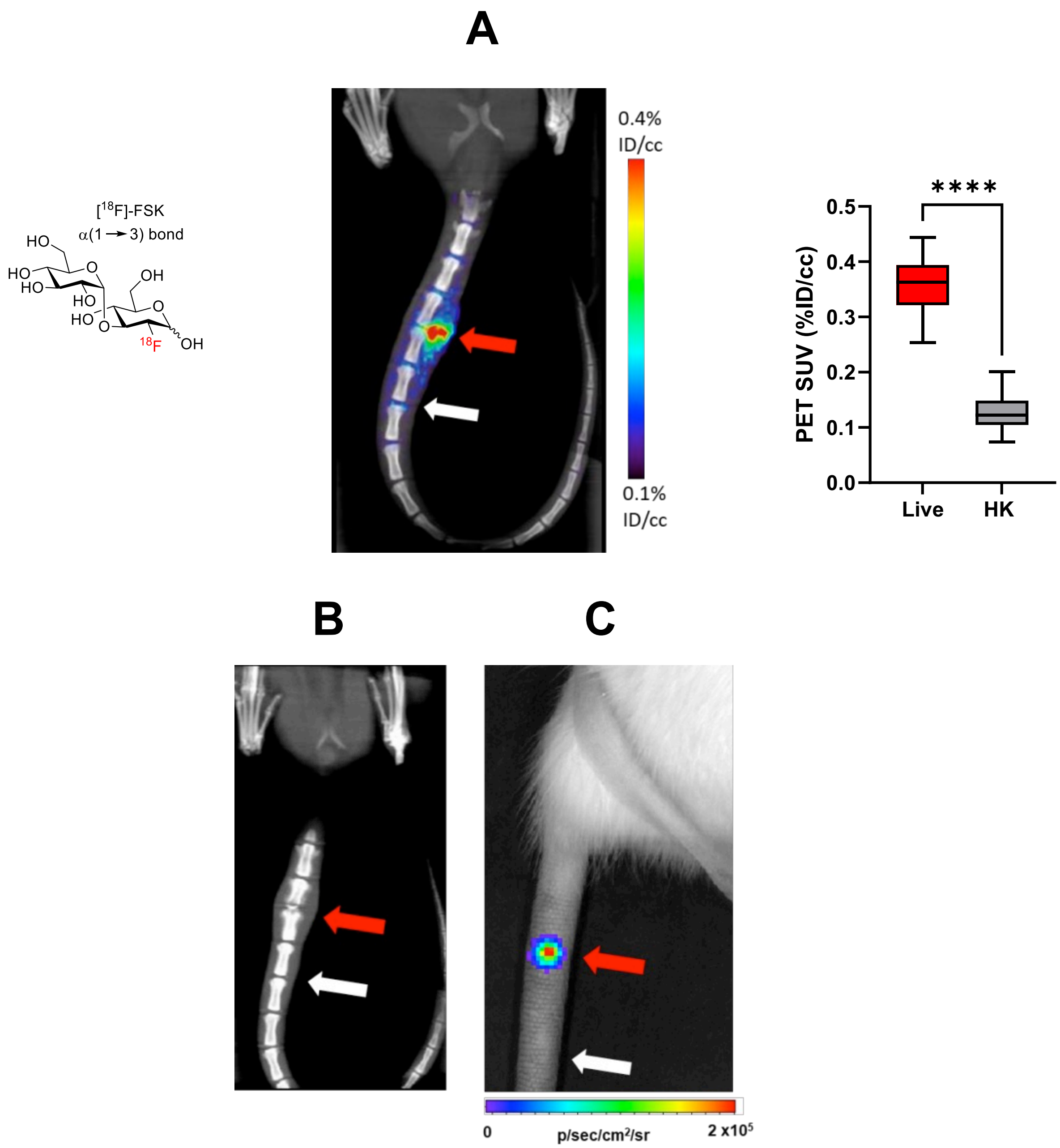

**Figure S14** *In vivo* experiment: [18F]FSK in rat vertebral discitis-osteomyelitis (VDO), *S. aureus* (Xen29), Day 4, inhibitor: Voglibose (5mg/ inj), Injection: ≈500uCi 18F-tracer, N = 5. The red arrows indicate the site of inoculation with live bacteria, while white arrows indicate the site of inoculation with heat-killed bacteria. (A) PET/CT imaging of *S. aureus* Xen29 vertebral discitis-osteomyelitis (VDO) in rat (*N* = 5) with [18F]FSK. ROI analysis, *live vs HK*: 2.8 fold excess, **** *P* value <0.0001 (unpaired t-test). (B) Computed tomography study performed at 10 days highlights the similarity between rodent and human discitis osteomyelitis. (C) Optical tomography image of rat tail showing bioluminescent signal from *S. aureus* Xen29 inoculation.

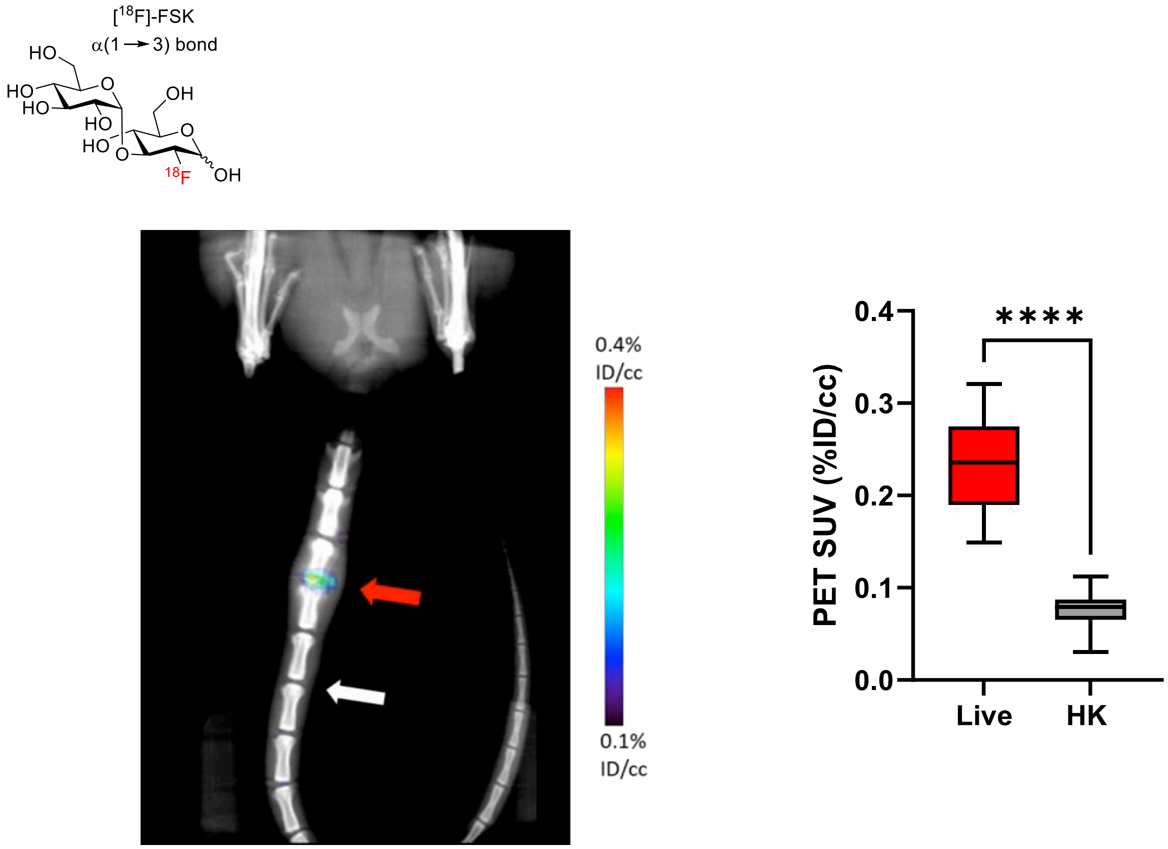

**Figure S15.** *In vivo* experiment: [$^{18}$F]FSK in rat vertebral discitis-osteomyelitis (VDO), *S. aureus* (Xen29), Day 10, inhibitor: Voglibose (5mg/ inj), Injection: ≈500uCi 18F-tracer, N = 3. The red arrows indicate the site of inoculation with live bacteria, while white arrows indicate the site of inoculation with heat-killed bacteria. ROI analysis, *live vs HK*: 3.1 fold excess, **** *P* value <0.0001 (unpaired t-test).

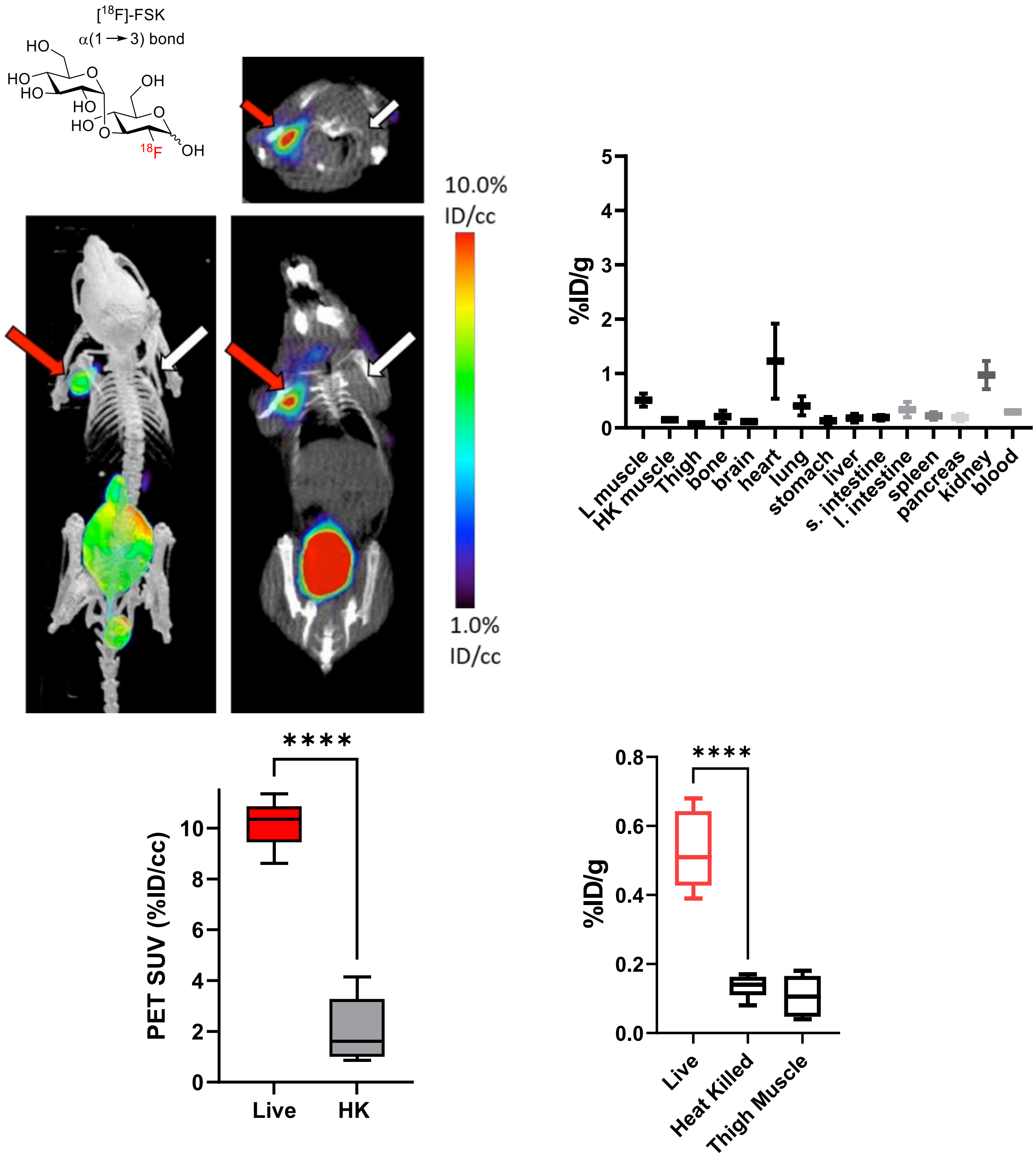

**Figure S16.** *In vivo* experiment: [$^{18}$F]FSK in mouse myositis model, *A. baumanni* (ATCC 19606), inhibitor: Voglibose (1mg/ inj), Injection: ≈200uCi 18F-tracer, N = 6. The red arrows indicate the site of inoculation with live bacteria, while white arrows indicate the site of inoculation with heat-killed bacteria. *Ex vivo* data was obtained following tissue harvesting on a gamma counter. ROI analysis, *live vs HK*: 4.9 fold excess, **** *P* value <0.0001, *ex vivo* analysis, *live vs HK*: 3.9 fold excess, **** *P* value < 0.0001 (unpaired t-test).

**Table S1. Bacterial strains**

The bacterial strains included in this study are listed in the table below.

| Strain | Phenotype or Genotype | Source or Reference |
|---|---|---|
| *S. aureus* | Wild-type | ATCC 12600 |
| *S. aureus* Xen29 | ATCC 12600 expressing the *Photorhabdus luminescens luxABCDE* genes | Xenogen USA |
| *S. aureus* MRSA 1 | MRSA | Clinical isolate, University of Nebraska Medical Center |
| *S. aureus* MRSA 2 | MRSA | Clinical isolate, University of Nebraska Medical Center |
| *S. aureus* MRSA 3 | MRSA | Clinical isolate, University of Nebraska Medical Center |
| *S. aureus* MRSA 4 | MRSA | Clinical isolate, University of Nebraska Medical Center |
| *L. monocytogenes* | Wild-type | ATCC 15313 |
| *S. epidermidis* | Wild-type | ATCC 35984 |
| *K. pneumoniae* | Wild-type | ATCC 13883 |
| *E. coli* | Wild-type | ATCC 25922 |
| *P. aeruginosa* PA01 | Wild-type | ATCC 10154 |
| *A. baumannii* | Wild-type | ATCC 19606 |
| *S. typhimurium* | Wild-type | ATCC 29630 |
| *P. mirabilis* | Wild-type | ATCC 29906 |
| *E. cloacae* | Wild-type | ATCC 7256 |
| *E. faecalis* | Wild-type | ATCC 19433 |

### B. Synthetic Procedures

#### B.1. General:

All chemical reagents were purchased from commercial sources (Acros Organics, Alfa Aesar, AK Scientific & Sigma-Aldrich) and used without further purification unless otherwise stated. All separatory cartridges were purchased from Waters. Reactions were monitored by thin layer chromatography (TLC) on precoated (250 μm) silica gel 60 F254 aluminum sheets and visualized under a UV-254 lamp followed by staining with potassium permanganate. Flash chromatography was performed on silica gel (60A pore size). $^{1}H$, $^{13}C$, $^{31}P$ and $^{19}F$ NMR spectra were obtained on a Bruker Avance III HD 400 mHz instrument at the UCSF Nuclear Magnetic Resonance Laboratory and data were processed using MestReNova. Abbreviations are as follows: s (singlet), d (doublet), t (triplet), q (quartet), m (multiplet). High resolution mass spectra (HRMS) services were provided by University of California, Berkeley Spectrometry Facility. The $^{18}F$ labeled compounds were characterized by developing the compounds in different solvent systems on silica gel TLC plates on glass followed by imaging on a radio TLC scanner (Bioscan AR2000). Analytical HPLC was performed using a Waters pump equipped with a manual Rheodyne injector (1 mL loop), a refracted index (RI) detector and a RAD detector. The stationary phase was YMC-Pack Polyamine II column and the mobile phase of 73:27 acetonitrile/$H_2O$ at a flowrate of 1 mL/min. For semi prep HPLC, a YMC-Pack Polyamine II stationary phase was used with a mobile phase of 73:27 acetonitrile/$H_2O$ at a flowrate of 4 mL/min, The radioactivity of the bacterial pallets and filtrate were counted on a γ counter (Hidex Automatic Gamma Counter).

#### B.2. Synthesis of precursor: β-D-glucose-1-phosphate (βGlc1-P)

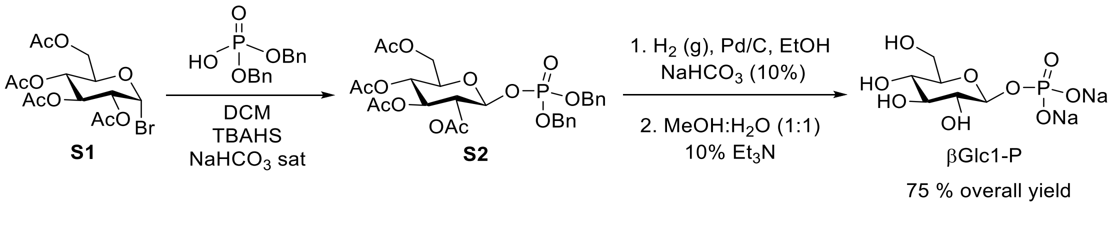

Dibenzyl(2,3,4,6-tetra-*O*-acetyl-β-D-glucopyranosyl) phosphate (**S2**).

In a 250 mL Erlenmeyer flask, dibenzyl phosphate (6.8 g, 24.4 mmol) and tetrabutylammonium hydrogen sulfate (4.1 g, 12.2 mmol) were added and mixed with 120 mL of saturated $NaHCO_3$ aqueous solution. After stirring at room temperature for 10 min, the mixture was added to a 500 mL round bottom flask containing bromide **S1** (5.0 g, 12.2 mmol) in DCM (120 mL). The biphasic reaction mixture was capped then stirred vigorously at room temperature for 72 hrs. Over the reaction time, the pH of the aqueous phase was maintained at pH 8-9 via addition of saturated $NaHCO_3$ solution. The organic phase was extracted using ethyl acetate, before being washed with saturated $NaHCO_3$, water, and brine. The organic phase was then dried with sodium sulfate, filtered, and evaporated under reduced pressure. The residue was purified via silica flash chromatography using 1% *t*-butanol in DCM (with a few drops of $Et_3N$) as eluant to give pure phosphate **S2** (5.2 g, 71% yield).

$^1$H NMR (400 MHz, $CDCl_3$) δ 7.46 – 7.30 (m, 10H), 5.38 (t, *J* = 7.6 Hz, 1H), 5.23 (d, *J* = 9.4 Hz, 1H), 5.16 (d, *J* = 7.8 Hz, 1H), 5.14 (d, *J* = 6.3 Hz, 1H), 5.11 (d, *J* = 7.4 Hz, 2H), 5.04 (d, *J* = 7.1 Hz, 2H), 4.26 (dd, *J* = 12.5, 4.8 Hz, 1H), 4.14 (dd, *J* = 12.4, 2.1 Hz, 1H), 3.83 (ddd, *J* = 10.0, 4.8, 2.2 Hz, 1H), 2.06 (s, 3H), 2.03 (s, 6H), 1.92 (s, 3H).

$^{31}$P NMR (162 MHz, $CDCl_3$) δ -3.20 (dd, *J* = 14.5, 7.3 Hz). The NMR values matched the reported literature.[1]

β-D-Glucose-1-phosphate (βGlc1-P).

In a 100 mL round bottom flask, phosphate **S2** (0.5 mg, 0.82 mmol) was hydrogenated (14.7 psi) over 5% Pd/C (94.3 mg), 14 mL of EtOH and 9 mL of 10% $NaHCO_3$ for 16 hrs at room temperature. The mixture was then filtered then concentrated. The residue was transferred to a 25 mL round bottom flask, to which was added $Et_3N$ (0.4 mL) and a 1:1 mixture of MeOH/$H_2O$ (5.5 mL). The mixture was stirred for 16 hrs before being concentrated then diluted with $H_2O$. The pH was lowered using Dowex 50W-X8 [$H^+$] resin until pH 7, before residue was filtered then lyophilized to obtain β-D-glucose-1-phosphate (βGlc1-P), (201 mg, 80 % yield).

$^1$H NMR (400 MHz, $D_2O$) δ 4.90 (t, *J* = 7.7 Hz, 1H), 3.91 (dd, *J* = 12.3, 2.1 Hz, 1H), 3.68 (dd, *J* = 12.3, 6.9 Hz, 1H), 3.56 – 3.47 (m, 2H), 3.34 (t, 1H), 3.31 (t, 1H).

$^{31}$P NMR (162 MHz, $D_2O$) δ 2.29 (d, *J* = 7.7 Hz). The NMR values matched the reported literature.[2]

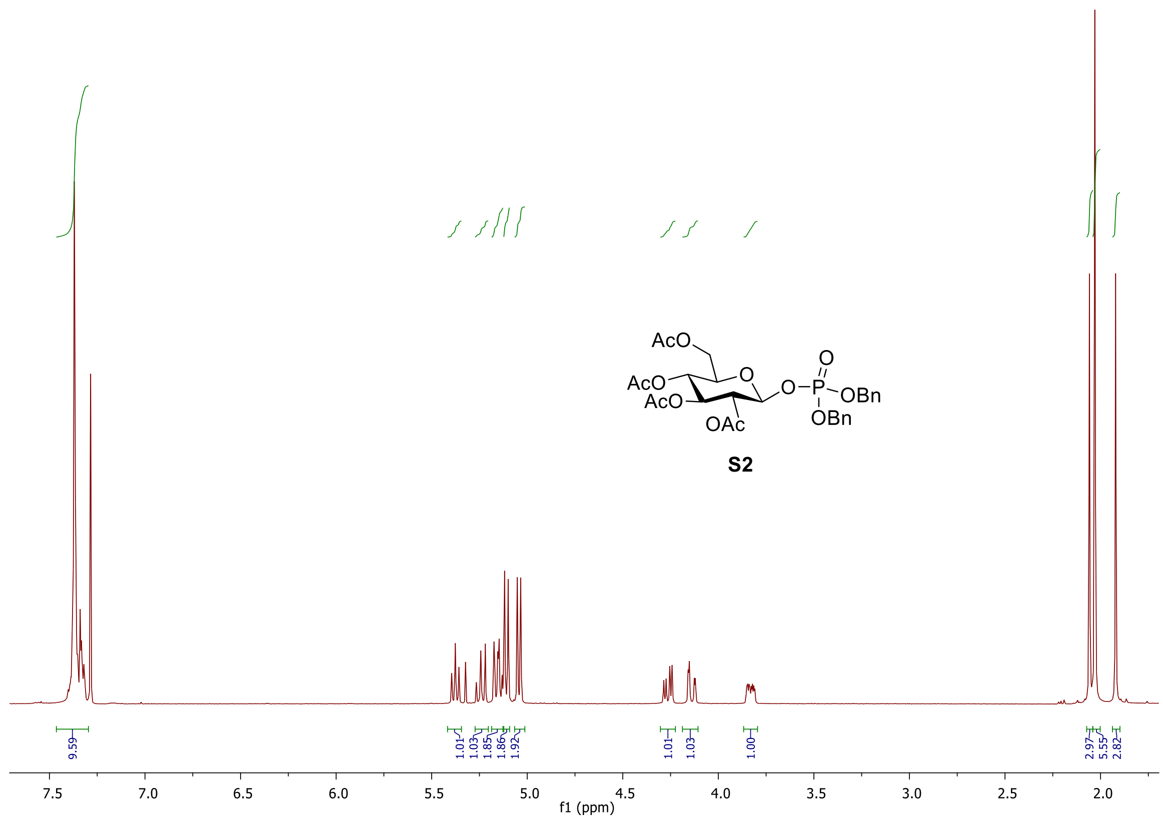

**Figure B.2.1.** $^{1}$H NMR of (**S2**) in $CDCl_3$.

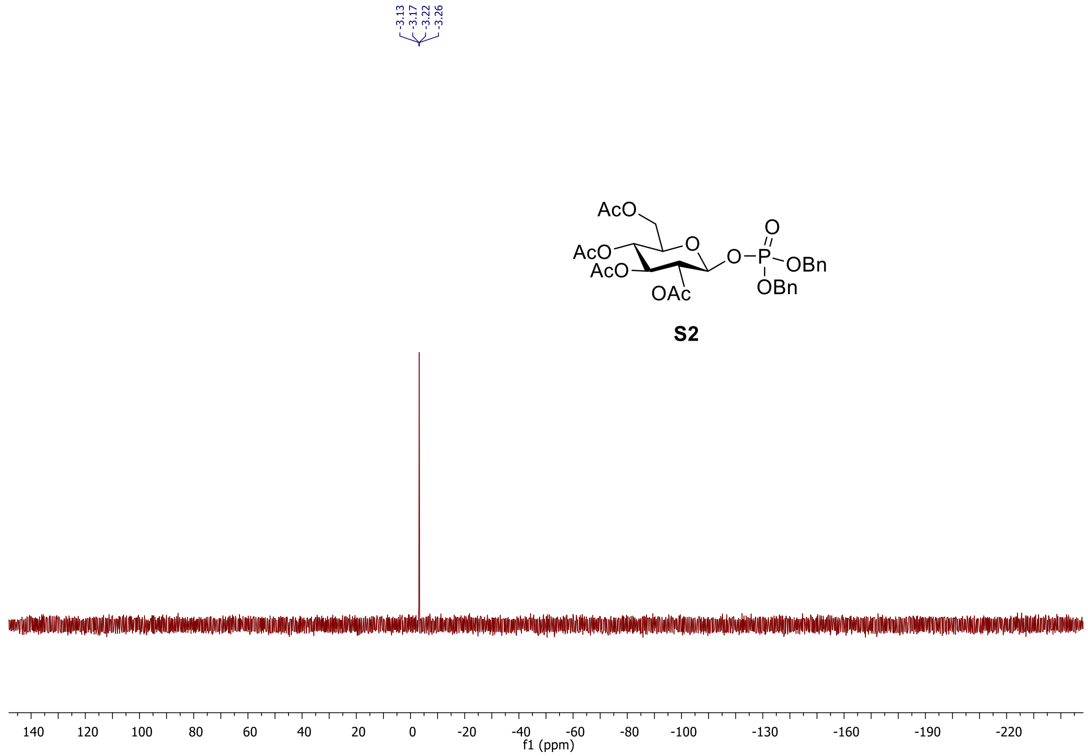

**Figure B.2.2.** $^{31}$P NMR of (**S2**) in $CDCl_3$.

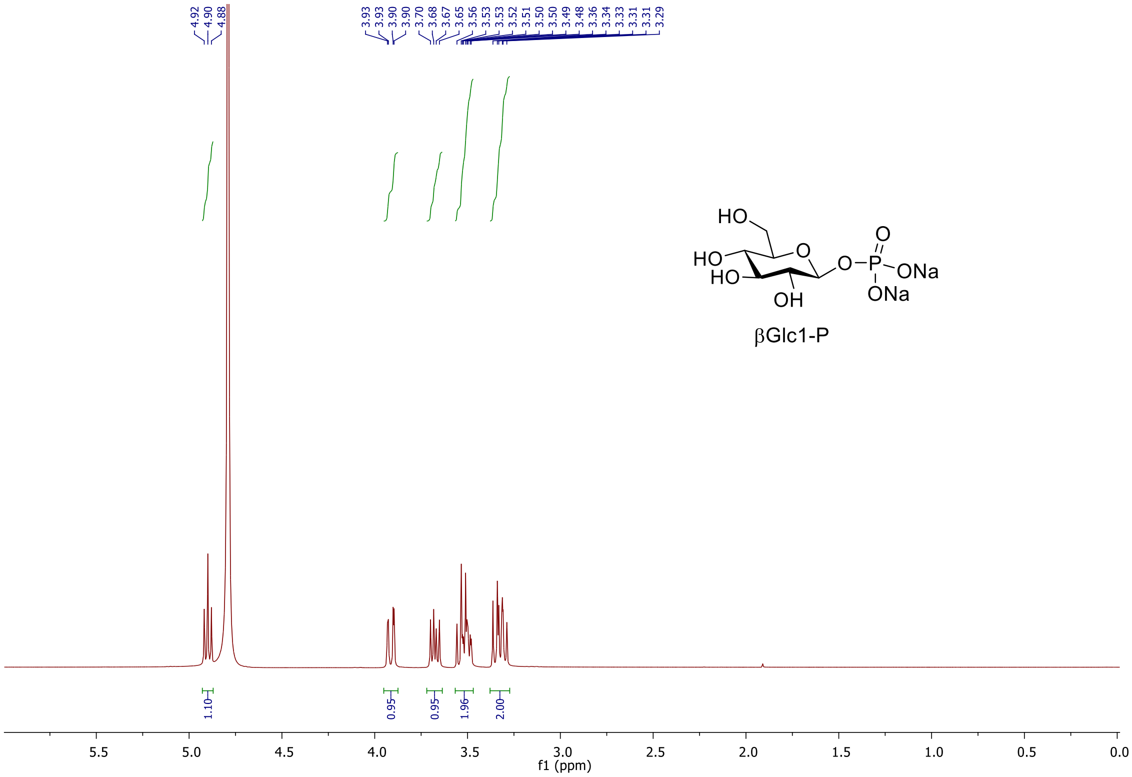

**Figure B.2.3.** $^{1}$H NMR of β-D-glucose-1-phosphate in $D_2O$.

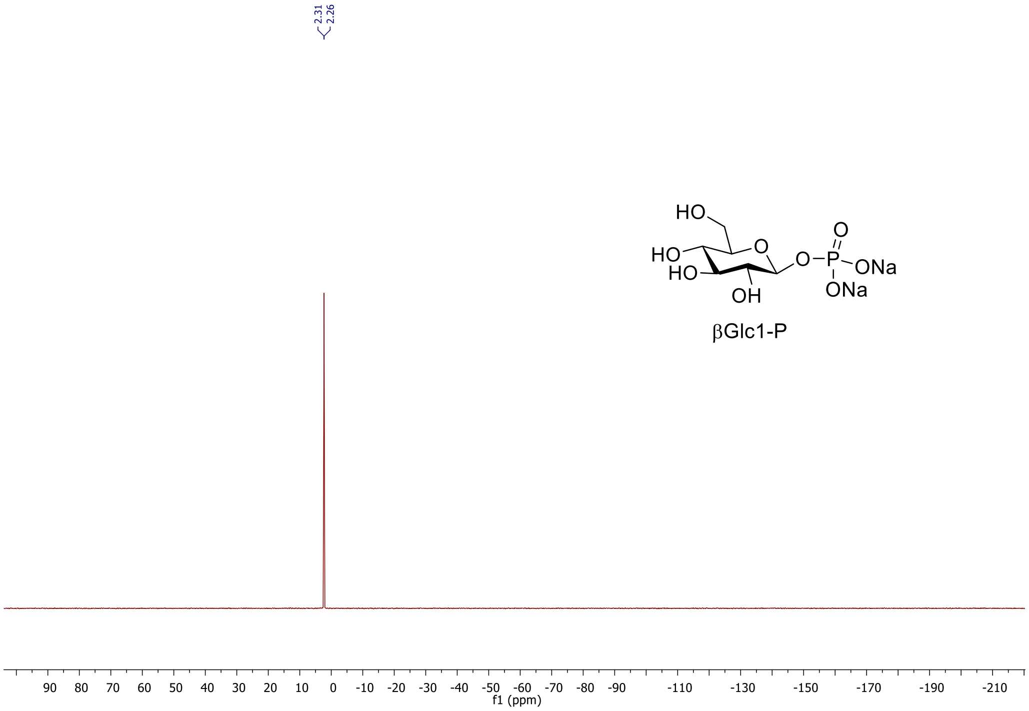

**Figure B.2.4.** $^{31}$P NMR of β-D-glucose-1-phosphate in $D_2O$.

### B.3. Synthesis of $^{19}$F Standards:

#### B.3.1. 2-deoxy-2-[$^{19}$F]fluoro-maltose and 2-deoxy-2-[$^{19}$F]fluoro-sakebiose

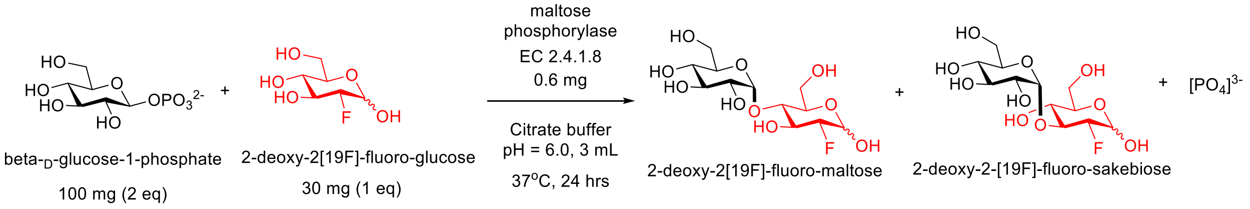

β-D-glucose-1-phosphate (βGlc1-P) (100 mg, 0.32 mmol), 2-deoxy-2[$^{19}$F]-fluoro-glucose (30 mg, 0.16 mmol) were added to a 10 mL round bottom flask containing maltose phosphorylase (EC 2.4.1.8, Sigma Aldrich), (0.6 mg, 6 units) in 3 mL of aqueous citrate buffer solution (pH = 6.0). The mixture was stirred for 24 hrs at 37°C. The residue was diluted with MeCN then purified via semi prep HPLC (YMC-Pack Polyamine II, 250 X 10 mm) using 70% MeCN/30 % $H_2O$ to yield 2-deoxy-2[$^{19}$F]-fluoro-maltose (22 mg, 41% yield) and 2-deoxy-2-[$^{19}$F]-fluoro-sakebiose (8 mg, 14% yield).

#### Semi-Prep HPLC Purification

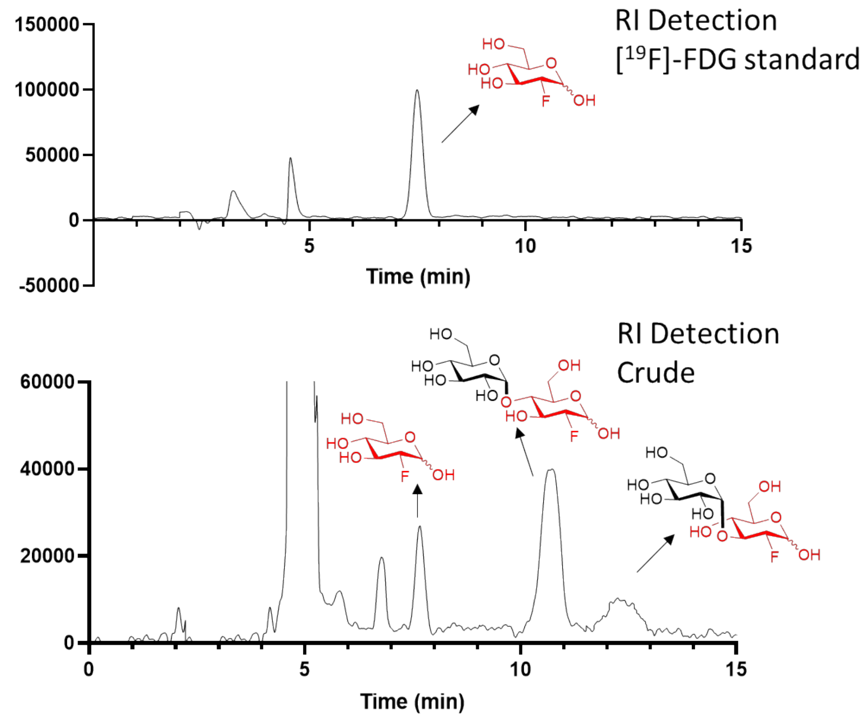

**Figure B.3.1.** Chromatograms for semi-prep HPLC purification of 2-deoxy-2[$^{19}$F]-fluoro-maltose and 2-deoxy-2-[$^{19}$F]-fluoro-sakebiose.

#### B.3.2. 2-deoxy-2-[$^{19}$F]fluoro-cellobiose

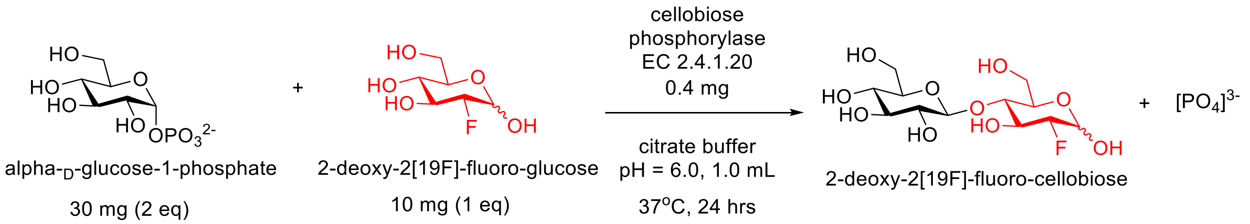

α-D-glucose-1-phosphate (αGlc1-P) (30 mg, 0.1 mmol), 2-deoxy-2[$^{19}$F]-fluoro-glucose (10 mg, 0.05 mmol) were added to a 5 mL round bottom flask containing cellobiose phosphorylase (EC 2.4.1.20), (0.4 mg, 6 units) in 1 mL of aqueous citrate buffer solution (pH = 6.0). The mixture was stirred for 24 hrs at 37°C. The residue was diluted with MeCN then purified via semi prep HPLC (YMC-Pack Polyamine II, 250 X 10 mm) using 70% MeCN/30 % $H_2O$ to yield 2-deoxy-2[$^{19}$F]-fluoro-cellobiose (10 mg, 59% yield).

#### Semi-Prep HPLC Purification

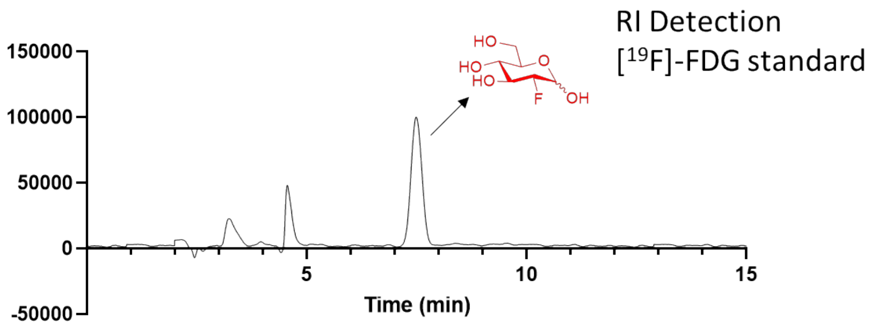

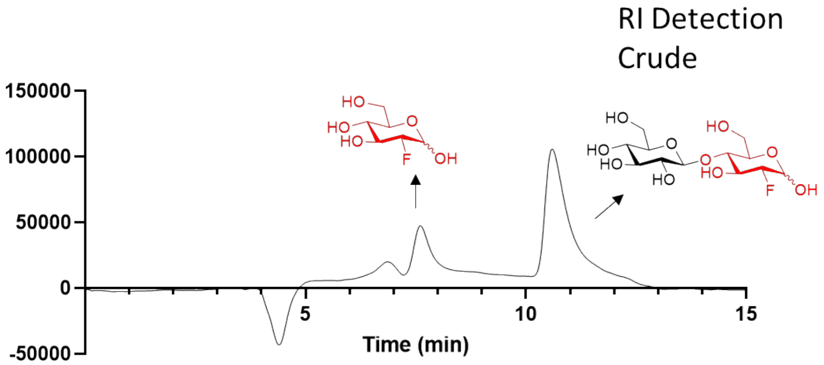

**Figure B.3.2** Chromatograms for semi-prep HPLC purification of 2-deoxy-2[$^{19}$F]-fluoro-cellobiose.

#### B.3.3. 2-deoxy-2-[$^{19}$F]fluoro-trehalose

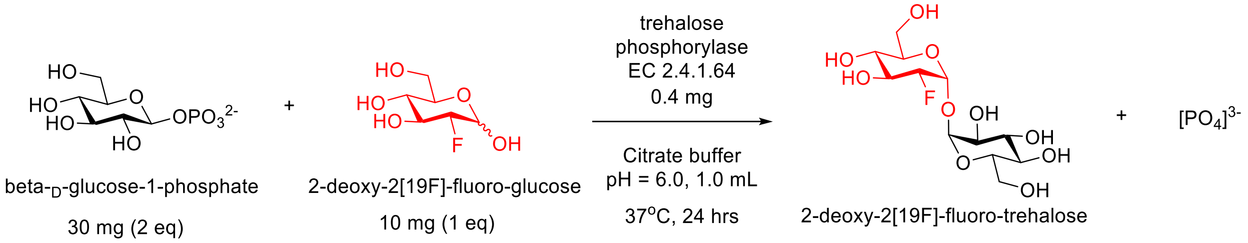

β-D-glucose-1-phosphate (βGlc1-P) (30 mg, 0.1 mmol), 2-deoxy-2[$^{19}$F]-fluoro-glucose (10 mg, 0.05 mmol) were added to a 5 mL round bottom flask containing trehalose phosphorylase (EC 2.4.1.64), (0.4 mg, 6 units) in 1 mL of aqueous citrate buffer solution (pH = 6.0). The mixture was stirred for 24 hrs at 37°C. The residue was diluted with MeCN then purified via semi prep HPLC (YMC-Pack Polyamine II, 250 X 10 mm) using 70% MeCN/30 % $H_2O$ to yield 2-deoxy-2[$^{19}$F]-fluoro-trehalose (8 mg, 47% yield).

#### Semi-Prep HPLC Purification

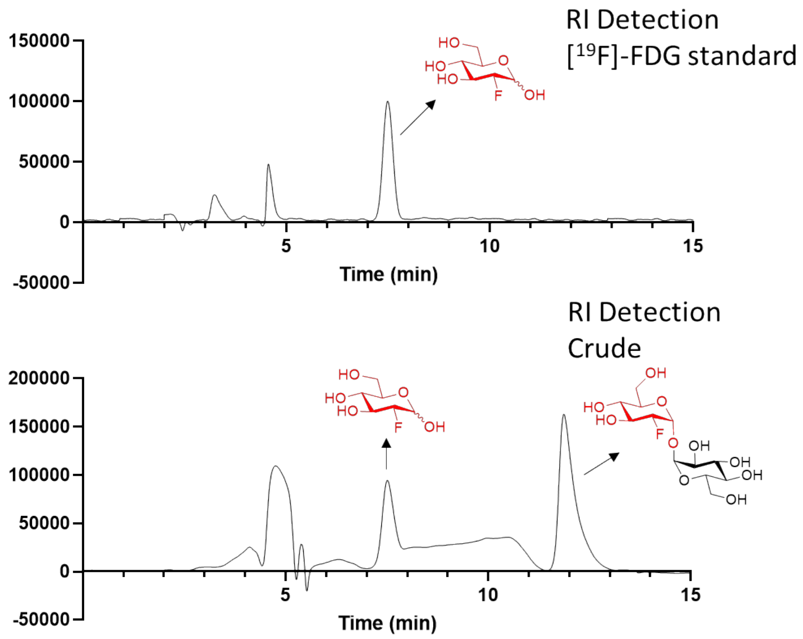

**Figure B.3.3.** Chromatograms for semi-prep HPLC purification of 2-deoxy-2[$^{19}$F]-fluoro-trehalose.

#### B.3.4. 2-deoxy-2-[$^{19}$F]fluoro-laminaribiose

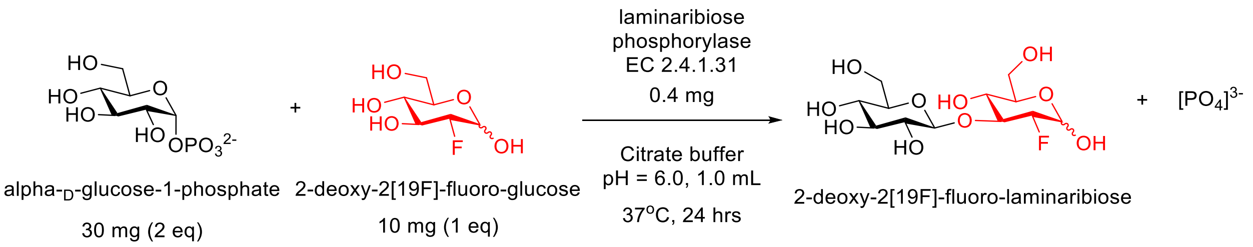

α-D-glucose-1-phosphate (αGlc1-P) (30 mg, 0.1 mmol), 2-deoxy-2[$^{19}$F]-fluoro-glucose (10 mg, 0.05 mmol) were added to a 5 mL round bottom flask containing laminaribiose phosphorylase (EC 2.4.1.31), (0.4 mg, 6 units) in 1 mL of aqueous citrate buffer solution (pH = 6.0). The mixture was stirred for 24 hrs at 37°C. The residue was diluted with MeCN then purified via semi prep HPLC (YMC-Pack Polyamine II, 250 X 10 mm) using 70% MeCN/30 % $H_2O$ to yield 2-deoxy-2[$^{19}$F]-fluoro-laminaribiose (5 mg, 29% yield).

**Semi-Prep HPLC Purification:**

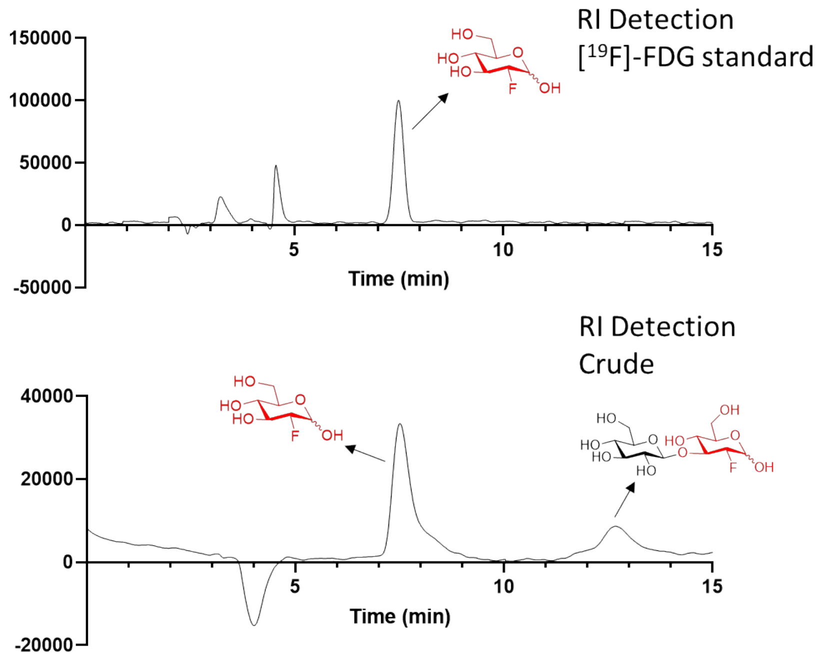

**Figure B.3.4.** Chromatograms for semi-prep HPLC purification of 2-deoxy-2[$^{19}$F]-fluoro-laminaribiose.

**NMR and HRMS analysis:**

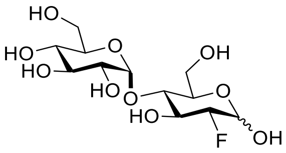

**2-deoxy-2[[19]F]-fluoro-maltose**

$^1$H NMR (400 MHz, $D_2O$) δ 5.39 – 5.30 (m, 3H), 4.83 (dd, *J* = 7.7, 2.3 Hz, 1H), 4.36 (ddd, *J* = 49.2, 9.6, 3.9 Hz, 1H), 4.20 – 3.96 (m, 3H), 3.89 (ddd, *J* = 10.0, 4.4, 2.4 Hz, 1H), 3.86 – 3.78 (m, 2H), 3.77 – 3.73 (m, 2H), 3.72 – 3.53 (m, 11H), 3.49 (ddd, *J* = 9.9, 3.9, 0.8 Hz, 2H), 3.34 (td, *J* = 9.5, 1.4 Hz, 2H).

$^{13}$C NMR (100 MHz, $D_2O$) δ 99.57 (C1'α), 99.41 (C1'β), 93.39 (d, *J* = 23.1 Hz, C1β), 92.67 (d, *J* = 183.6 Hz, C2β), 89.95 (d, *J* = 186.0 Hz, C2α), 89.43 (d, *J* = 21.3 Hz, C1α), 75.96 (d, *J* = 14.6 Hz, C4α), 75.88 (d, *J* = 14.9 Hz, C4β), 74.58 (d, *J* = 6.5 Hz, C5β), 74.49 (d, *J* = 24.2 Hz, C3β), 72.77 (C5'α), 72.74 (C5'β), 72.67 (C3'α), 72.66 (C3'β), 71.66 (C2'α), 71.57 (d, *J* = 18.2 Hz, C3α), 71.56 (C2'β), 69.71(C5α), 69.26 (C4'α, C4'β), 60.49 - 60.31 (C6α, C6'α, C6β, C6'β).

$^{19}$F NMR (376 MHz, $D_2O$) δ -199.90 – -200.12 (m), -200.24 (dd, *J* = 49.2, 13.7 Hz).

The NMR values matched the reported literature.[3]

HRMS (ESI) m/z calculated for $C_{12}H_{21}FO_{10}$ (M+Na) 367.1, found 367.1.

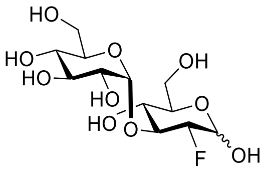

**2-deoxy-2-[[19]F]-fluoro-sakebiose**

$^1$H NMR (400 MHz, $D_2O$) δ 5.37 (d, *J* = 3.9 Hz, 1H), 5.23 (t, *J* = 4.3 Hz, 2H), 4.85 (dd, *J* = 7.9, 2.7 Hz, 1H), 4.44 (ddd, *J* = 48.9, 9.4, 3.9 Hz, 1H), 4.13 (dt, *J* = 53.2, 9.6 Hz, 1H), 4.00 (dt, *J* = 13.1, 10.5 Hz, 1H), 3.87 (m, 1H), 3.83 – 3.57 (m, 15H), 3.46 (m, 3H), 3.35 (t, *J* = 9.4 Hz, 2H).

$^{13}$C NMR (100 MHz, $D_2O$) δ 99.15 (C1'α), 99.09 (C1'β), 93.57 (d, *J* = 23.7 Hz, C1β), 91.43 (d, *J* = 185.7 Hz, C2β), 89.64 (d, *J* = 21.7 Hz, C1α), 88.71 (d, *J* = 188.2 Hz, C2α), 79.65 (d, *J* = 17.0 Hz, C3β), 77.06 (d, *J* = 17.0 Hz, C3α), 75.75 (C5β), 72.84 (C3'α), 72.83 (C3'β), 71.75 (C5'α), 71.71

(C5'β), 71.51 (C2'α), 71.46 (C2'β), 71.05 (C5α), 69.71 (d, *J* = 8.0 Hz, C4α), 69.53 (d, *J* = 7.7 Hz, C4β), 69.31 (C4'α +C4'β), 60.34 - 60.10 (C6α, C6'α, C6β, C6'β).

$^{19}$F NMR (376 MHz, $D_2O$) δ -196.49 (dd, *J* = 50.8, 15.1 Hz), -196.72 (dd, *J* = 48.9, 13.1 Hz).

HRMS (ESI) m/z calculated for $C_{12}H_{21}FO_{10}$ (M+Na) 367.1, found 367.1.

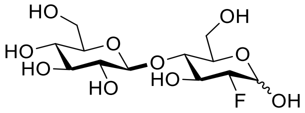

**2-deoxy-2[$^{19}$F]-fluoro-cellobiose**

$^{1}$H NMR (400 MHz, $D_2O$) δ 5.45 (d, *J* = 3.9 Hz, 1H), 4.93 (dd, *J* = 7.8, 2.4 Hz, 1H), 4.54 (d, *J* = 7.8 Hz, 2H), 4.48 (ddd, *J* = 49.5, 9.5, 3.9 Hz, 1H), 4.24 – 4.08 (m, 2H), 4.02 – 3.86 (m, 7H), 3.86 – 3.78 (dd, *J* = 12.4, 5.0 Hz, 1H), 3.78 – 3.69 (m, 4 H), 3.65 (ddd, *J* = 9.8, 4.9, 2.1 Hz, 1H), 3.58 – 3.38 (m, 6H), 3.33 (m, 2H).

$^{13}$C NMR (100 MHz, $D_2O$) δ 102.45 (C1'α+ C1'β), 93.45 (C1β), 93.22 (C2β), 89.90 (C2α), 89.32 (C1α), 77.93 (C4α+C4β), 75.94 (C5'α+C5'β), 75.42 (C3'α+C3'β), 74.86 (C5β), 73.07 (C2'α+C2'β), 72.66 (C3β), 69.92 (C5α), 69.54 (C3α), 69.44 (C4'α+C4'β), 60.56 (C6'α+C6'β), 59.80 (C6β), 59.63 (C6α).

$^{19}$F NMR (376 MHz, $D_2O$) δ -199.14 (ddd, *J* = 51.2, 15.2, 2.3 Hz), -199.33 (dd, *J* = 49.3, 13.5 Hz). The NMR values matched the reported literature.[4]

HRMS (ESI) m/z calculated for $C_{12}H_{21}FO_{10}$ (M+Na) 367.1, found 367.1.

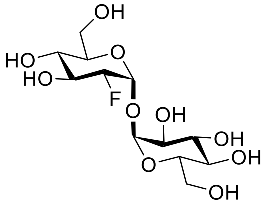

**2-deoxy-2[$^{19}$F]-fluoro-trehalose**

$^{1}$H NMR (400 MHz, D2O) δ 5.44 (d, *J* = 3.9 Hz, 1H), 5.22 (d, *J* = 3.7 Hz, 1H), 4.51 (ddd, *J* = 49.0, 9.6, 3.9 Hz, 1H), 4.13 (dt, *J* = 13.2, 9.4 Hz, 1H), 3.95 – 3.72 (m, 7H), 3.67 (dd, *J* = 9.9, 3.7 Hz, 1H), 3.52 (t, *J* = 9.6 Hz, 1H), 3.46 (t, *J* = 9.4 Hz, 1H).

$^{13}C$ NMR (100 MHz, $D_2O$) δ 94.00 (C1'), 91.19 (d, *J* = 21.6 Hz, C1), 89.47 (d, *J* = 187.7 Hz, C2), 72.58 (C3'), 72.19 (C5+C5'), 71.08 (d, *J* = 17.4 Hz, C3), 70.93 (C2'), 69.55 (C4'), 69.02 (d, *J* = 8.0 Hz, C4), 60.46 (C6), 60.26 (C6').

$^{19}F$ NMR (376 MHz, $D_2O$) δ -201.15 (dd, *J* = 49.0, 13.3 Hz).

HRMS (ESI) m/z calculated for $C_{12}H_{21}FO_{10}$ (M+Na) 367.1, found 367.1.

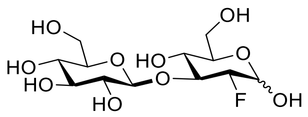

**2-deoxy-2[$^{19}$F]-fluoro-laminaribiose**

$^{1}H$ NMR (400 MHz, $D_2O$) δ 5.46 (d, *J* = 3.8 Hz, 1H), 4.94 (dd, *J* = 7.9, 2.5 Hz, 1H), 4.68 (dd, *J* = 12.1, 8.0 Hz, 2H), 4.61 (ddd, *J* = 49.5, 9.4, 3.9 Hz, 1H), 4.30 (dt, *J* = 53.2, 9.1 Hz, 1H), 4.19 (dt, *J* = 12.6, 9.3 Hz, 1H), 4.09 (dt, *J* = 14.6, 8.7 Hz, 1H), 3.98 – 3.86 (m, 4H), 3.86 – 3.70 (m, 5H), 3.62 – 3.46 (m, 7H), 3.46 – 3.39 (m, 2H), 3.39 – 3.32 (m, 2H).

$^{13}C$ NMR (100 MHz, $D_2O$) δ 102.45 (C1'α+ C1'β), 93.45 (C1β), 92.87 (C2β), 90.17 (C2α), 89.61 (C1α), 81.65 (C3β), 79.23 (C3α), 75.89 (C5'α+C5'β), 75.70 (C5β), 75.54 (C3'α+C3'β), 73.19 (C2'α+C2'β), 71.08 (C5α), 69.56 (C4'α+C4'β), 67.67 (C4α+C4β), 60.66 (C6'α+C6'β), 60.50 (C6β), 60.28 (C6α).

$^{19}F$ NMR (376 MHz, $D_2O$) δ -198.87 (dd, *J* = 51.0, 14.6 Hz), -199.20 (dd, *J* = 49.1, 12.7 Hz).

HRMS (ESI) m/z calculated for $C_{12}H_{21}FO_{10}$ (M+Na) 367.1, found 367.1.

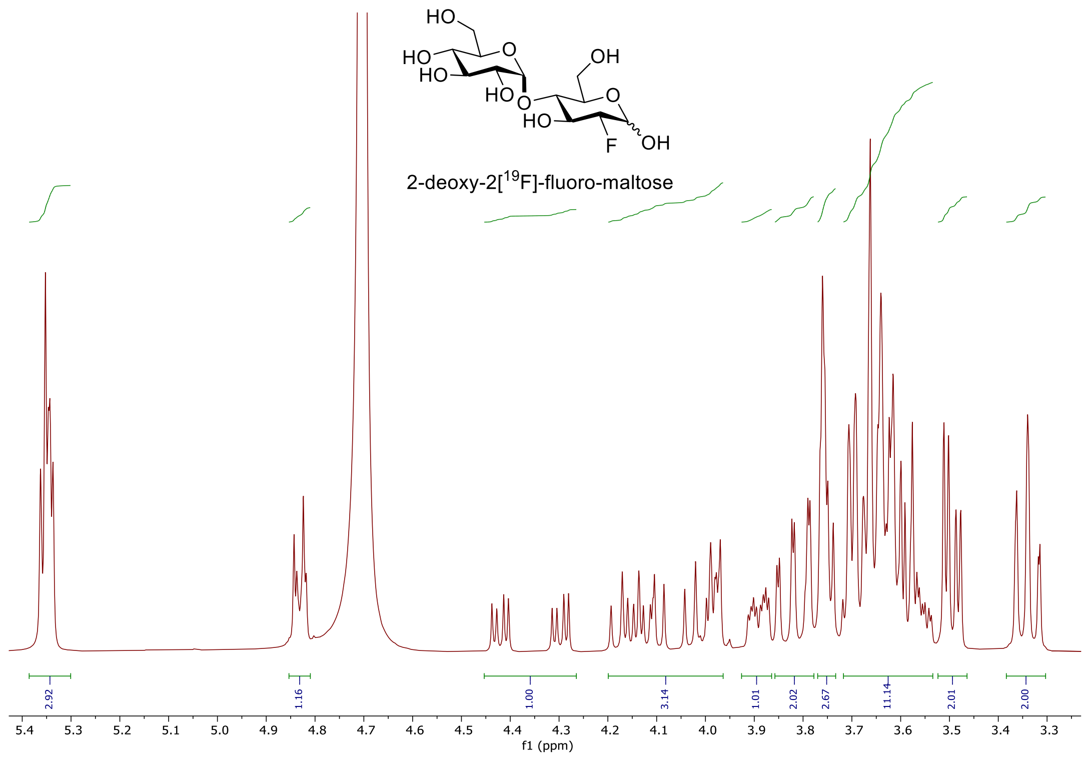

**Figure B.3.5.** $^{1}$H NMR of 2-deoxy-2[$^{19}$F]-fluoro-maltose in $D_2O$.

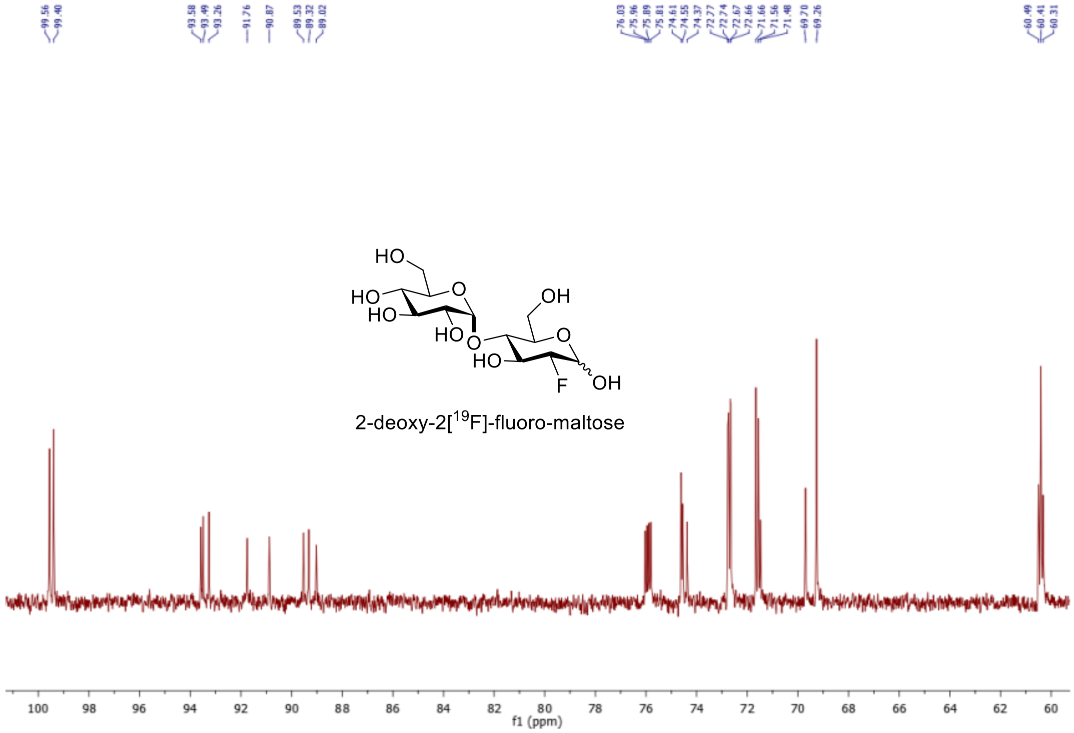

**Figure B.3.6.** $^{13}$C NMR of 2-deoxy-2[$^{19}$F]-fluoro-maltose in $D_2O$.

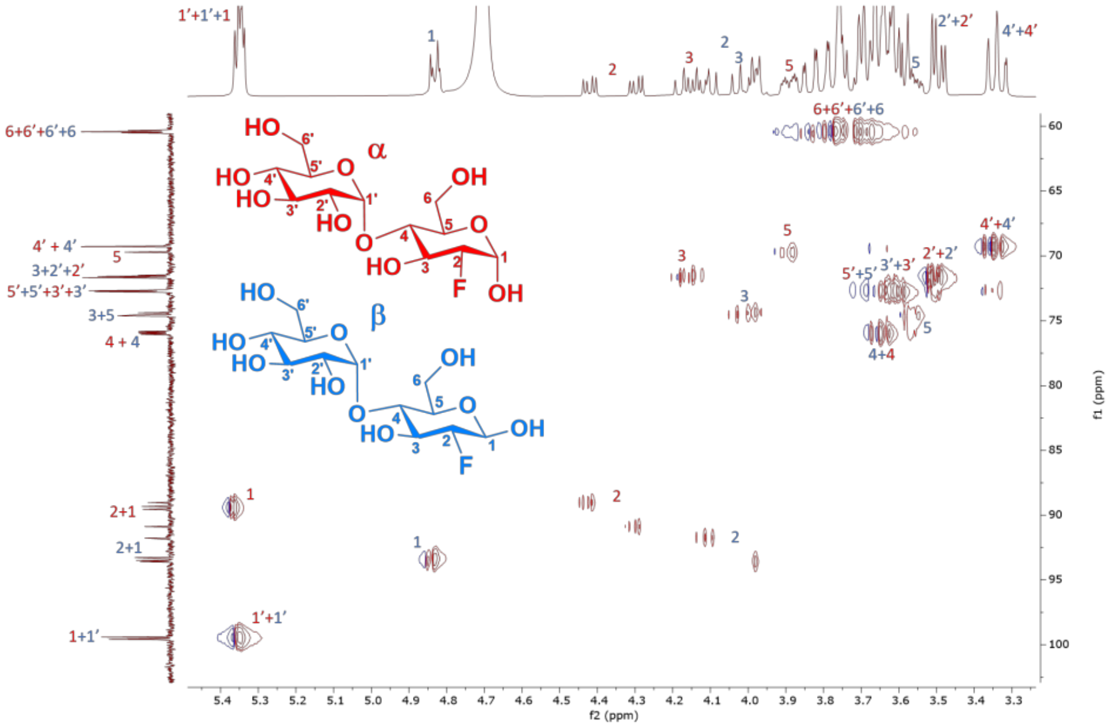

**B.3.7.** HSQC NMR of 2-deoxy-2[$^{19}$F]-fluoro-maltose in $D_2O$.

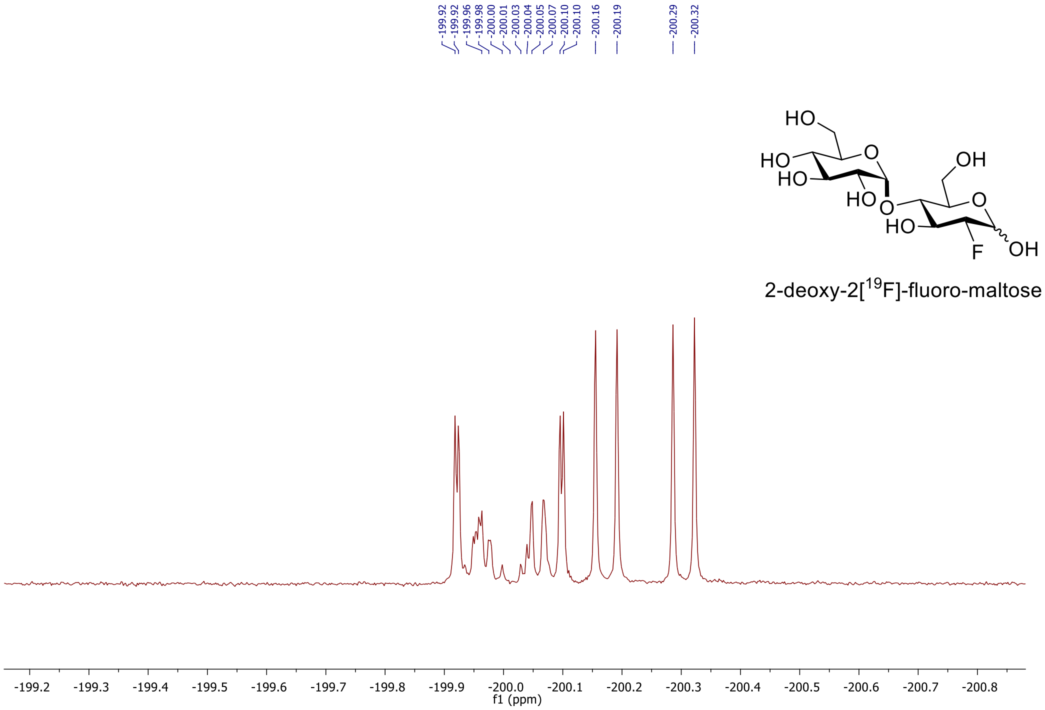

**Figure B.3.8.** $^{19}$F NMR of 2-deoxy-2[$^{19}$F]-fluoro-maltose in $D_2O$.

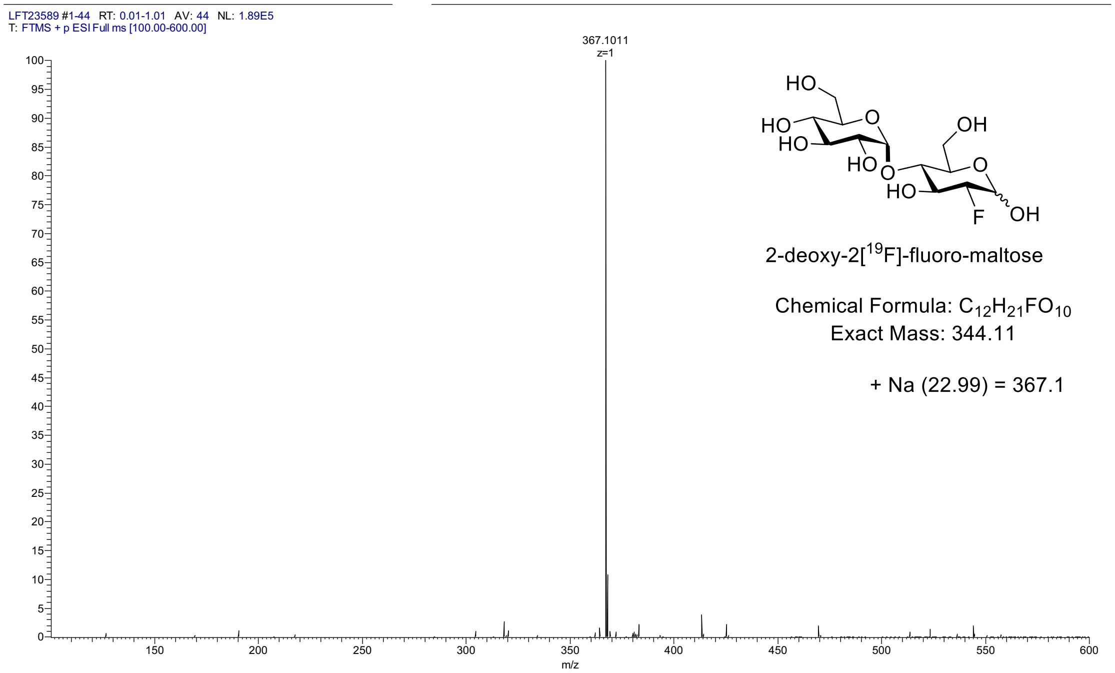

**Figure B.3.9.** HRMS of 2-deoxy-2[$^{19}$F]-fluoro-maltose.

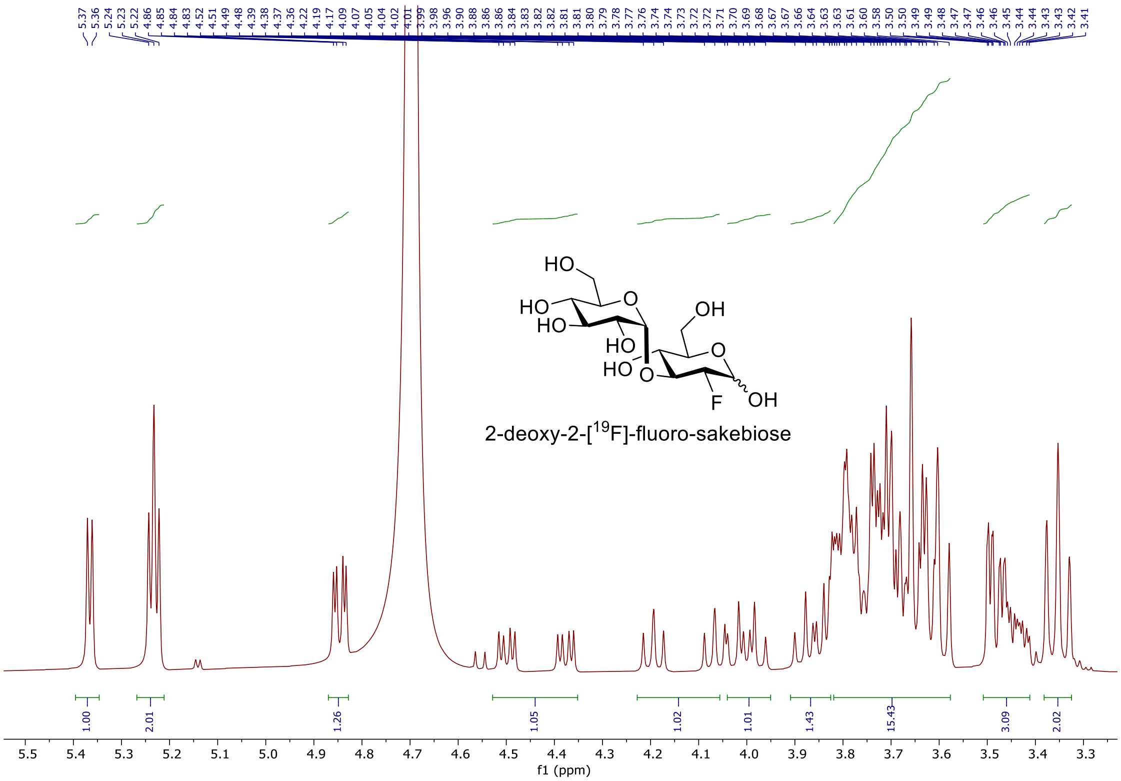

**Figure B.3.10.** $^{1}$H NMR of 2-deoxy-2[$^{19}$F]-fluoro-sakebiose in $D_2O$.

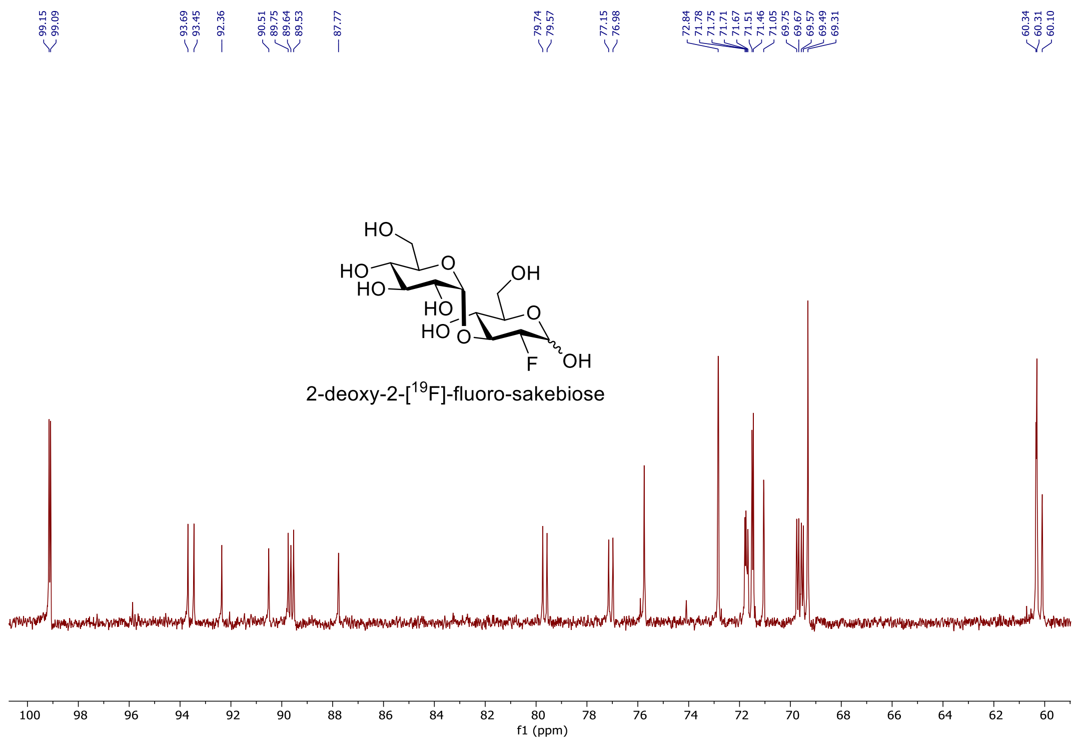

**Figure B.3.11.** $^{13}C$ NMR of 2-deoxy-2[$^{19}F$]-fluoro-sakebiose in $D_2O$.

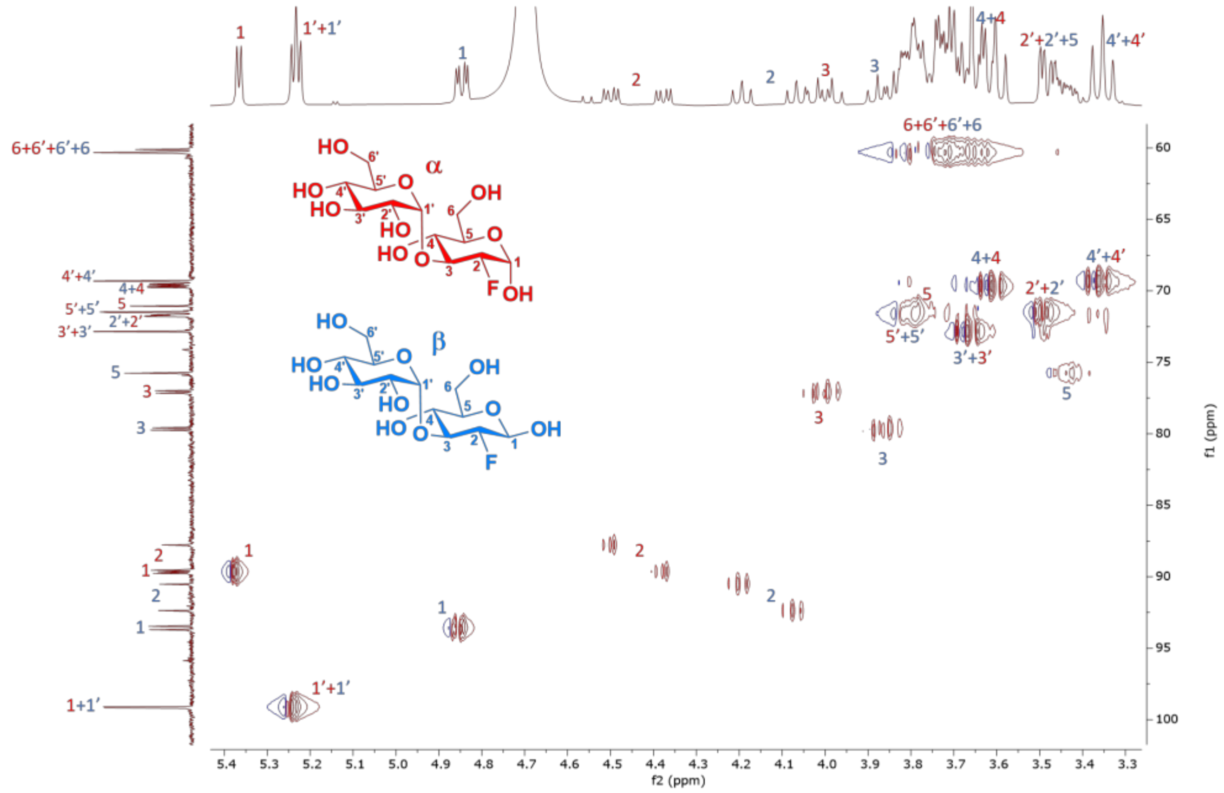

**Figure B.3.12.** HSQC NMR of 2-deoxy-2[$^{19}F$]-fluoro-sakebiose in $D_2O$.

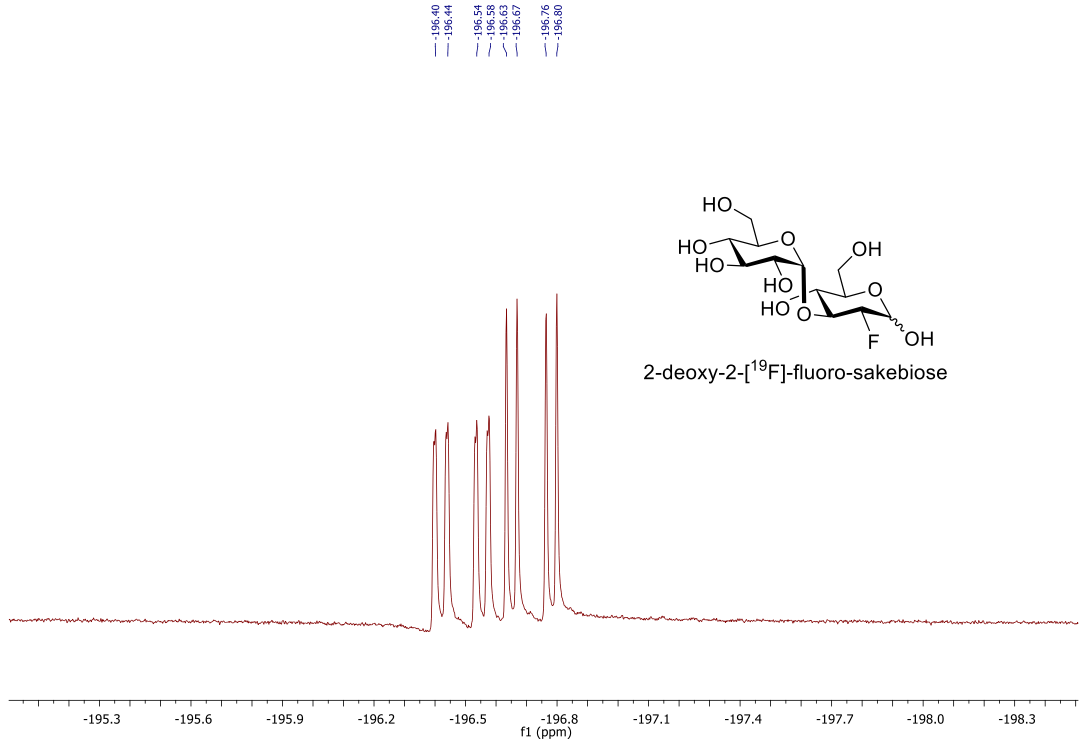

**Figure B.3.13.** $^{19}$F NMR of 2-deoxy-2[$^{19}$F]-fluoro-sakebiose in $D_2O$.

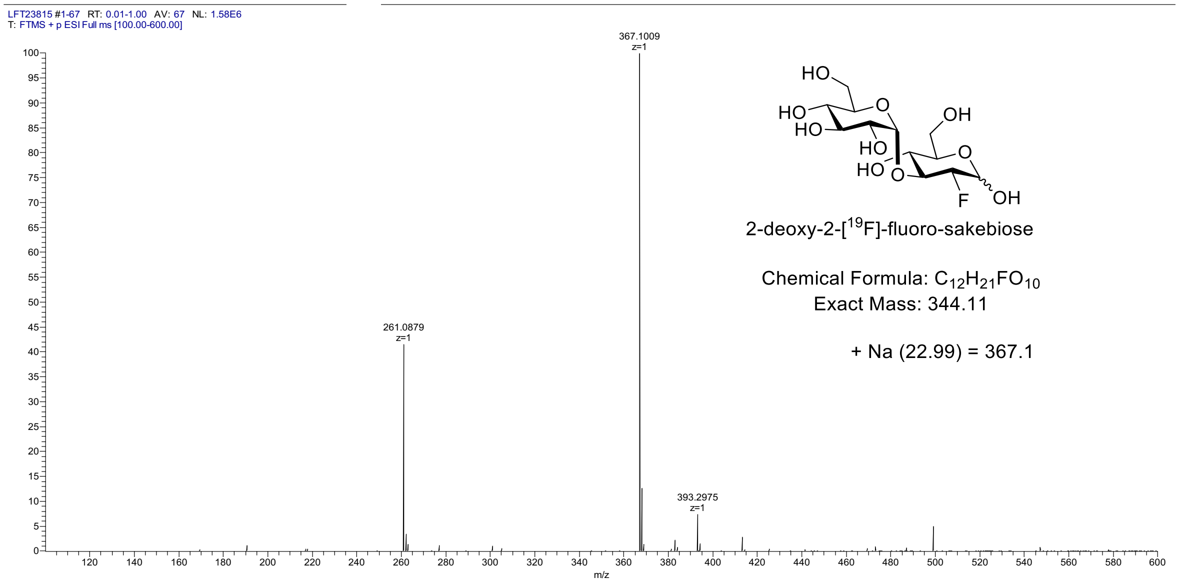

**Figure B.3.14.** HRMS of 2-deoxy-2[$^{19}$F]-fluoro-sakebiose.

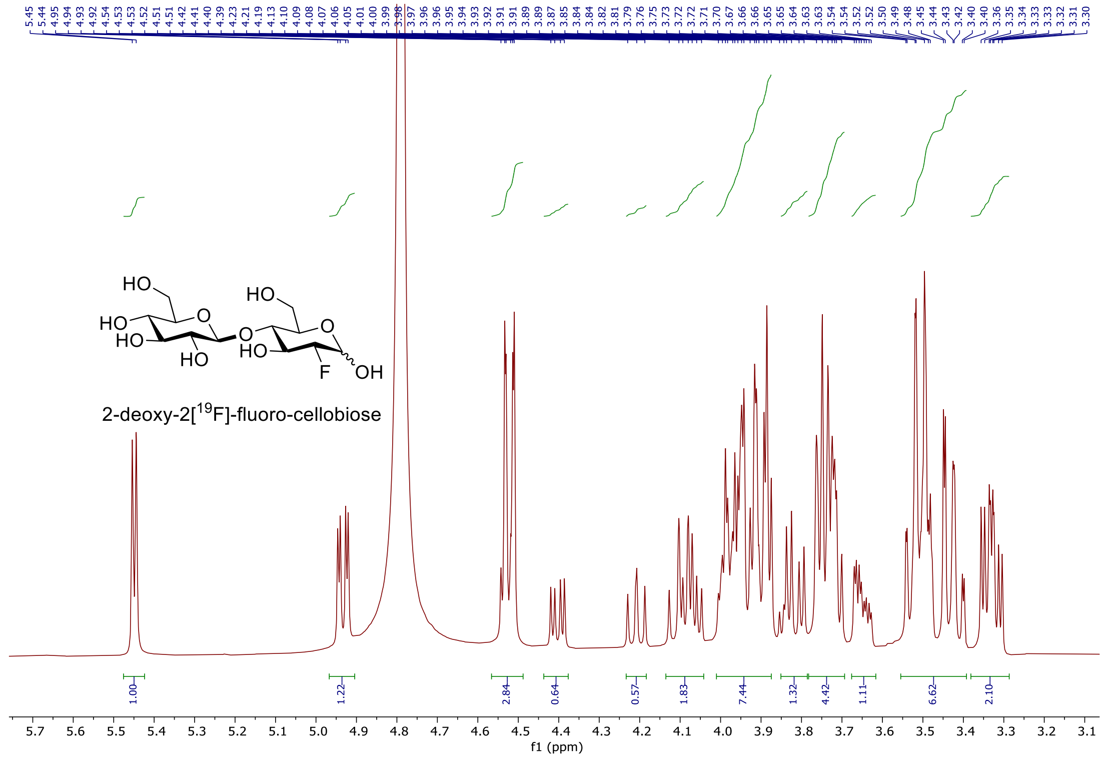

**Figure B.3.15.** $^{1}H$ NMR of 2-deoxy-2[$^{19}$F]-fluoro-cellobiose in $D_2O$.

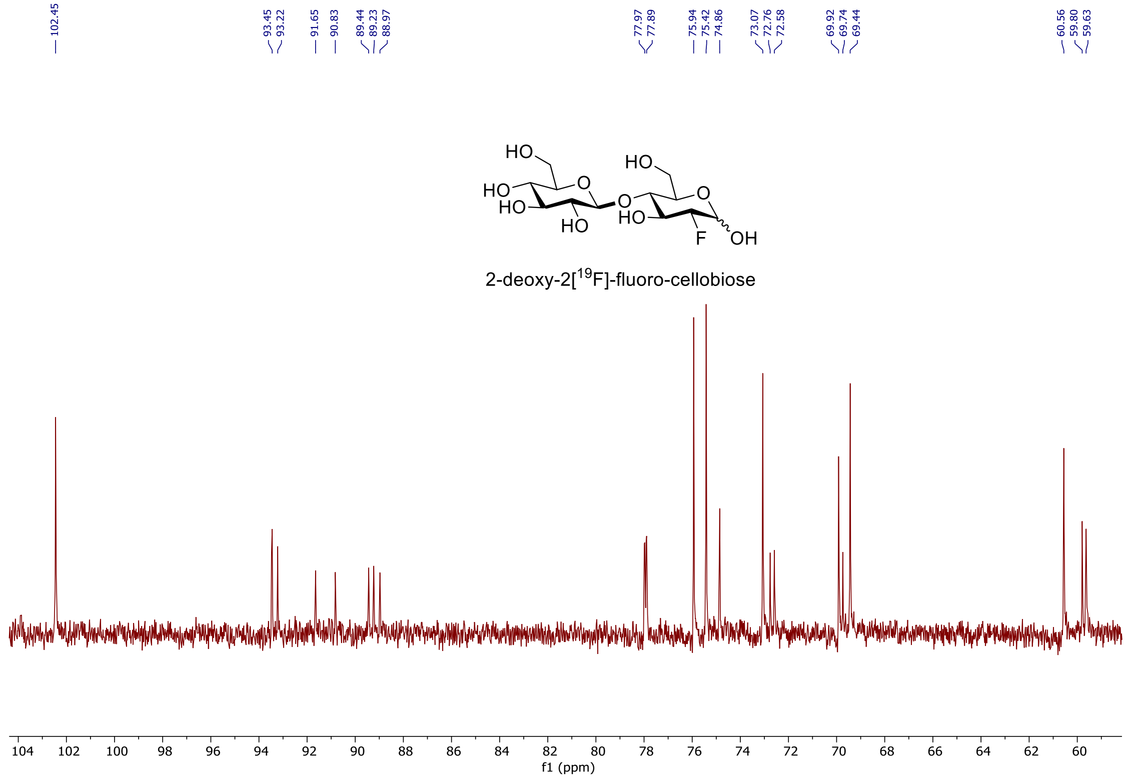

**Figure B.3.16.** $^{13}C$ NMR of 2-deoxy-2[$^{19}$F]-fluoro-cellobiose in $D_2O$.

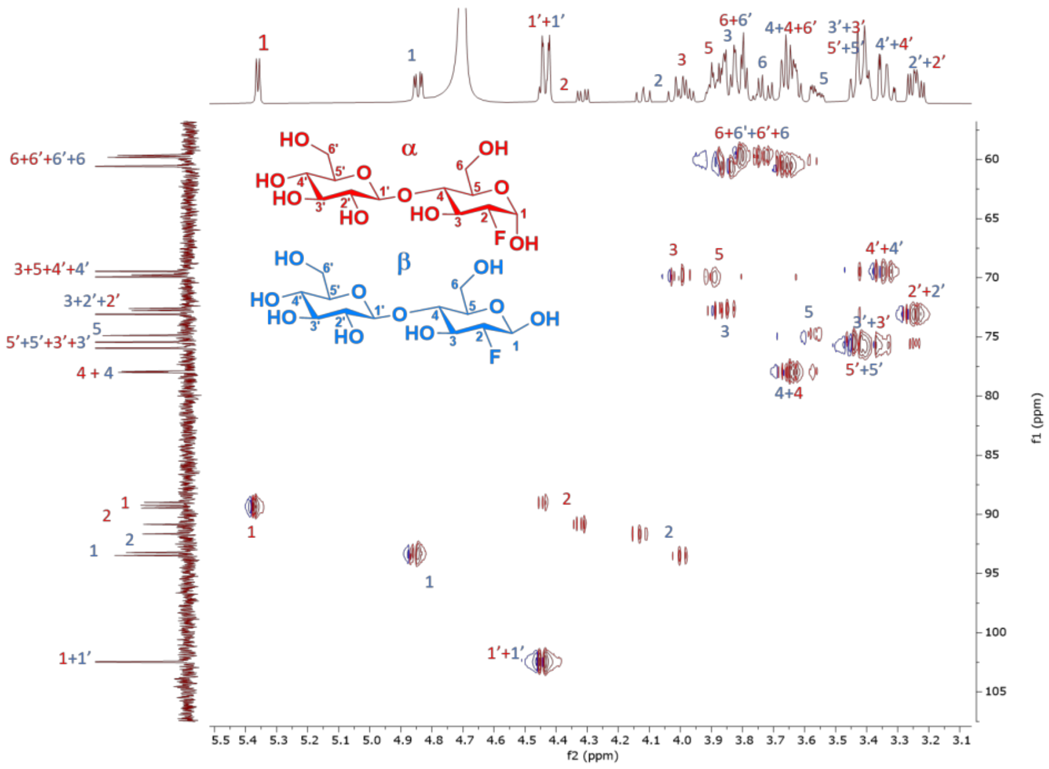

**Figure B.3.17.** HSQC NMR of 2-deoxy-2[19F]-fluoro-cellobiose in $D_2O$

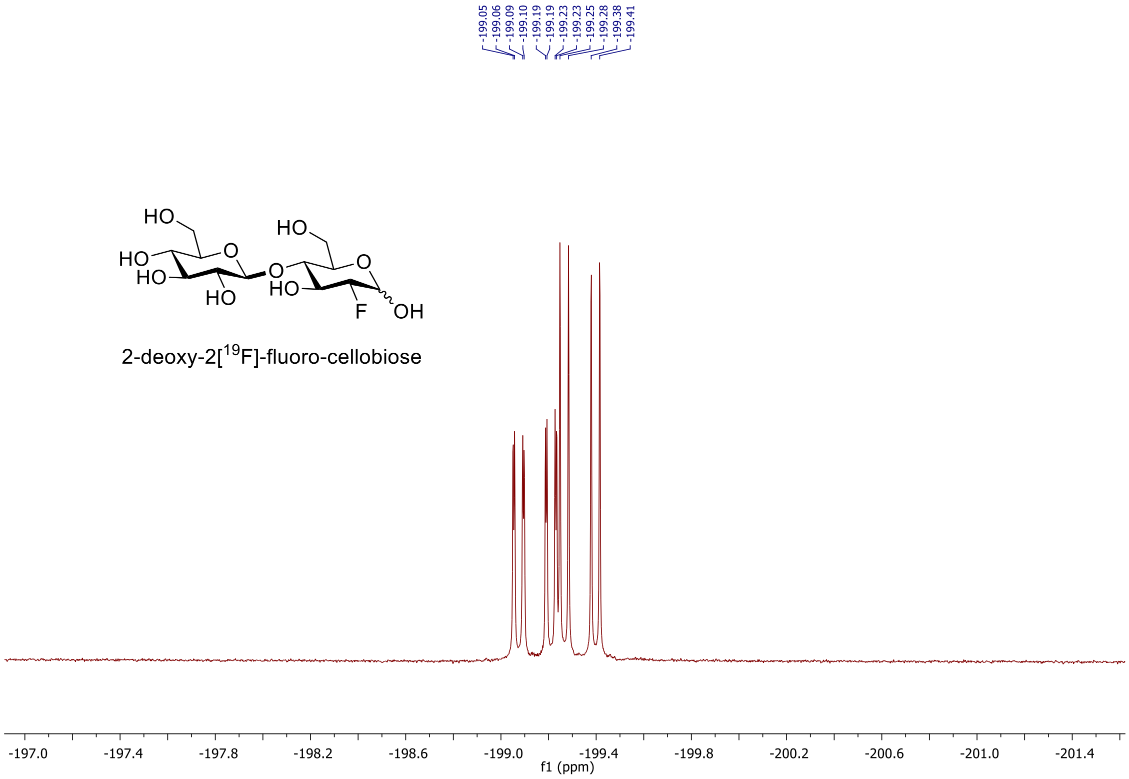

**Figure B.3.18.** $^{19}F$ NMR of 2-deoxy-2[19F]-fluoro-cellobiose in $D_2O$.

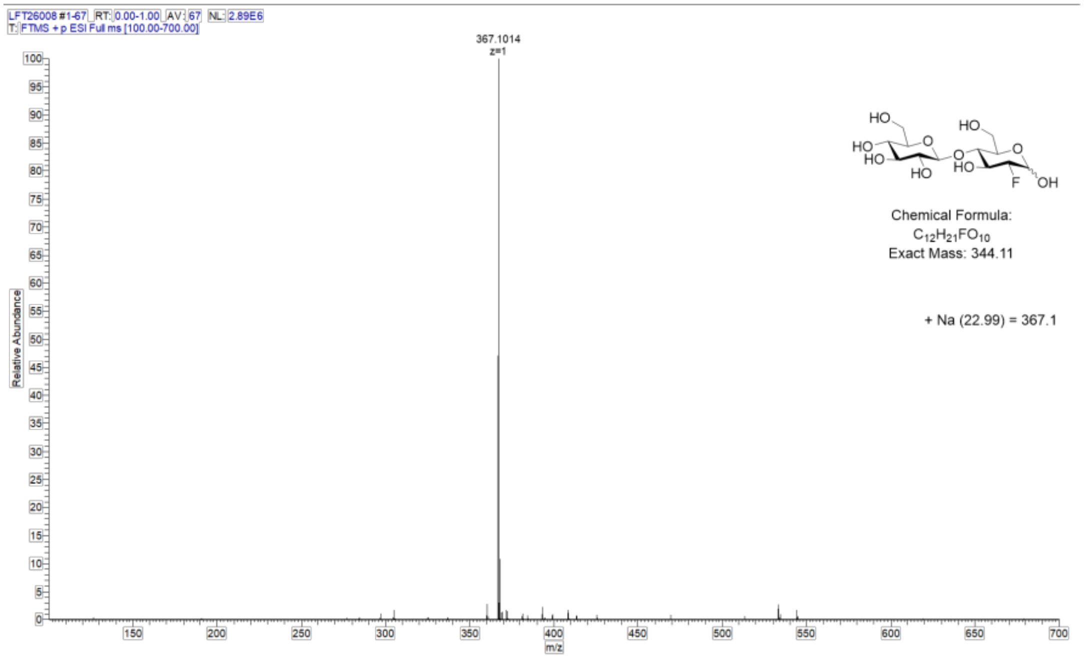

**Figure B.3.19.** HRMS of 2-deoxy-2[19F]-fluoro-cellobiose.

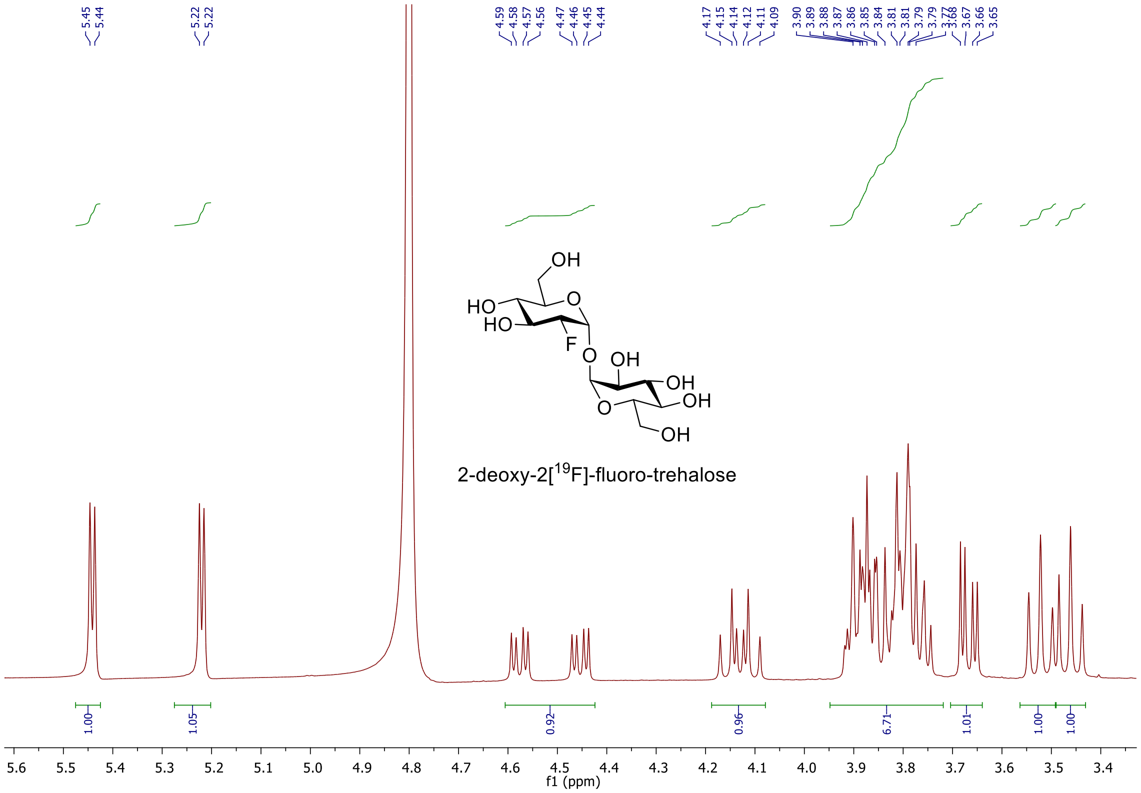

**Figure B.3.20.** $^{1}H$ NMR of 2-deoxy-2[$^{19}F$]-fluoro-trehalose in $D_2O$.

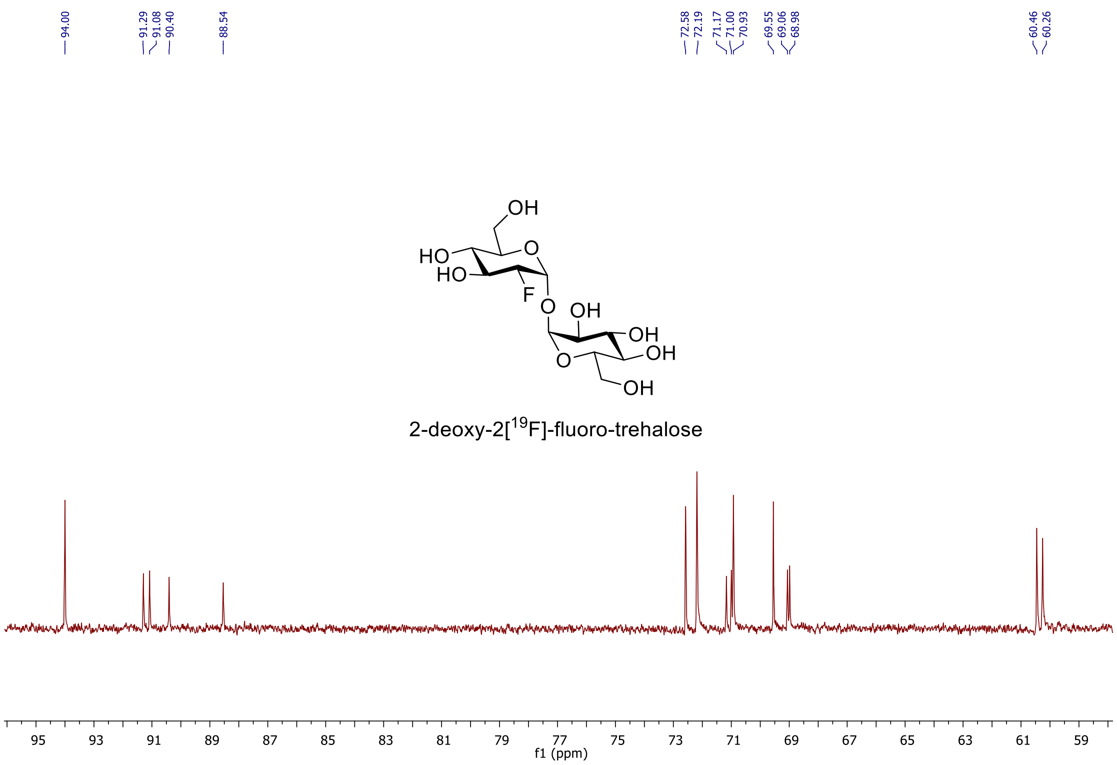

**Figure B.3.21.** $^{13}C$ NMR of 2-deoxy-2[$^{19}$F]-fluoro-trehalose in $D_2O$.

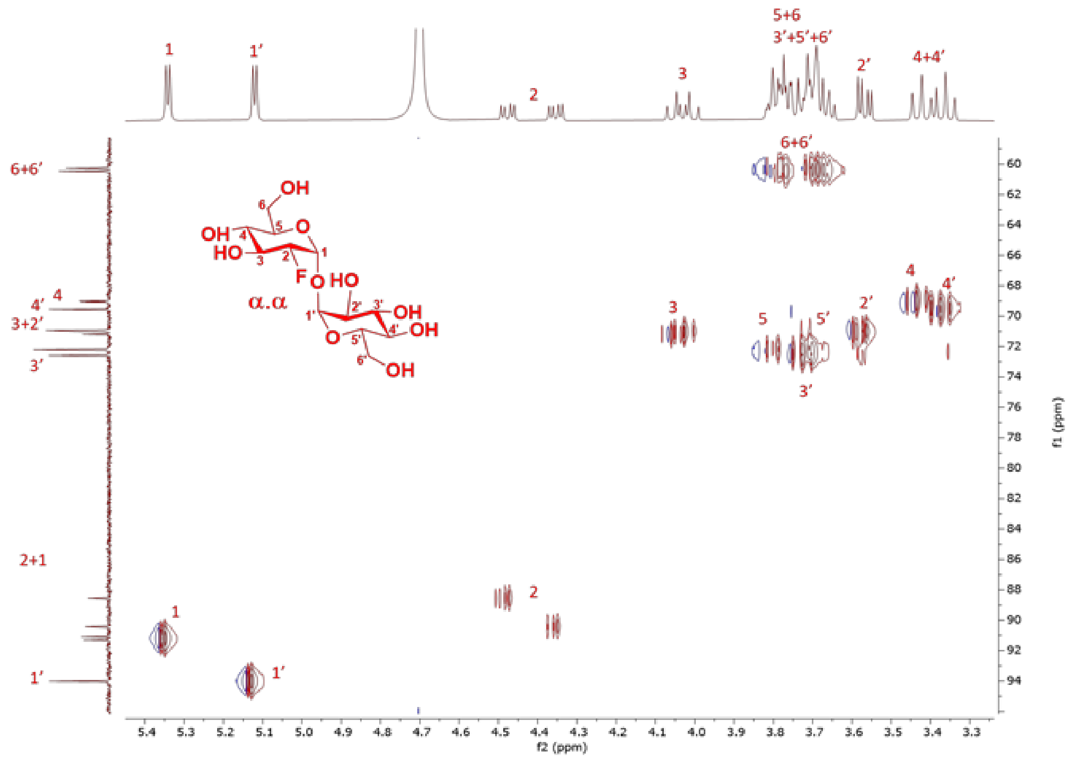

**Figure B.3.22.** HSQC NMR of 2-deoxy-2[$^{19}$F]-fluoro-trehalose in $D_2O$

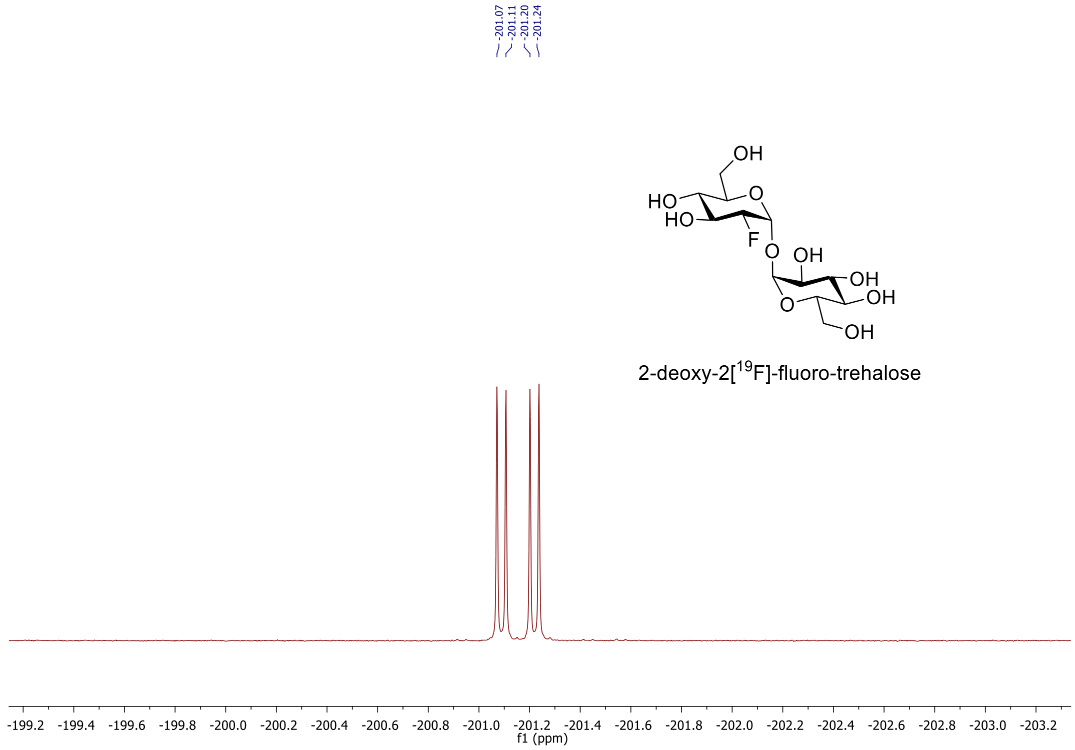

**Figure B.3.23.** $^{19}F$ NMR of 2-deoxy-2[$^{19}F$]-fluoro-trehalose in $D_2O$.

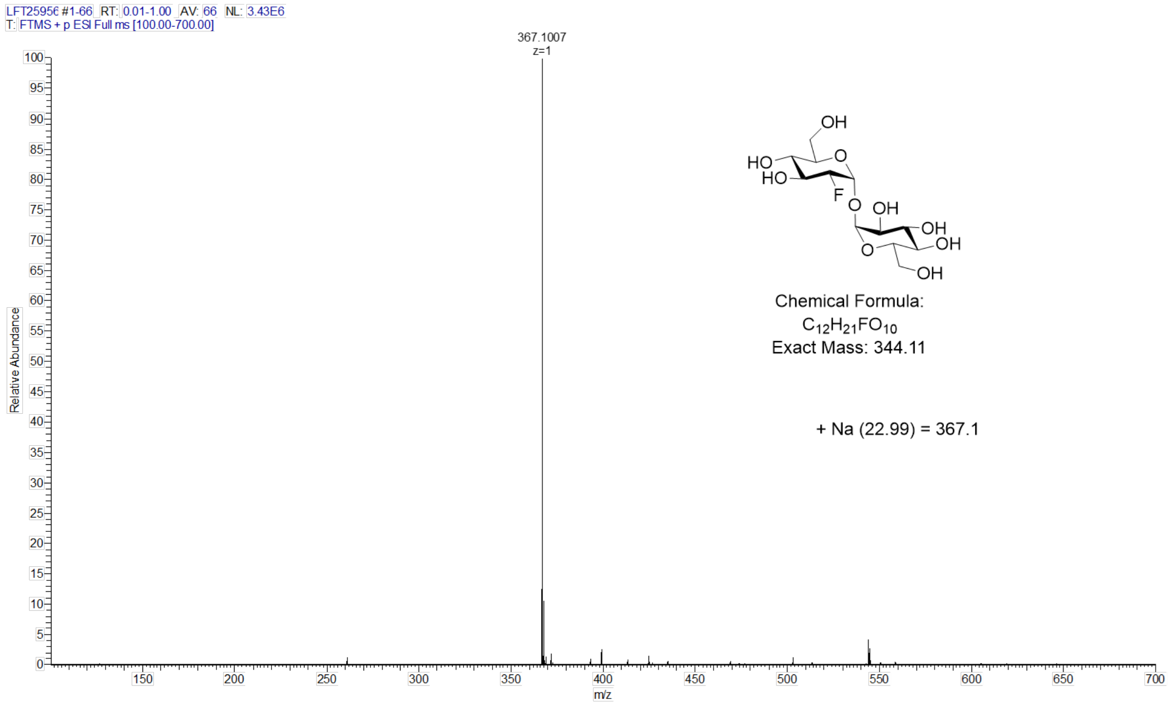

**Figure B.3.24.** HRMS of 2-deoxy-2[$^{19}F$]-fluoro-trehalose.

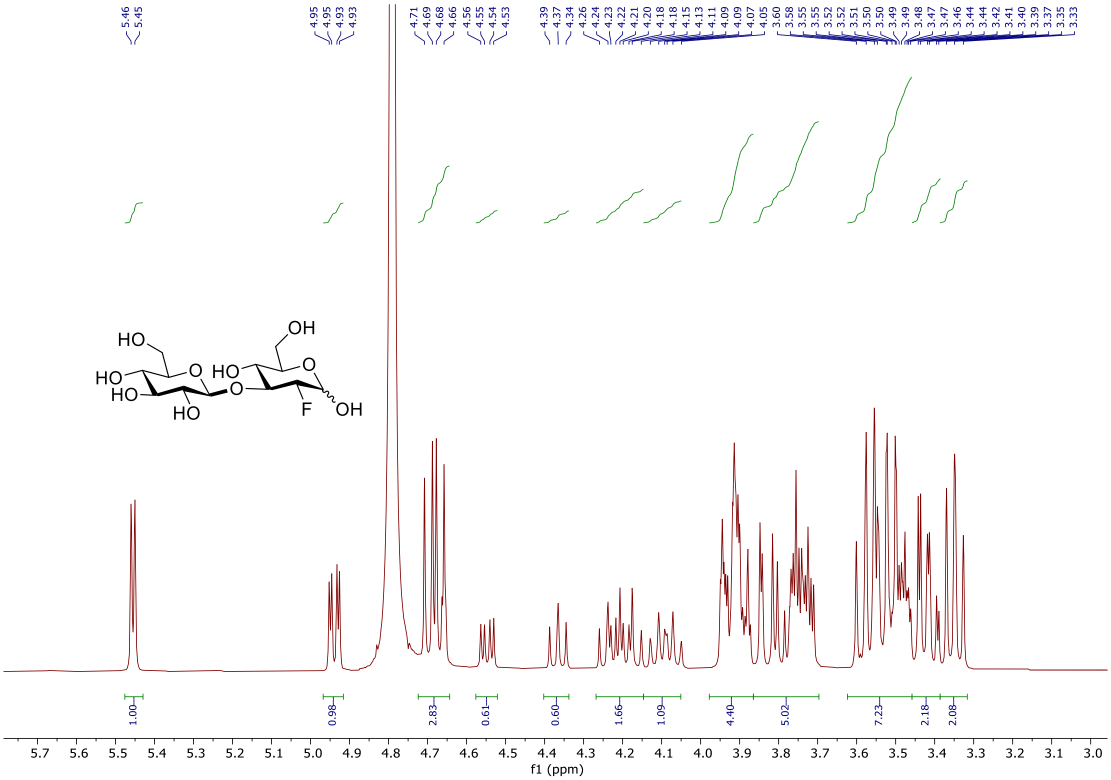

**Figure B.3.25.** $^{1}$H NMR of 2-deoxy-2[$^{19}$F]-fluoro-laminaribiose in $D_2O$.

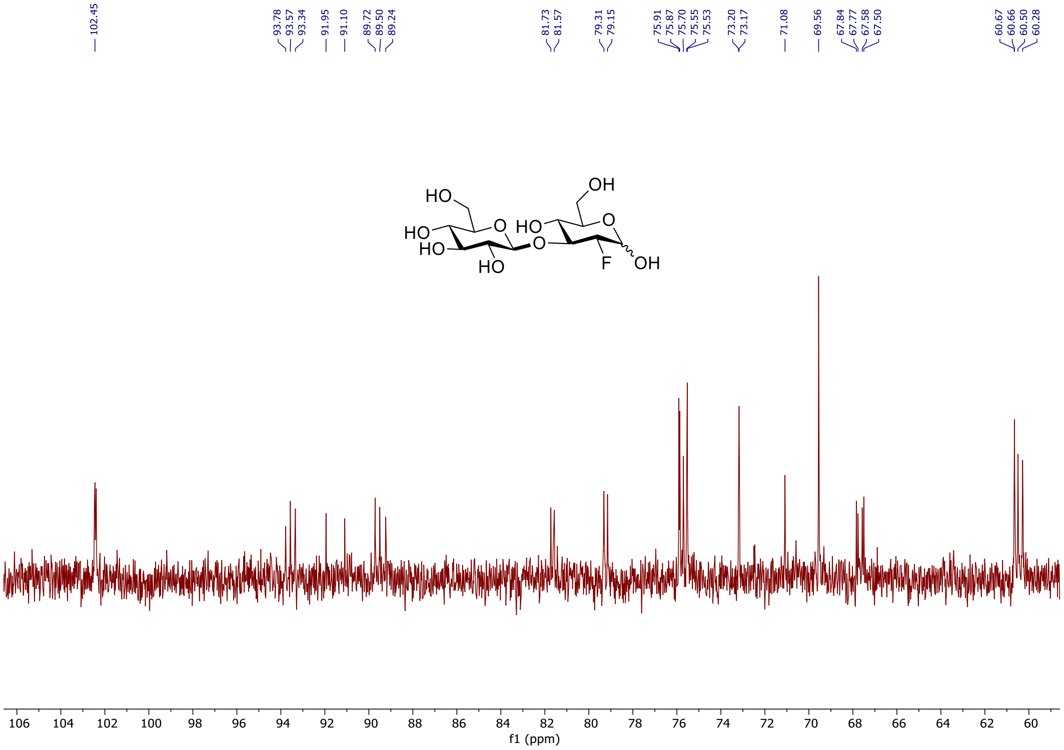

**Figure B.3.26.** $^{13}$C NMR of 2-deoxy-2[$^{19}$F]-fluoro-laminaribiose in $D_2O$.

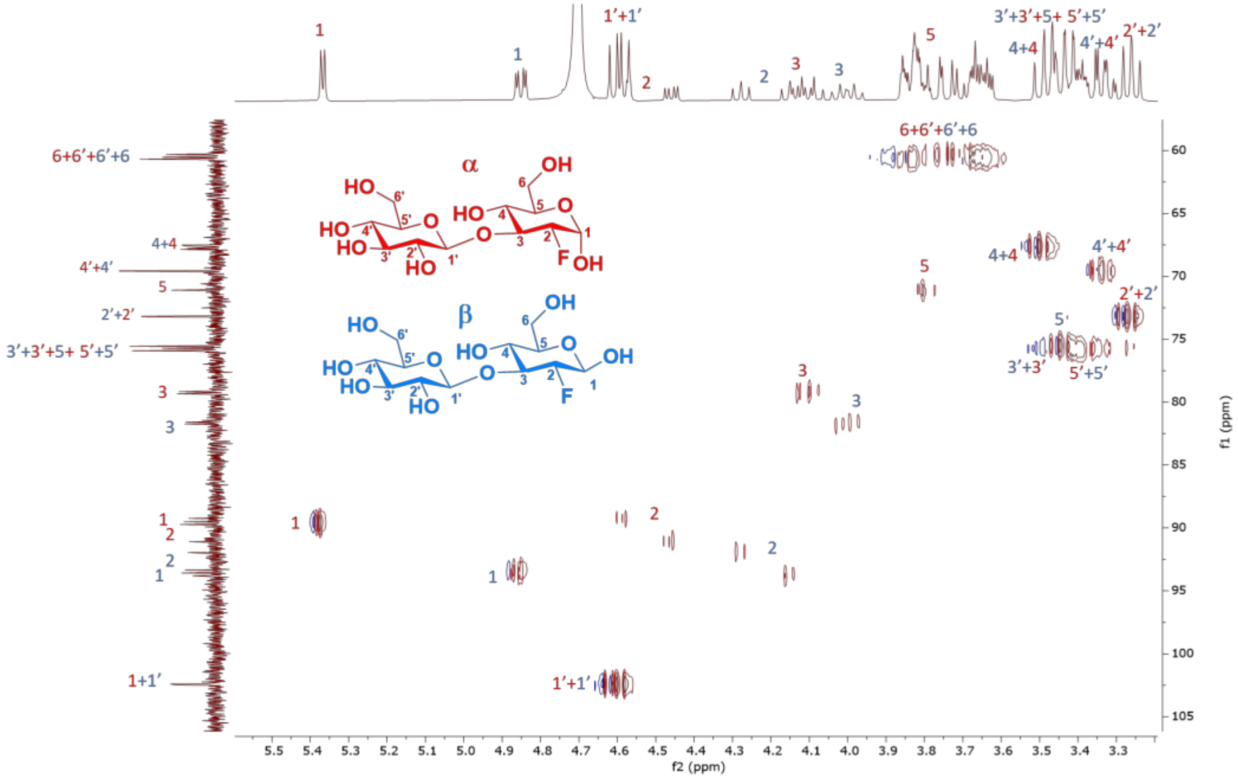

**Figure B.3.27.** HSQC NMR of 2-deoxy-2[[19]F]-fluoro-laminaribiose in $D_2O$.

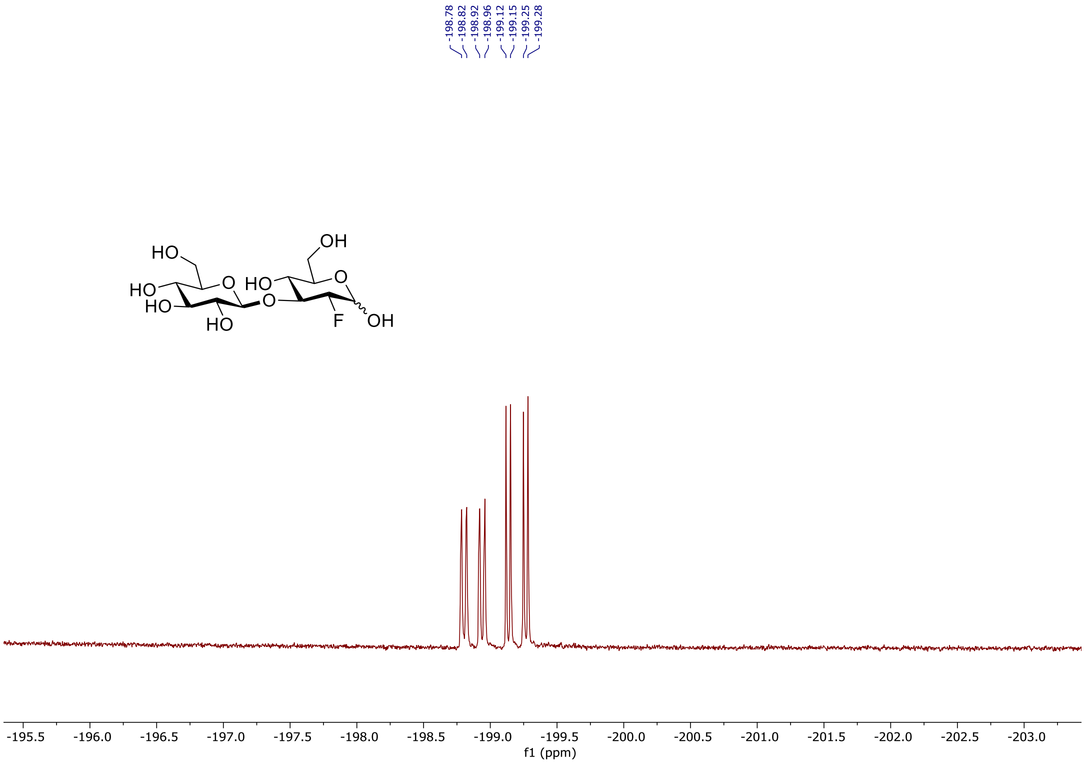

**Figure B.3.28.** [19]F NMR of 2-deoxy-2[[19]F]-fluoro-laminaribiose in $D_2O$.

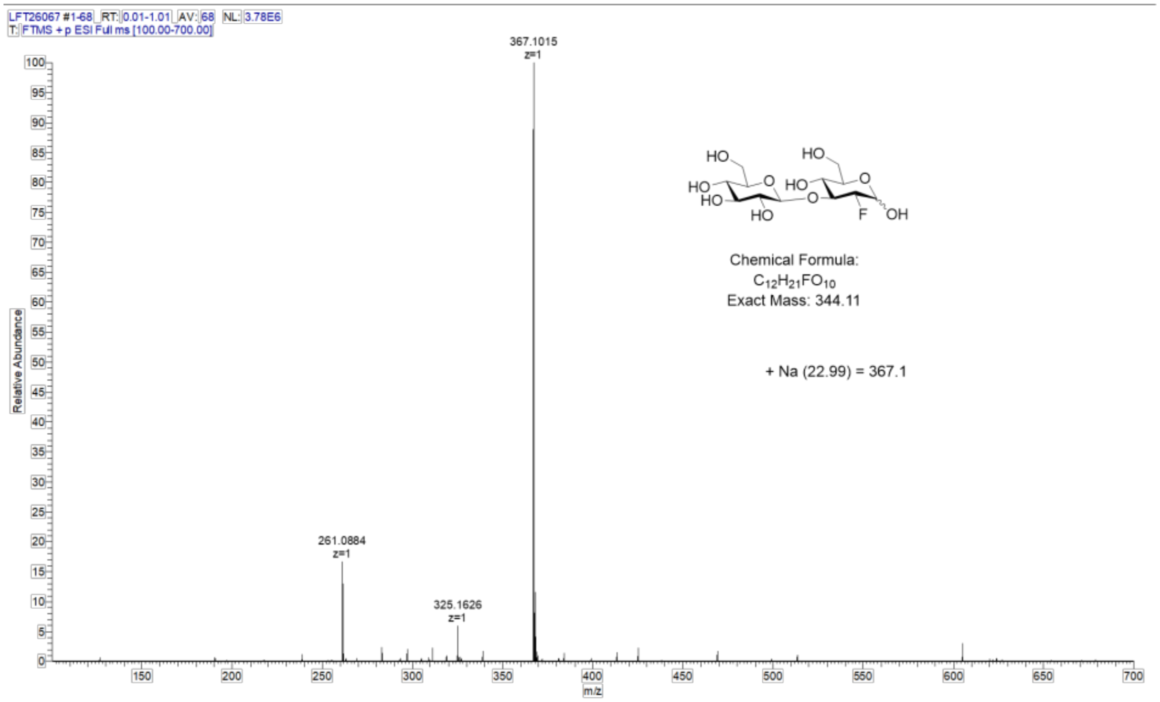

**Figure B.3.29.** HRMS of 2-deoxy-2[$^{19}$F]-fluoro-laminaribiose.

### C. Radiochemistry

#### C.1. Radiochemical procedure for 2-deoxy-[$^{18}$F]-fluoro-D-glucose ([$^{18}$F]FDG) production:

2-deoxy-[$^{18}$F]-fluoro-D-glucose ([$^{18}$F]FDG) was produced in aqueous solution using standard methods.[5]

#### C.2. Radiochemical procedure for 2-deoxy-[$^{18}$F]-fluoro-D-sorbitol ([$^{18}$F]FDS) production:

2-deoxy-[$^{18}$F]-fluoro-D-sorbitol ([$^{18}$F]FDS) was synthesized using procedure reported by Weinstein et al.[6]

**C.3. Radiosynthesis of 2-deoxy-[$^{18}$F]-fluoro-maltose ([$^{18}$F]FDM) and 2-deoxy-2-[$^{18}$F]-fluoro-sakebiose ([$^{18}$F]FSK):**

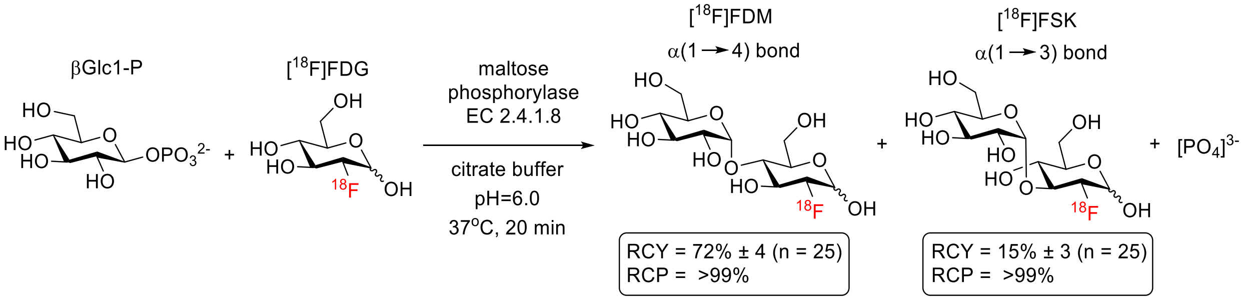

In a 4 mL borosilicate vial containing PTFE stir bar, maltose phosphorylase (EC 2.4.1.8, Sigma Aldrich), (0.3 mg, 3 units) and βGlc1-P (6 mg, 0.020 mmol) were added. A dose of [$^{18}$F]FDG (10-15 mCi) in citrate buffer (0.1M, pH=6.0, 0.4-0.5 mL) was directly transferred to the vial and the mixture was stirred at 37 °C for 20 min. The mixture was diluted with MeCN then filtered through C18 light cartridge, before being purified via semi prep HPLC (YMC-Pack Polyamine II, 250 X 10 mm) using mobile phase 73% MeCN/27 % $H_2O$. Both [$^{18}$F]FDM and [$^{18}$F]FSK were isolated in 5-7mL fractions. The fractions were then diluted with MeCN (40 mL) before being passed through Sep-pak Plus $NH_2$ Cartridge at 5 mL/min to trap each dimer product. After flushing the cartridge with air and $N_2$ gas, they were eluted using saline solution for direct formulation before use in vitro or in vivo. [$^{18}$F]FDM (RCY= 72% ± 4%, RCP=99%), [$^{18}$F]FSK (15%± 3%, RCP=99%) (N=25). Chemical purity of both [$^{18}$F]FDM and [$^{18}$F]FSK were confirmed by analytical HPLC.

#### HPLC analysis of [$^{18}$F]FDM and [$^{18}$F]FSK

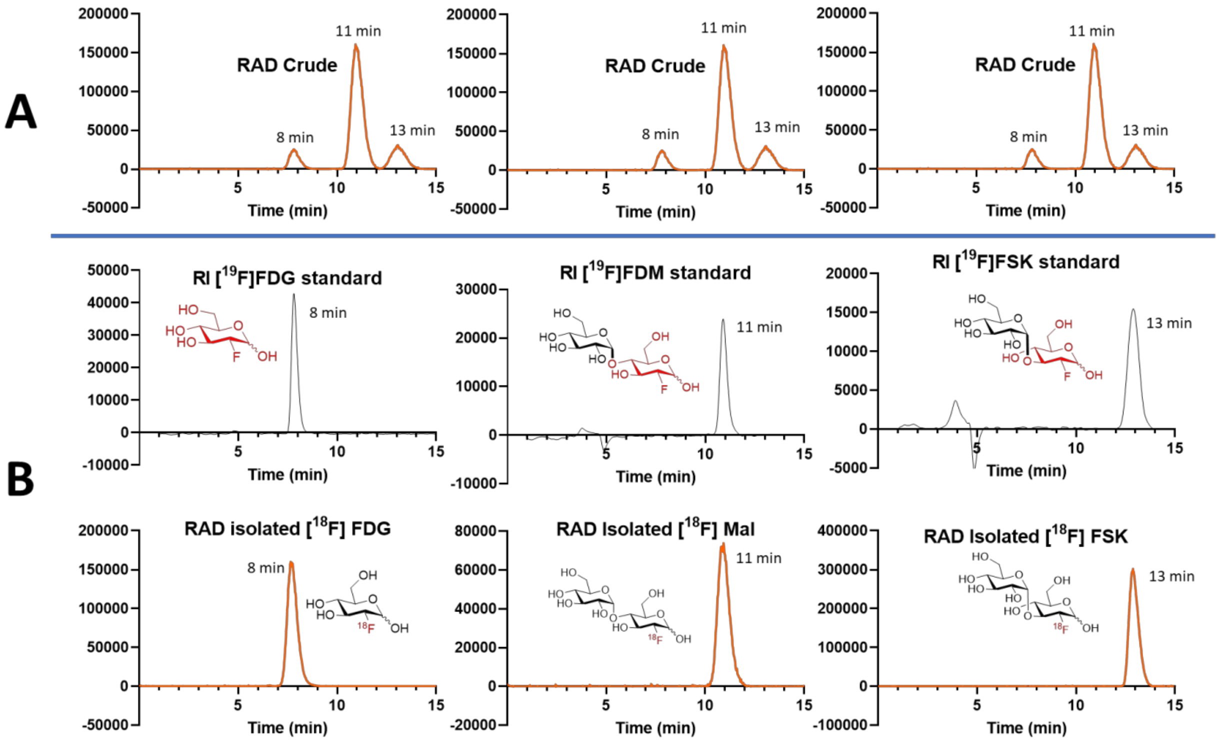

**Figure C.3.1.** HPLC analysis of 2-deoxy-[$^{18}$F]-fluoro-maltose ([$^{18}$F]FDM) and 2-deoxy-2-[$^{18}$F]-fluoro-sakebiose ([$^{18}$F]FSK) with YMC-Pack Polyamine II, 250 X 4.6 mm using mobile phase 73% MeCN/27 % $H_2O$. A) Injection of crude, radioactivity (RAD) detection B) Co-injection of "cold" $^{19}$F standard and "hot" $^{18}$F isolated tracer, with both refractive index (RI) detection and radioactivity (RAD) detection.

**C.4. Radiosynthesis of 2-deoxy-2[$^{18}$F]-fluoro-cellobiose ([$^{18}$F]FDC):**

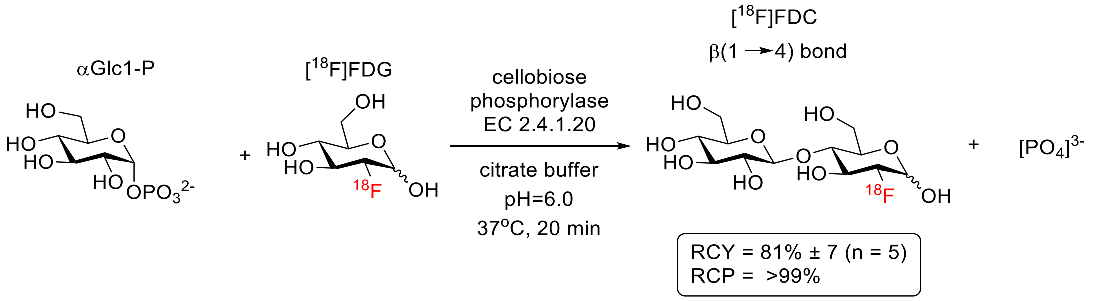

In a 4 mL borosilicate vial containing PTFE stir bar, cellobiose phosphorylase (EC 2.4.1.20, Prof. Bernd Nidetzky lab, Graz University of Technology), (0.3 mg, 3 units) and αGlc1-P (6 mg, 0.020 mmol) were added. A dose of [$^{18}$F]FDG (10-15 mCi) in citrate buffer (0.1M, pH=6.0, 0.4-0.5 mL) was directly transferred to the vial and the mixture was stirred at 37 °C for 20 min. The mixture was diluted with MeCN then filtered through C18 light cartridge, before being purified via semi prep HPLC (YMC-Pack Polyamine II, 250 X 10 mm) using mobile phase 73% MeCN/27 % $H_2O$. [$^{18}$F]FDC was isolated in 5-7mL fractions. The fractions were then diluted with MeCN (40 mL) before being passed through Sep-pak Plus $NH_2$ Cartridge at 5 mL/min to trap each dimer product. After flushing the cartridge with air and $N_2$ gas, the tracer was eluted using saline solution for further analysis. [$^{18}$F]FDC (RCY= 81% ± 7%, RCP=99%), (N=5). Chemical purity of [$^{18}$F]FDC was confirmed by analytical HPLC.

### HPLC analysis of [$^{18}$F]FDC

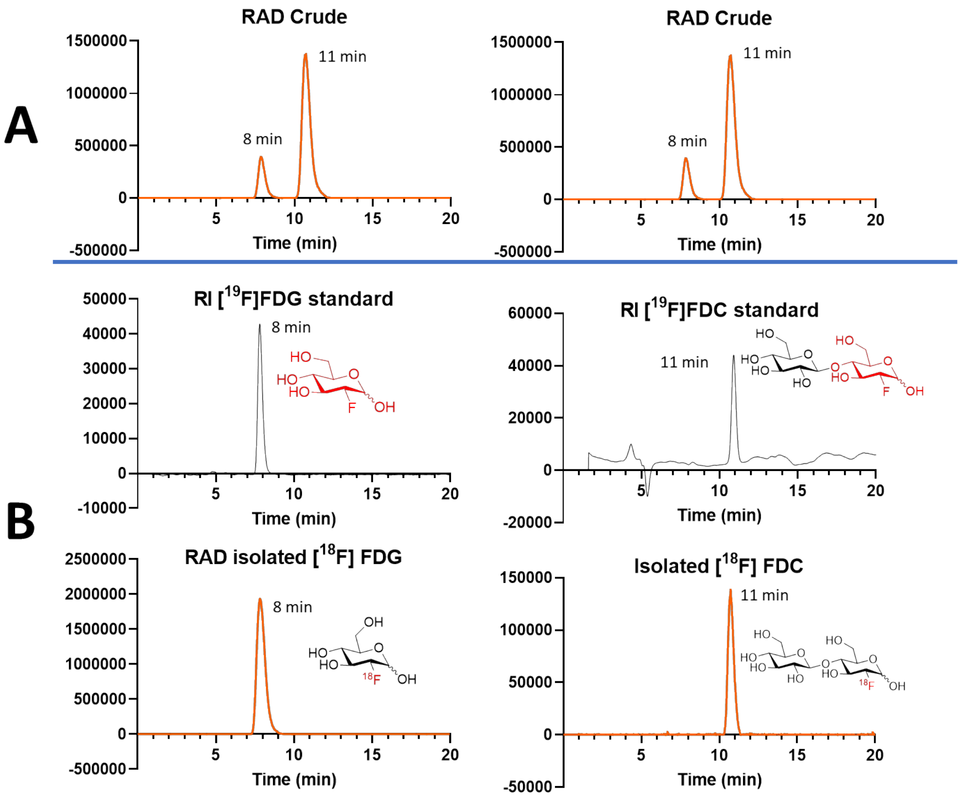

**Figure C.4.1.** HPLC analysis of 2-deoxy-2[$^{18}$F]-fluoro-cellobiose ([$^{18}$F]FDC) with YMC-Pack Polyamine II, 250 X 4.6 mm using mobile phase 73% MeCN/27 % $H_2O$. A) Injection of crude, radioactivity (RAD) detection B) Co-injection of “cold” $^{19}$F standard and “hot” $^{18}$F isolated tracer, with both refractive index (RI) detection and radioactivity (RAD) detection.

### C.5. Radiosynthesis of 2-deoxy-2[$^{18}$F]-fluoro-trehalose ([$^{18}$F]FDT):

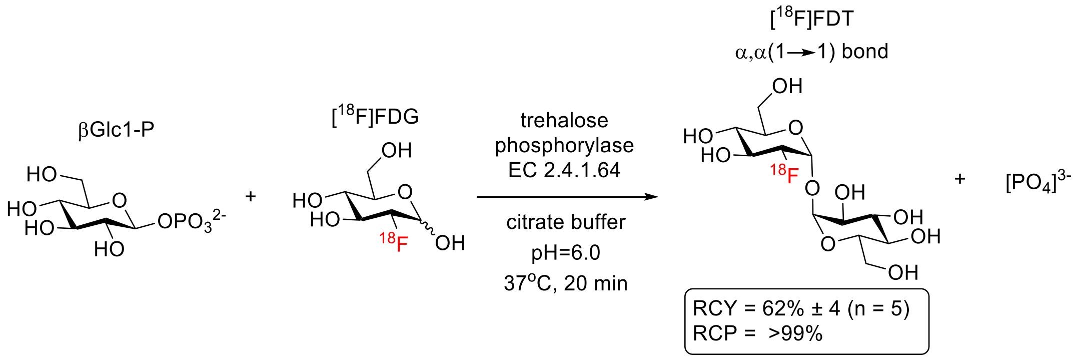

In a 4 mL borosilicate vial containing PTFE stir bar, trehalose phosphorylase (EC 2.4.1.64, Prof. Tom Desmet lab, Ghent University), (0.3 mg, 3 units) and βGlc1-P (6 mg, 0.020 mmol) were added. A dose of [$^{18}$F]FDG (10-15 mCi) in citrate buffer (0.1M, pH=6.0, 0.4-0.5 mL) was directly transferred to the vial and the mixture was stirred at 37 °C for 20 min. The mixture was diluted with MeCN then filtered through C18 light cartridge, before being purified via semi prep HPLC (YMC-Pack Polyamine II, 250 X 10 mm) using mobile phase 73% MeCN/27 % $H_2O$. [$^{18}$F]FDT was isolated in 5-7mL fractions. The fractions were then diluted with MeCN (40 mL) before being passed through Sep-pak Plus $NH_2$ Cartridge at 5 mL/min to trap each dimer product. After flushing the cartridge with air and $N_2$ gas, the tracer was eluted using saline solution for further analysis. [$^{18}$F]FDT (RCY= 62% ± 4%, RCP=99%), (N=5). Chemical purity of [$^{18}$F]FDT was confirmed by analytical HPLC.

**HPLC analysis of [$^{18}$F]FDT**

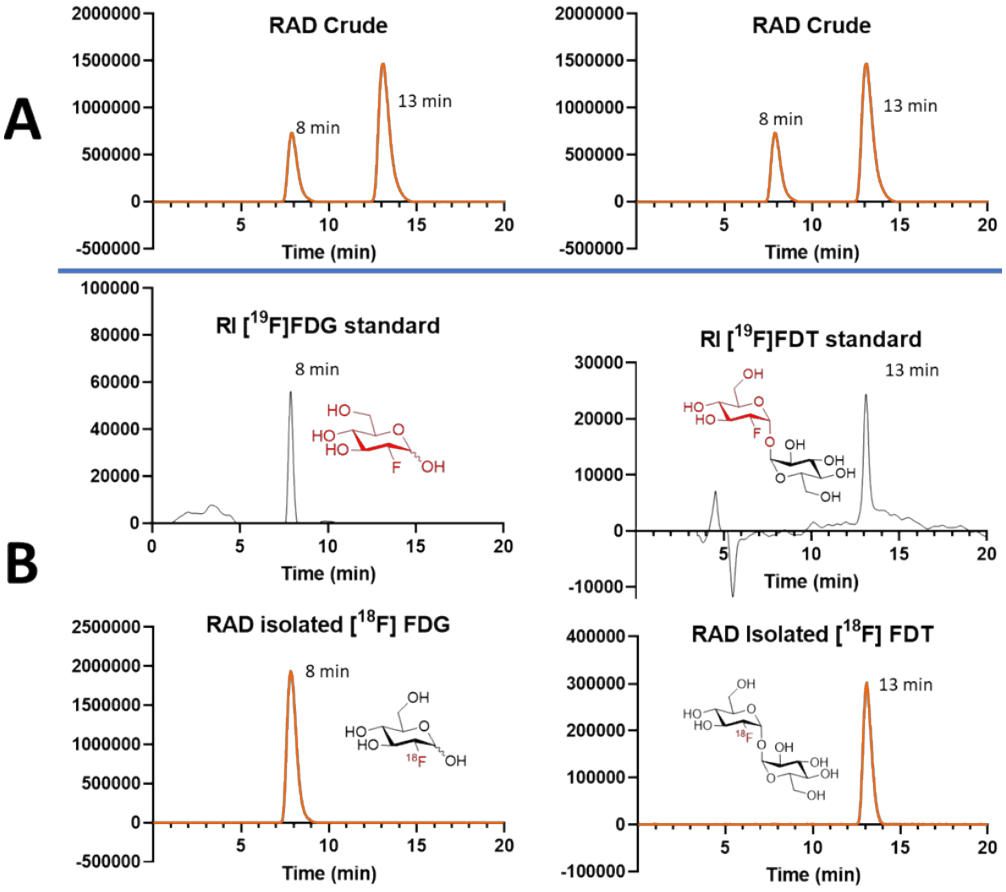

**Figure C.5.1.** HPLC analysis of 2-deoxy-2[$^{18}$F]-fluoro-trehalose ([$^{18}$F]FDT) with YMC-Pack Polyamine II, 250 X 4.6 mm using mobile phase 73% MeCN/27% $H_2O$. A) Injection of crude, radioactivity (RAD) detection B) Co-injection of “cold” $^{19}$F standard and “hot” $^{18}$F isolated tracer, with both refractive index (RI) detection and radioactivity (RAD) detection.

**C.6. Radiosynthesis of 2-deoxy-2[$^{18}$F]-fluoro-laminaribiose ([$^{18}$F]FDL):**

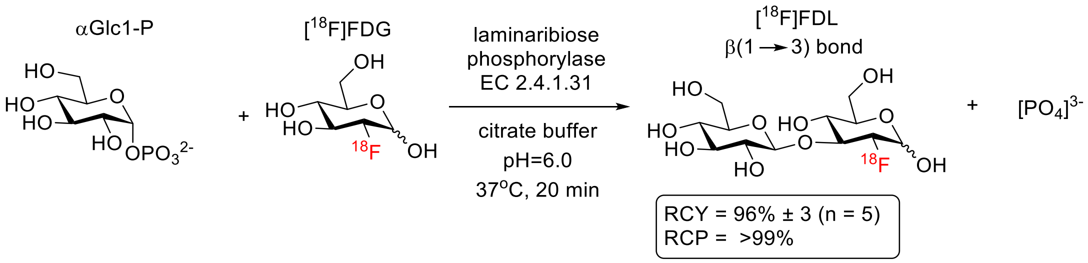

In a 4 mL borosilicate vial containing PTFE stir bar, laminaribiose phosphorylase (EC 2.4.1.31, Prof. Tom Desmet lab, Ghent University), (0.3 mg, 3 units) and αGlc1-P (6 mg, 0.020 mmol) were added. A dose of [$^{18}$F]FDG (10-15 mCi) in citrate buffer (0.1M, pH=6.0, 0.4-0.5 mL) was directly transferred to the vial and the mixture was stirred at 37 °C for 20 min. The mixture was diluted with MeCN then filtered through C18 light cartridge, before being purified via semi prep HPLC (YMC-Pack Polyamine II, 250 X 10 mm) using mobile phase 73% MeCN/27 % $H_2O$. [$^{18}$F]FDL was isolated in 5-7mL fractions. The fractions were then diluted with MeCN (40 mL) before being passed through Sep-pak Plus $NH_2$ Cartridge at 5 mL/min to trap each dimer product. After flushing the cartridge with air and $N_2$ gas, the tracer was eluted using saline solution for further analysis. [$^{18}$F]FDL (RCY= 96% ± 3%, RCP=99%), (N=5). Chemical purity of [$^{18}$F]FDL was confirmed by analytical HPLC.

**HPLC analysis of [$^{18}$F]FDL**

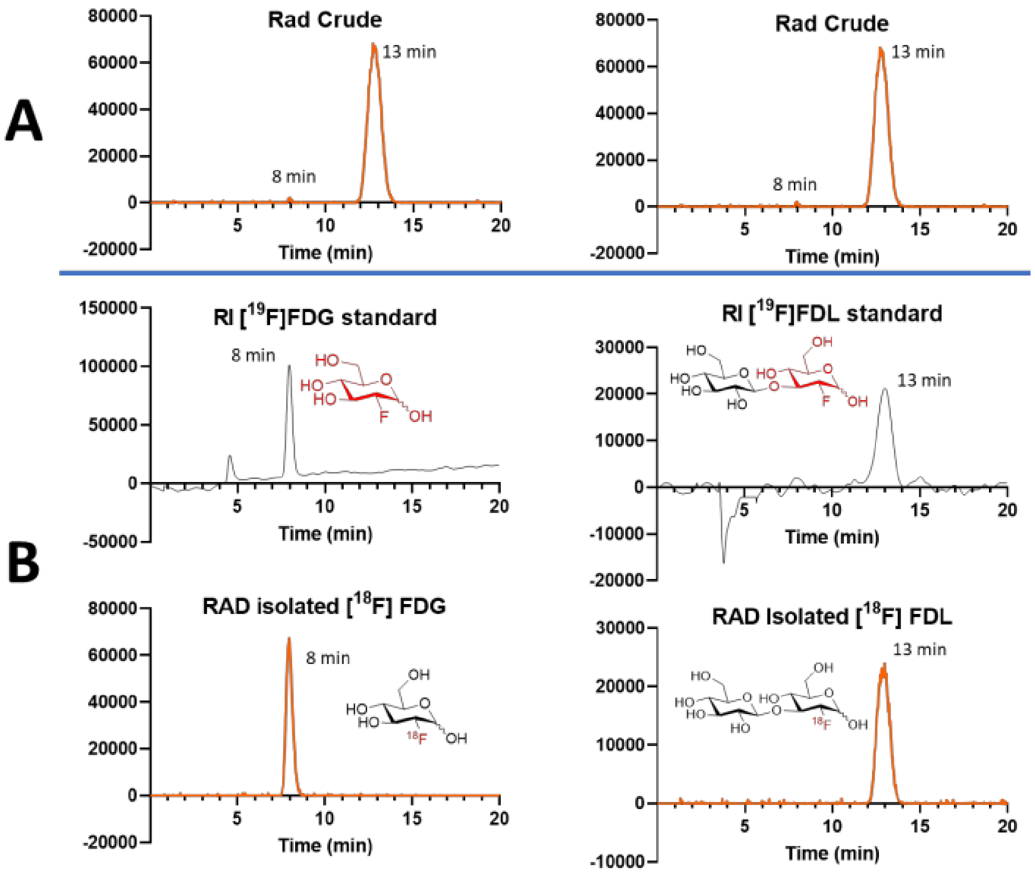

**Figure C.6.1.** HPLC analysis of 2-deoxy-2[$^{18}$F]-fluoro-laminaribiose ([$^{18}$F]FDL) with YMC-Pack Polyamine II, 250 X 4.6 mm using mobile phase 73% MeCN/27 % $H_2O$. A) Injection of crude, radioactivity (RAD) detection B) Co-injection of "cold" $^{19}$F standard and "hot" $^{18}$F isolated tracer, with both refractive index (RI) detection and radioactivity (RAD) detection.

**C.7. Radiosynthesis of 2-deoxy-2-[$^{18}$F]-fluoro-sakebiose ([$^{18}$F]FSK) with sakebiose phosphorylase:**

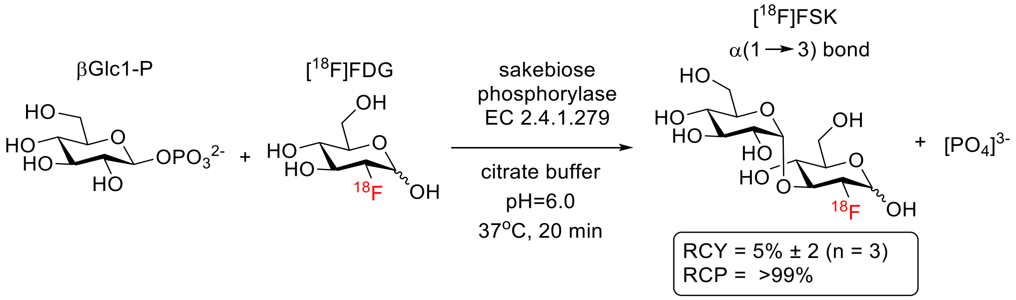

In a 4 mL borosilicate vial containing PTFE stir bar, sakebiose phosphorylase (EC 2.4.1.279, Creative Enzymes), (0.3 mg, 3 units) and βGlc1-P (6 mg, 0.020 mmol) were added. A dose of [$^{18}$F]FDG (10-15 mCi) in citrate buffer (0.1M, pH=6.0, 0.4-0.5 mL) was directly transferred to the vial and the mixture was stirred at 37 °C for 20 min. The mixture was diluted with MeCN then filtered through C18 light cartridge, before being purified via semi prep HPLC (YMC-Pack Polyamine II, 250 X 10 mm) using mobile phase 73% MeCN/27 % $H_2O$. [$^{18}$F]FSK was isolated in 5 mL fraction. The fraction was then diluted with MeCN (40 mL) before being passed through Sep-pak Plus $NH_2$ Cartridge at 5 mL/min to trap each dimer product. After flushing the cartridge with air and $N_2$ gas, the tracer was eluted using saline solution for further analysis. [$^{18}$F]FSK (RCY= 5% ± 2%, RCP=99%), (N=3). Chemical purity of [$^{18}$F]FSK was confirmed by analytical HPLC.

**HPLC analysis of [$^{18}$F]FSK**

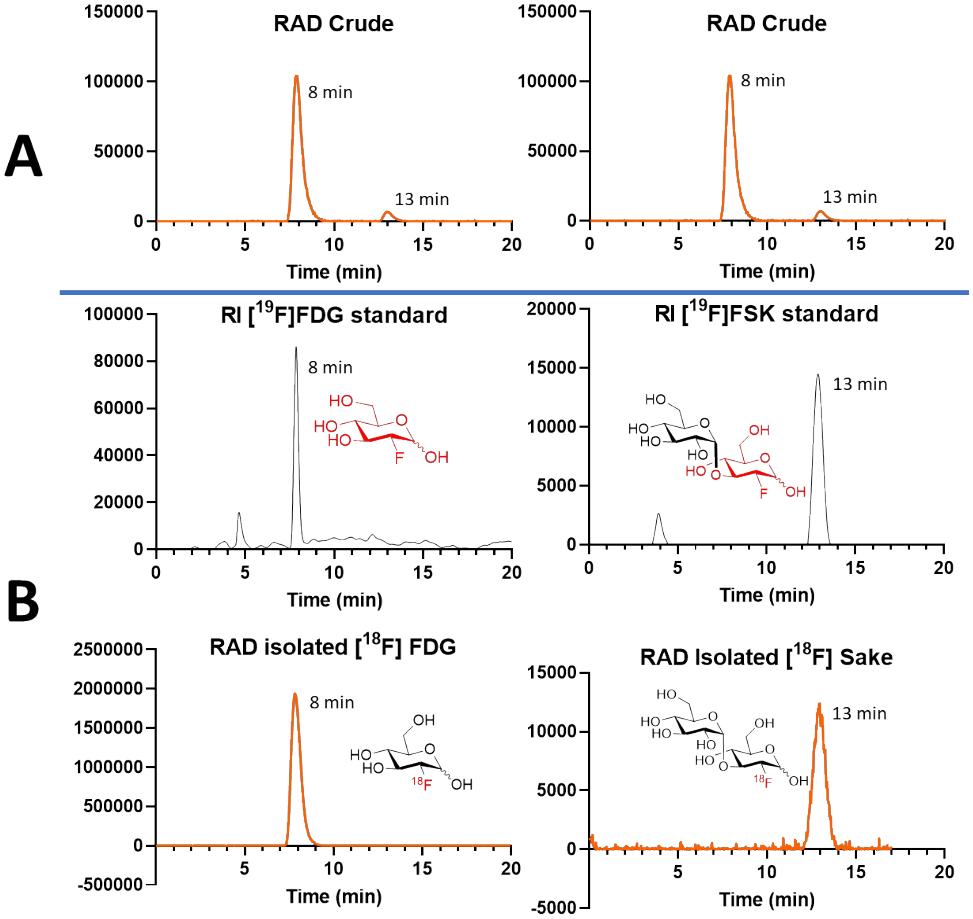

**Figure C.7.1.** HPLC analysis of 2-deoxy-2-[$^{18}$F]-fluoro-sakebiose ([$^{18}$F]FSK) with YMC-Pack Polyamine II, 250 X 4.6 mm using mobile phase 73% MeCN/27 % $H_2O$. A) Injection of crude, radioactivity (RAD) detection B) Co-injection of “cold” $^{19}$F standard and “hot” $^{18}$F isolated tracer, with both refractive index (RI) detection and radioactivity (RAD) detection.

### C.8. Control Experiment: Reaction without enzyme:

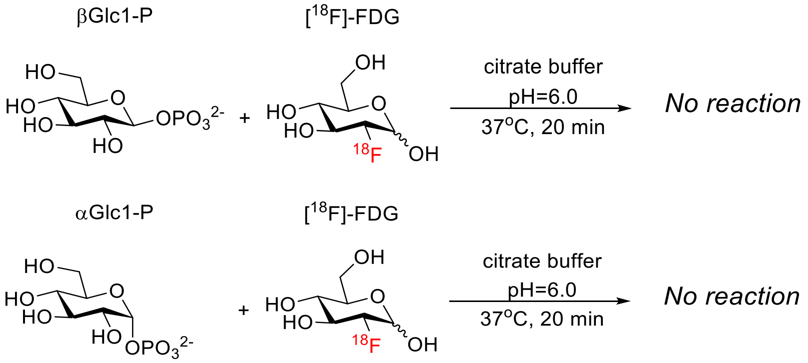

The reaction was conducted with the same conditions except without the presence of phosphorylase.

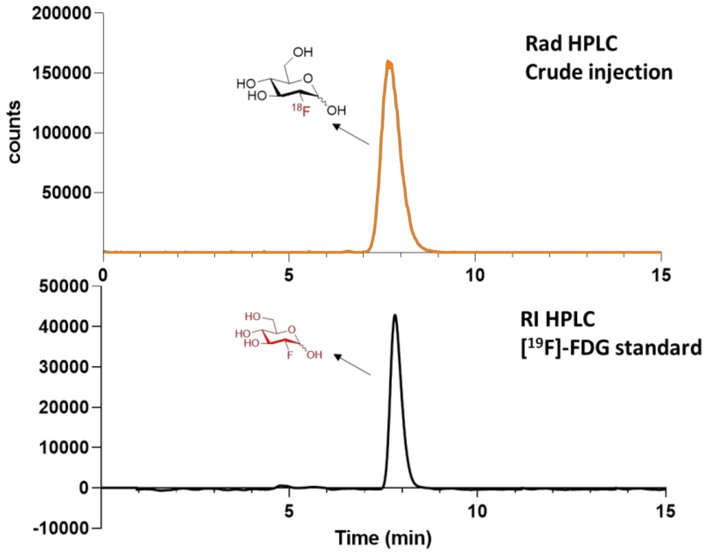

**Figure C.8.1.** HPLC analysis of control reaction: Rad HPLC only showed presence of unreacted [$^{18}$F]FDG when phosphorylase was omitted from reaction set up.
